# HaloTag9: an engineered protein tag to improve fluorophore performance

**DOI:** 10.1101/2021.04.01.438024

**Authors:** Michelle S. Frei, Miroslaw Tarnawski, Julia Roberti, Birgit Koch, Julien Hiblot, Kai Johnsson

**Affiliations:** Department of Chemical Biology, Max Planck Institute for Medical Research, Jahnstrasse 29, 69120 Heidelberg, Germany; Institute of Chemical Sciences and Engineering (ISIC), École Polytechnique Fédérale de Lausanne (EPFL), 1015 Lausanne, Switzerland; Protein Expression and Characterization Facility, Max Planck Institute for Medical Research, Jahnstrasse 29, 69120 Heidelberg, Germany; Leica Microsystems CMS GmbH, Am Friedensplatz 3, 68163 Mannheim, Germany

## Abstract

HaloTag9 is an engineered variant of HaloTag7 with up to 40% higher brightness and increased fluorescence lifetime when labeled with fluorogenic rhodamines. Moreover, combining HaloTag9 with HaloTag7 and other fluorescent probes enabled live-cell multiplexing using a single fluorophore and the generation of a fluorescence lifetime-based biosensor. The increased brightness of HaloTag9 and its use in fluorescence lifetime multiplexing makes it a powerful tool for live-cell imaging.

## Introduction

The need for brighter, more photostable, and spectrally diverse fluorophores for fluorescence microscopy has drawn the attention to small-molecule fluorophores, which exhibit superior photophysical properties than fluorescent proteins (FPs). Particularly, the combination of fluorogenic rhodamine-based fluorophores, which increase fluorescence upon target binding, with self-labeling protein (SLP) tags such as HaloTag7 has become a powerful tool for these purposes (Fig. 1A)^1–3^. The fluorogenicity of rhodamines is based on an environmentally sensitive equilibrium between a closed, non-fluorescent, spirocyclic and an open, fluorescent, quinoid form (Fig. 1B)^4^. While numerous strategies have been introduced to control the open-closed equilibrium of rhodamines via chemical modifications^4–7^, little attention has been paid to the influence of the protein, even though the fluorogenic turn-on is mostly determined by the change in environment^8–10^. Consequently, SLP engineering offers an alternative approach to control the fluorogenicity and the photophysical properties of fluorogenic rhodamines. Here we describe the engineering of HaloTag7 to increase the brightness and fluorescence lifetime of bound rhodamines for live-cell microscopy.

**Figure 1:**
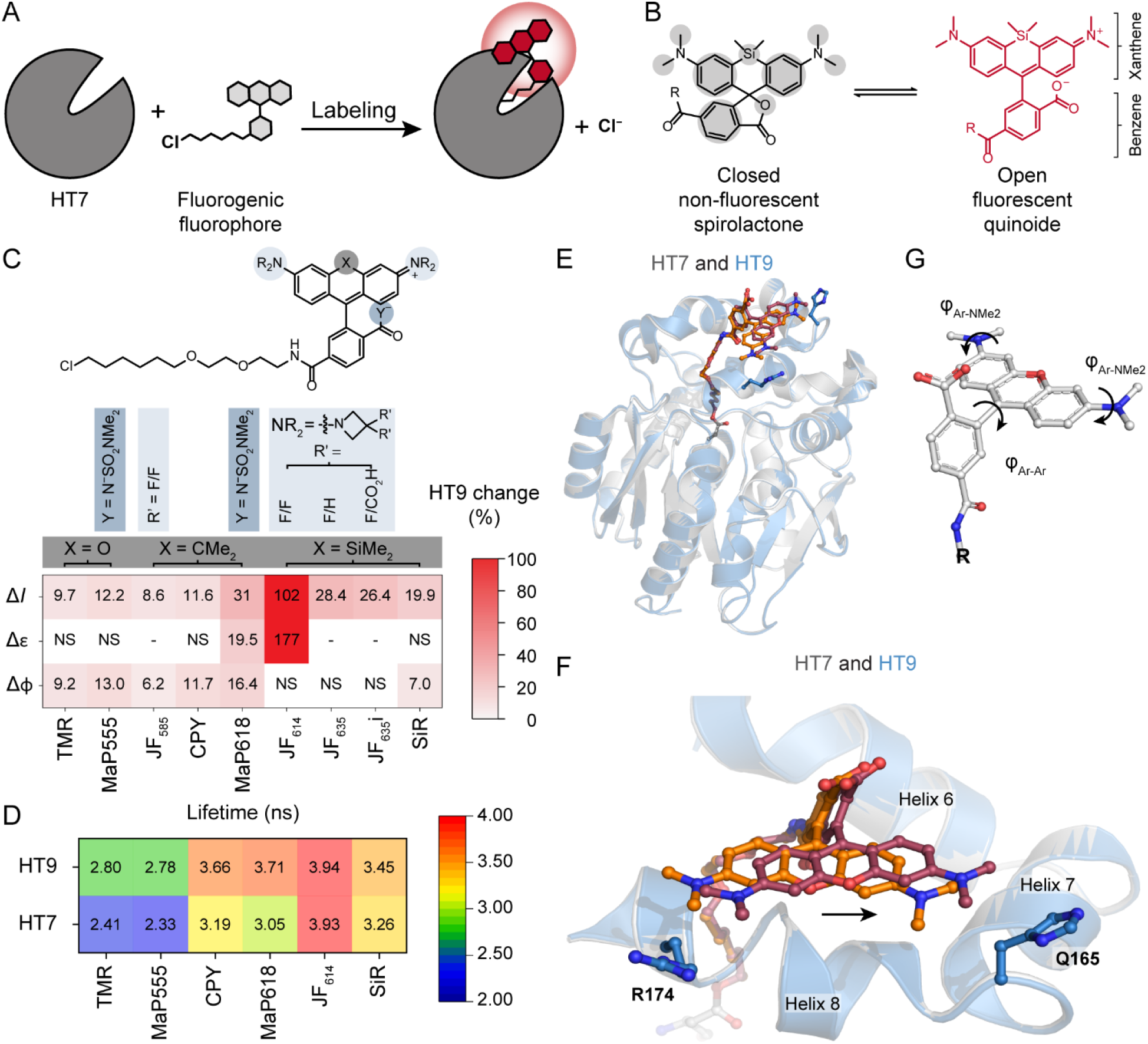
HaloTag9 characterization and comparison to HaloTag7. **A** Scheme of HaloTag7 labeling with a fluorogenic fluorophore, resulting in a fluorescence increase (turn-on) once the fluorophore is bound to the protein tag. **B** Environmentally sensitive open-closed equilibrium of SiR, a fluorogenic rhodamine fluorophore. The equilibrium position can be tuned through environmental changes or through chemical modifications indicated by the grey areas. R = chloroalkane (CA). **C** Relative *in vitro* fluorescence intensity (Δ*I*), extinction coefficient (Δε), and quantum yield (Δϕ) changes of labeled HaloTag9 compared to HaloTag7. Unless otherwise stated R = Me and Y^−^ = O^−^ in the generalized chemical structure (Supplementary Table S3; Δ*I*: mean, *N* = 2 replicates each 4 measurements; Δε: mean, *N* = 6 measurements; Δϕ: mean, *N* = 3 measurements; NS = not significant, - = not determined). **D** Fluorescence lifetimes (τ) of different rhodamines on HaloTag9 and HaloTag7 (mean, for *N* see Supplementary Table S8). **E** Structural comparison of HaloTag7-TMR (PDB ID: 6Y7A, 1.4 Å) and HaloTag9-TMR (PDB ID: 6ZVY, 1.4 Å, Chain A), which are represented as cartoon in grey or blue, respectively. H165 and R174 of HaloTag9 are represented as blue sticks. TMR is highlighted as orange (HaloTag7) and violet (HaloTag9) sticks. **F** Zoom onto the rhodamine binding site (helix 6-8). The arrow indicates the shift of the fluorophore, positioning the xanthene core closer toward residue 165 in HaloTag9 as compared to HaloTag7. **G** Dihedral angles *φ*_Ar-Ar_ and *φ*_Ar-NMe2_ of TMR.

## Results

To engineer the rhodamine-binding site of HaloTag7, ten amino acids in close proximity to tetramethyl-rhodamine (TMR) on the HaloTag7-TMR crystal structure (PDB ID: 6Y7A, 1.4 Å) were chosen for randomization by site-saturation mutagenesis. The resulting libraries were screened for increases in brightness upon labeling with the fluorogenic fluorophore silicon rhodamine (SiR-CA) (Fig. 1B, Supplementary Fig. S1). This led to the identification of the double mutant HaloTag9 (HaloTag7-Q165H-P174R), which increased the brightness of SiR by 19.9±0.5% compared to HaloTag7 (Supplementary Fig. S2, Supplementary Table S1-2). Characterization with a panel of 46 different fluorophores revealed that HaloTag9 increases the brightness of 14 rhodamines, including fluorophores based on the popular SiR, carbopyronine (CPY) and TMR scaffolds, which cover the spectral range between 550 and 650 nm (Fig. 1C, Supplementary Fig. S3, Supplementary Table S3-4). The most pronounced changes in fluorescence intensity (Δ*I*) for each scaffold were found for the fluorogenic rhodamines: JF_614_-CA^10^ (Δ*I* = 102±11%), MaP618-CA^6^ (Δ*I* = 31±4%), and MaP555-CA^6^ (Δ*I* = 12.2±1.7%), rendering these fluorophores even more fluorogenic. These increases are comparable to those achieved through chemical modifications of rhodamines, for example through the introduction of azetidines (e.g. 5-42%)^11^. In addition to rhodamines, increases in brightness were also observed for a fluorescein derivative and one cyanine (Cy3-CA; Supplementary Fig. S3C). All other tested fluorophores, most of which are non-fluorogenic, showed similar properties on HaloTag9 and HaloTag7. HaloTag9 and HaloTag7 displayed similar labeling kinetics and thermostability (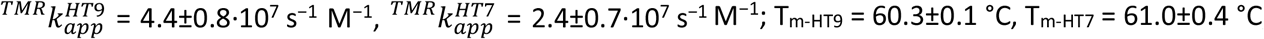; Supplementary Fig. S4, Supplementary Table S5).

The increases in brightness are due to changes in extinction coefficient (Δε) and/or quantum yield (Δϕ; Fig. 1C, Supplementary Table S6-7, 9). Fluorogenic fluorophores such as JF_614_-CA (Δ*I* = 102±11%) showed large increases in extinction coefficient (Δε = 177±17%), which were attributed to a shift in the open-closed equilibrium toward the open state when going from HaloTag7 to HaloTag9. In contrast, non-fluorogenic and fully open fluorophores like TMR-CA (Δ*I* = 9.7±0.6%) only showed changes in quantum yield (Δϕ = 9.2±0.7%). Generally, changes in quantum yield correlated with changes in fluorescence lifetime (Δτ) and were most pronounced for MaP618-CA (Δϕ = 16.4±0.4%, Δτ = 21.4±0.6%; Fig. 1D, Supplementary Table S6-9).

The molecular mechanisms underlying these changes were analyzed using the obtained HaloTag9-TMR crystal structure (PDB ID: 6ZVY, 1.4 Å; Fig. 1E-F, Supplementary Fig. S5-6; Supplementary Table S10-12). Structural alignment with HaloTag7-TMR (PDB ID: 6Y7A, 1.4 Å) highlighted the displacement of the xanthene core on HaloTag9 toward α-helix 7 (RMSD_xanth_ = 1.82±0.16 Å). This induces an increase in the rotational energy barriers around the aniline groups of TMR (*φ*_Ar-NMe2_) and a rise in the dihedral angle (*φ*_Ar-Ar_) between the xanthene and the benzene ring (*φ*_Ar-Ar-HT7_ = 119.0°, *φ*_Ar-Ar-HT9_ 122.32±0.21°; Fig. 1G, Supplementary Fig. S7-8). Both are favorable for brightness as non-radiative decay pathways via rotational motion or photoinduced electron transfer are suppressed^12,13^. This mechanism helps to rationalize the relatively small effects on quantum yield observed for azetidine-bearing rhodamines as non-radiative decay pathways via rotational motion are already reduced for them^11^. Additionally, the two mutations (Q165H-P174R) influence the local electrostatic potential around the fluorophore, going from predominantly negative on HaloTag7 to slightly positive on HaloTag9 (Supplementary Fig. S9). Such a change in the rhodamine’s solvation shell might stabilize its carboxylate group, favoring the open form and therefore increasing the extinction coefficient.

We then investigated the brightness of labeled HaloTag9 relative to HaloTag7 in living U-2 OS cells by confocal (Δ*I*_Ensemble_) and fluorescence correlation spectroscopy (FCS) measurements (Δ*I*_FCS_). Nine rhodamines, which were brighter *in vitro*, were tested *in cellulo* and all except one showed significantly higher brightness when bound to HaloTag9 than to HaloTag7 (Fig. 2A-B, Supplementary Fig. S10-11, Supplementary Table S13). The fluorophores with the largest improvements were MaP618-CA (Δ*I*_Ensemble_ = 39.2±0.9%, Δ*I*_FCS_ = 59±4%) and JF_614_-CA (Δ*I*_Ensemble_ = 28.0±1.1%, Δ*I*_FCS_ = 21±3%). The increased brightness is also evident when comparing live-cell images of the microtubule binding protein Cep41 labeled either with HaloTag7 or HaloTag9 (Fig 2C-D). HaloTag9 was further applicable to live-cell stimulated emission depletion (STED) microscopy and the fluorophores tested showed similar photostability as on HaloTag7 (Supplementary Fig. S12-14). Thus, HaloTag9 significantly increases the performance of fluorogenic fluorophores such as MaP618-CA in both live-cell confocal and STED microscopy and it is also expected to do so in other contexts such as biosensing (*vide infra*) or *in vivo* microscopy^7,14^.

**Figure 2:**
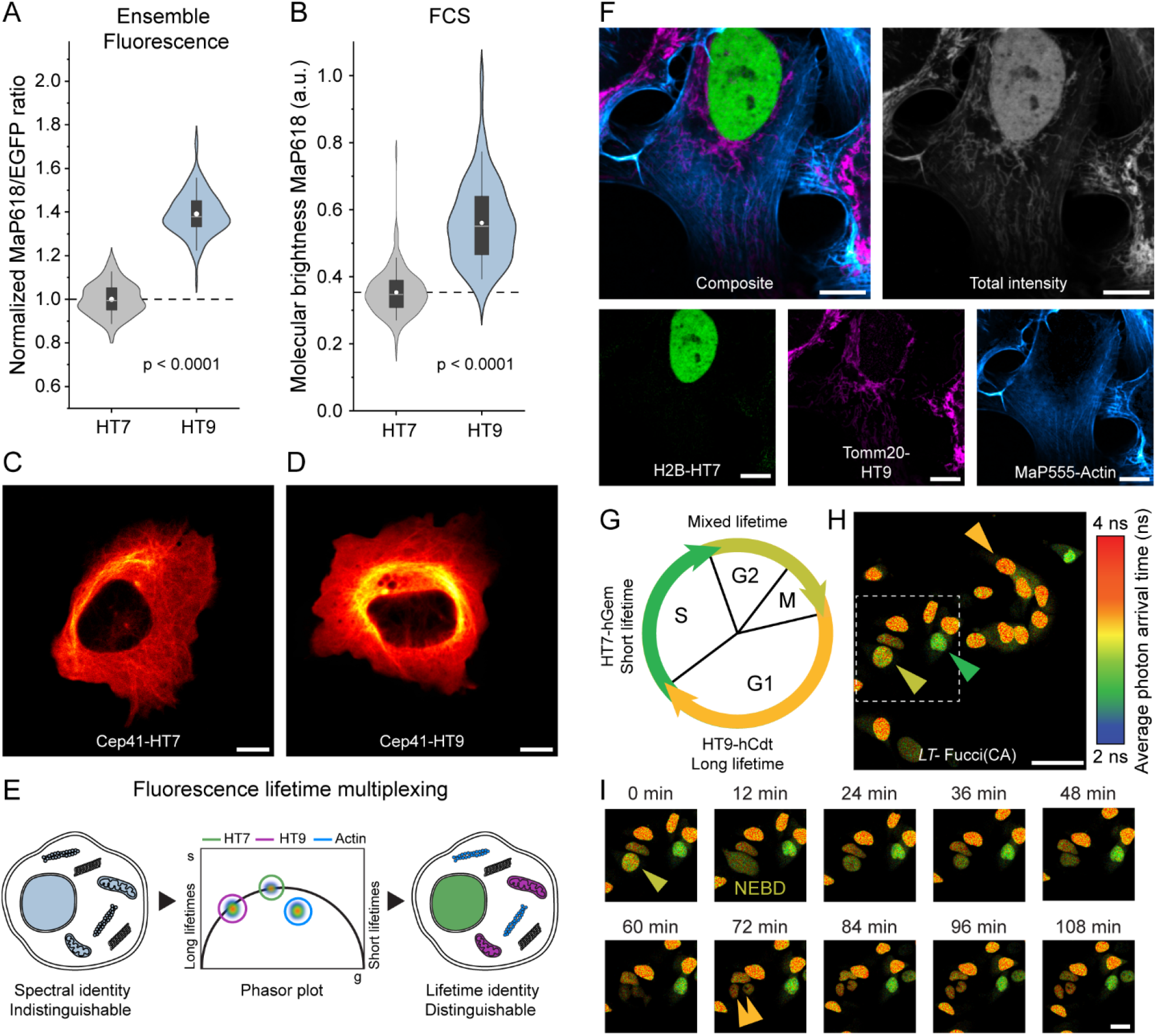
Application of HaloTag9 in live-cell microscopy and biosensing. **A-B** *In cellulo* brightness measurements comparing HaloTag7 and HaloTag9 (U-2 OS cells cytosol) labeled with MaP618-CA. Fluorescence intensities were measured by confocal microscopy (**A**) or FCS (**B**). Both methods demonstrate the superior performance of HaloTag9 compared to HaloTag7 (Δ*I*_Ensemble_ = 39.2±0.9%, Δ*I*_FCS_ = 59±4%; *N* = 120 and 119 cells, *N* = 155 and 170 traces from 36 cells, from three independent experiments, Supplementary Table S13). Distribution = light grey or blue, box = 25%–75% percentile, whiskers = 5%– 95% percentile, white line = median, circle = mean, dashed line = mean of HaloTag7 for comparison. p-Values are given based one-sided t-tests (α = 5%, r = −0.02, 0.27, DF = 237 and 323). **C-D** Confocal fluorescence microscopy images of microtubules labeled with HaloTag7 (**C**) or HaloTag9 (**D**). U-2 OS cells expressing the microtubule marker Cep41 as a fusion of HaloTag7 or HaloTag9 labeled with MaP618-CA (1 μM, 3 h). The brightness of the images was scaled to the expression level of the two proteins in the two cells. Scale bars, 10 μm. **E** Schematic view of fluorescence lifetime multiplexing using only one rhodamine for three targets (nucleus, mitochondria, and F-actin). Their spectral identity does not allow them to be distinguished but they can be separated using fluorescence lifetime information via phasor analysis. **F** Fluorescence lifetime multiplexing of U-2 OS cells expressing histone H2B as a HaloTag7 fusion and the outer-mitochondrial membrane protein Tomm20 as a HaloTag9 fusion. Both proteins were labeled with MaP555-CA (1 μM, 3 h). In addition, MaP555-Actin was used to label F-Actin (2 μM, 3 h). The composite, the total intensity and the three individual images with the separated structures are given. Scale bars, 10 μm. **G** Schematic overview of the *LT*-Fucci(CA) biosensor. During the G1 phase mainly HaloTag9-hCdt is present and the nuclei will therefore present long average photon arrival times (3.7 ns/orange). During S phase HaloTag7-hGem is predominant, resulting in shorter average photon arrival times (3.1 ns/green) and during G2 and M phase a mixture of both will be present (∼3.4 ns/light-green). **H** Representative FastFLIM image of U-2 OS cells stably expressing the *LT*-Fucci(CA) biosensor labeled with MaP618-CA (1 μM). Cells in three different cell stages can be found (orange arrowhead: G1, green arrowhead: S, light-green arrowhead: G2 and M). Scale bar, 50 μm. **I** Zoom-in from panel (**H**) showing the division of a cell over time. The dividing cell (light-green arrowhead), the two daughter cells (orange arrowhead) and the moment of nuclear envelope breakdown (NEBD) are indicated (Supplementary video 1). Scale bar, 25 μm.

Beyond improving fluorophore brightness, the change in fluorescence lifetime going from HaloTag7 (τ_MaP618_ = 3.1 ns) to HaloTag9 (τ_MaP618_ = 3.7 ns) creates the opportunity to use both tags simultaneously for fluorescence lifetime multiplexing using a single fluorophore (Fig. 2E)^15^. HaloTag7 and HaloTag9 were thus co-expressed as fusion proteins of the histone protein H2B and the outer mitochondrial membrane protein Tomm20 in U-2 OS cells. They were then labeled with a single fluorophore and the two tags distinguished by fluorescence lifetime imaging microscopy (FLIM) using phasor analysis^16^. We performed fluorescence lifetime multiplexing in two different spectral windows using either MaP555-CA or MaP618-CA and with different combinations of cellular targets either in fixed or living cells (Supplementary Fig. S15-18). Furthermore, pairing the two HaloTags with probes such as MaP555- or MaP618-Actin, -Tubulin or -DNA^6^, allowed to multiplex even three species in living cells within the same spectral channel (Fig. 2F, Supplementary Fig. S19-20). Additionally, super-resolved images were acquired by STED-FLIM using only one depletion laser, while separating two species based on fluorescence lifetime information (Supplementary Fig. S21). To the best of our knowledge, this is the first time multi-target FLIM was achieved using a single fluorophore on a subcellular level in living cells, highlighting the potential of the combination of HaloTag9 and HaloTag7 for live-cell fluorescence lifetime multiplexing.

Subsequently, we exploited the difference in fluorescence lifetime of the two HaloTags for the generation of a fluorescence lifetime-based biosensor to monitor cell cycle progression. Based on Fucci biosensors, which rely on the cell cycle-dependent degradation of hCdt and Geminin (hGem) fragments fused to a green and red FP^17^, lifetime-Fucci(CA) (*LT*-Fucci(CA)) was developed, replacing the two FPs with HaloTag7 and HaloTag9. Specifically, the cell cycle stage of U-2 OS cells stably expressing *LT*-Fucci(CA) was clearly indicated by the average photon arrival time of the nuclei upon labeling with MaP618-CA and could be followed over 24 h when providing the fluorogenic fluorophore continuously (3.7 ns-orange-G1; 3.1 ns-green-S; ∼3.4 ns-light-green-G2/M; Fig. 2G-I, Supplementary video 1). Phasor analysis of cells in the G2/M phase, in which both hCdt and hGem are present, allowed to attribute relative amounts of the two proteins. Due to the large fluorescence lifetime changes of *LT*-Fucci(CA)-MaP618, TauContrast imaging, a confocal technique without the need for FLIM instrumentation, could be used to assess the cell cycle stage^18^. Additionally, the biosensor’s color could be simply switched by labeling with MaP555-CA and two *LT*-Fucci biosensors with alternative degradation cycles were generated (Supplementary Fig. S22-23, Supplementary videos 2-3). As *LT*-Fucci biosensors only occupy one variable spectral channel, they are ideally suited for combination with other biosensors or probes. *LT*-Fucci(CA)-MaP618, for instance, can be combined with the green spectral region and, due to its narrow emission spectrum, also with the far-red region (Supplementary Fig. S23)^6,19^. We thus simultaneously performed FLIM measurements of *LT*-Fucci(CA)-MaP618 and the RhoA GTPase activity biosensor Raichu-RhoA-CR (Clover-mRuby2) during cell division (Supplementary Fig. S24)^20^. The flexibility to choose the biosensor’s color at the labeling step and the improved multiplexing capabilities sets *LT*-Fucci biosensors apart from the recently published FUCCI-Red (mKate2-mCherry)^19^.

In summary, HaloTag9 increases the brightness and fluorescence lifetime of fluorogenic rhodamines relative to HaloTag7. This improves its performance in fluorescence microscopy and opens up new possibilities for fluorescence lifetime multiplexing and the generation of fluorescence lifetime-based biosensors.

## Data availability

Plasmids encoding for HaloTag9 and fusions thereof have been deposited on Addgene. Accession codes can be found in Supplementary Table S13. The X-ray crystal structure of HaloTag9-TMR has been deposited to the PDB with deposition code 6ZVY. Correspondence and requests for materials should be addressed to K.J.

## Acknowledgements

This work was supported by the Max Planck Society and the École Polytechnique Fédérale de Lausanne. M.S.F was supported by the Deutsche Forschungsgemeinschaft (DFG, German Research Foundation) SFB TRR 186.

The authors thank I. Schlichting for X-ray data collection. Diffraction data were collected at the Swiss Light Source, beamline X10SA, of the Paul Scherrer Institute, Villigen, Switzerland. The authors thank A. Bergner, B. Mathes, D. Schmidt, M. Huppertz, and C. Gondrand, for provision of reagents. We acknowledge L. Lavis and J. Grimm for the provision of JF fluorophores. We thank J. Reinstein and J. Wilhelm for help with the stop flow measurements, E. d’Este for help with STED microscopy and F. Schneider for help with FCS measurements.

## Author contributions

M.S.F. performed the *in vitro* screen; produced, characterized and applied all HaloTag variants; performed computational studies; performed confocal, FCS, FLIM, and STED microscopy and the analysis thereof. M.T. solved the crystal structure of HaloTag9. M.S.F. and J.H. analyzed the crystal structure. J.H. and M.T. performed thermostability measurements. M.S.F. and B.K. generated stable cell lines. J.R. performed species separation based on phasor analysis, STED-FLIM, and TauContrast imaging. K.J. supervised the work. M.S.F., J.H. and K.J. wrote the manuscript with input from all authors.

## References

1. Xue, L., Karpenko, I. A., Hiblot, J. & Johnsson, K. Imaging and manipulating proteins in live cells through covalent labeling. Nat. Chem. Biol. 11, 917–923 (2015).

2. Péresse, T. & Gautier, A. Next-Generation Fluorogen-Based Reporters and Biosensors for Advanced Bioimaging. Int. J. Mol. Sci. 20, 6142 (2019).

3. Los, G. V et al. HaloTag: A Novel Protein Labeling Technology for Cell Imaging and Protein Analysis. ACS Chem. Biol. 3, 373–382 (2008).

4. Lavis, L. D. Teaching Old Dyes New Tricks: Biological Probes Built from Fluoresceins and Rhodamines. Annu. Rev. Biochem. 86, 825–843 (2017).

5. Zheng, Q. et al. Rational Design of Fluorogenic and Spontaneously Blinking Labels for Super-Resolution Imaging. ACS Cent. Sci. 5, 1602–1613 (2019).

6. Wang, L. et al. A general strategy to develop cell permeable and fluorogenic probes for multicolour nanoscopy. Nat. Chem. 12, 165–172 (2020).

7. Grimm, J. B. et al. A general method to optimize and functionalize red-shifted rhodamine dyes. Nat. Methods 17, 815–821 (2020).

8. Presman, D. M. et al. Quantifying transcription factor binding dynamics at the single-molecule level in live cells. Methods 123, 76–88 (2017).

9. Erdmann, R. S. et al. Labeling Strategies Matter for Super-Resolution Microscopy: A Comparison between HaloTags and SNAP-tags. Cell Chem. Biol. 26, 584–592 (2019).

10. Deo, C. et al. The HaloTag as a general scaffold for far-red tunable chemigenetic indicators. Nat. Chem. Biol. (2021). doi:10.1038/s41589-021-00775-w

11. Grimm, J. B. et al. A general method to improve fluorophores for live-cell and single-molecule microscopy. Nat. Methods 12, 244–250 (2015).

12. Savarese, M. et al. Fluorescence lifetimes and quantum yields of rhodamine derivatives: New insights from theory and experiment. J. Phys. Chem. A 116, 7491–7497 (2012).

13. Savarese, M. et al. Non-radiative decay paths in rhodamines: New theoretical insights. Phys. Chem. Chem. Phys. 16, 20681–20688 (2014).

14. Grimm, J. B. et al. A general method to fine-tune fluorophores for live-cell and in vivo imaging. Nat. Methods 14, 987–994 (2017).

15. Niehörster, T. et al. Multi-target spectrally resolved fluorescence lifetime imaging microscopy. Nat. Methods 13, 257–262 (2016).

16. Digman, M. A., Caiolfa, V. R., Zamai, M. & Gratton, E. The phasor approach to fluorescence lifetime imaging analysis. Biophys. J. 94, L14–L16 (2008).

17. Sakaue-Sawano, A. et al. Genetically Encoded Tools for Optical Dissection of the Mammalian Cell Cycle. Mol. Cell 68, 626–640 (2017).

18. Roberti, M. J. et al. TauSense: a fluorescence lifetime-based tool set for everyday imaging. Nat. Methods (2020).

19. Shirmanova, M. V et al. FUCCI - Red: a single - color cell cycle indicator for fluorescence lifetime imaging. Cell. Mol. Life Sci. (2021). doi:10.1007/s00018-020-03712-7

20. Lam, A. J. et al. Improving FRET dynamic range with bright green and red fluorescent proteins. Nat. Methods 9, 1005–1012 (2012).

