## Supplementary Information for "HaloTag9: an engineered protein tag to improve fluorophore performance"

Note: This is a revised version of the manuscript "HaloTag8: an engineered protein tag to improve fluorophore performance". In here the name of the generated protein tag was altered to HaloTag9 in order to avoid any confusion with another tool published under the name HaloTag8[1].

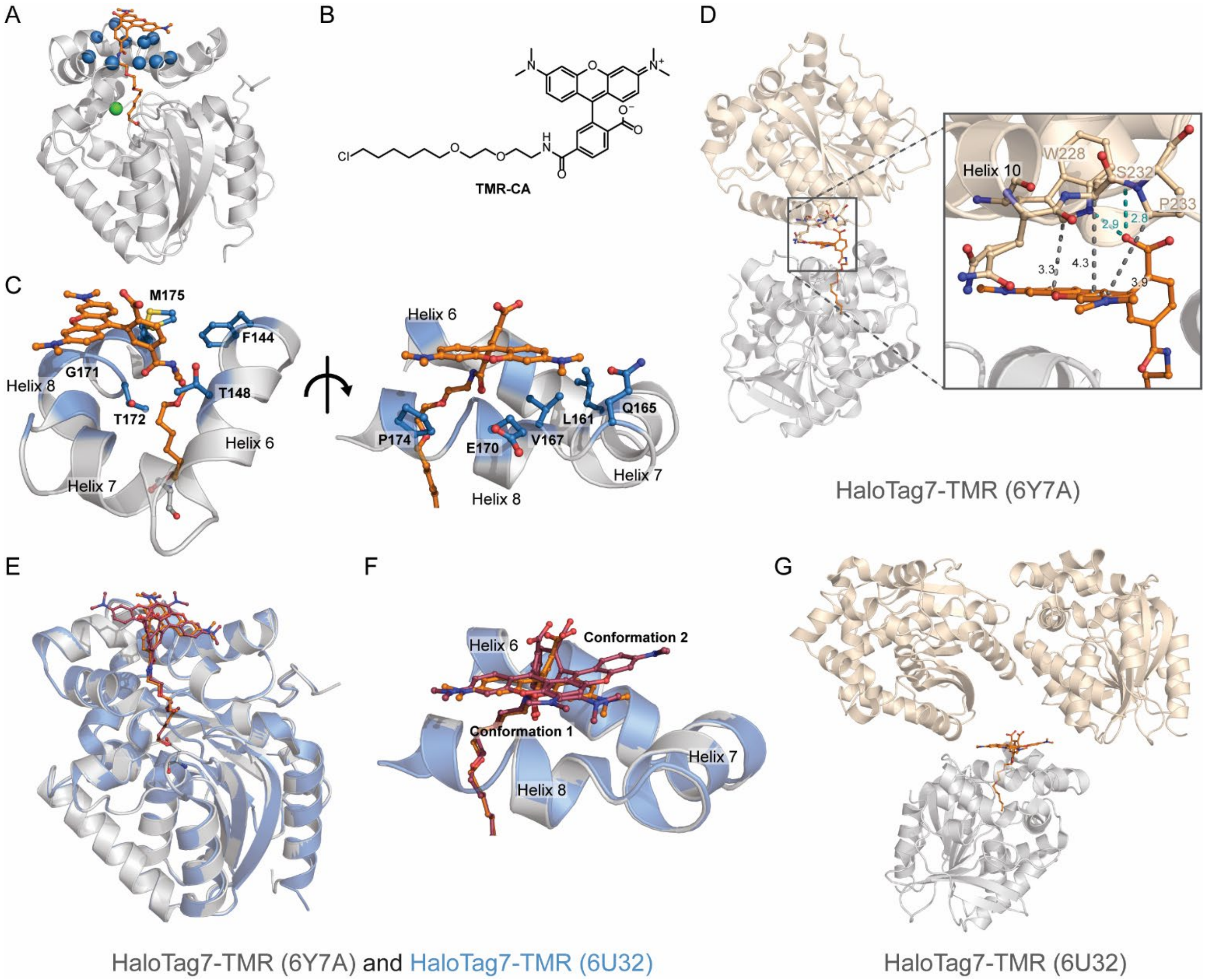

**Supplementary Figure S1:** Library design based on the structure of HaloTag7-TMR. **A** Crystal structure of HaloTag7-TMR (PDB ID: 6Y7A, 1.4 Å). The protein is represented as grey cartoon and TMR as orange sticks. The Cα of the ten amino acids chosen for site-saturation mutagenesis are highlighted as blue spheres. The chlorine atom is shown as a green sphere. **B** Chemical structure of TMR-CA. **C** TMR binding site (helices 6-8) on the HaloTag7-TMR crystal structure. Same structural representation, despite that the ten amino acids' side chains are represented by blue sticks. **D** Interactions between two HaloTag7-TMR monomers within the crystal packing. The carboxylic acid of TMR is hydrogen bonded (teal dashes) to the next monomer via S232 and W228 (cream). In addition, the distances from W228 (Cδ), S232 (Cα), and P233 (Cδ) to the xanthene ring system (grey dashes) are given in Ångström. Only W228 comes close enough to the xanthene system to be involved in efficient van der Waals interactions. **E-F** Overlay (**E**) and zoom (**F**) of HaloTag7 (grey) and an alternative crystal structure of HaloTag7-TMR (PDB ID: 6U32, 1.8 Å)[2]. TMR is given in stick representation (orange (6Y7A) and violet (6U32)). The 6U32 structure shows two fluorophore conformations: one almost identical to the 6Y7A one (conformation 1), the second (conformation 2) places the xanthene core over the turn between helix 7 and helix 8. **G** Crystal packing of the 6U32 HaloTag7-TMR structure. The fluorophore is not constrained by neighboring monomers, allowing TMR to adopt conformation 2.

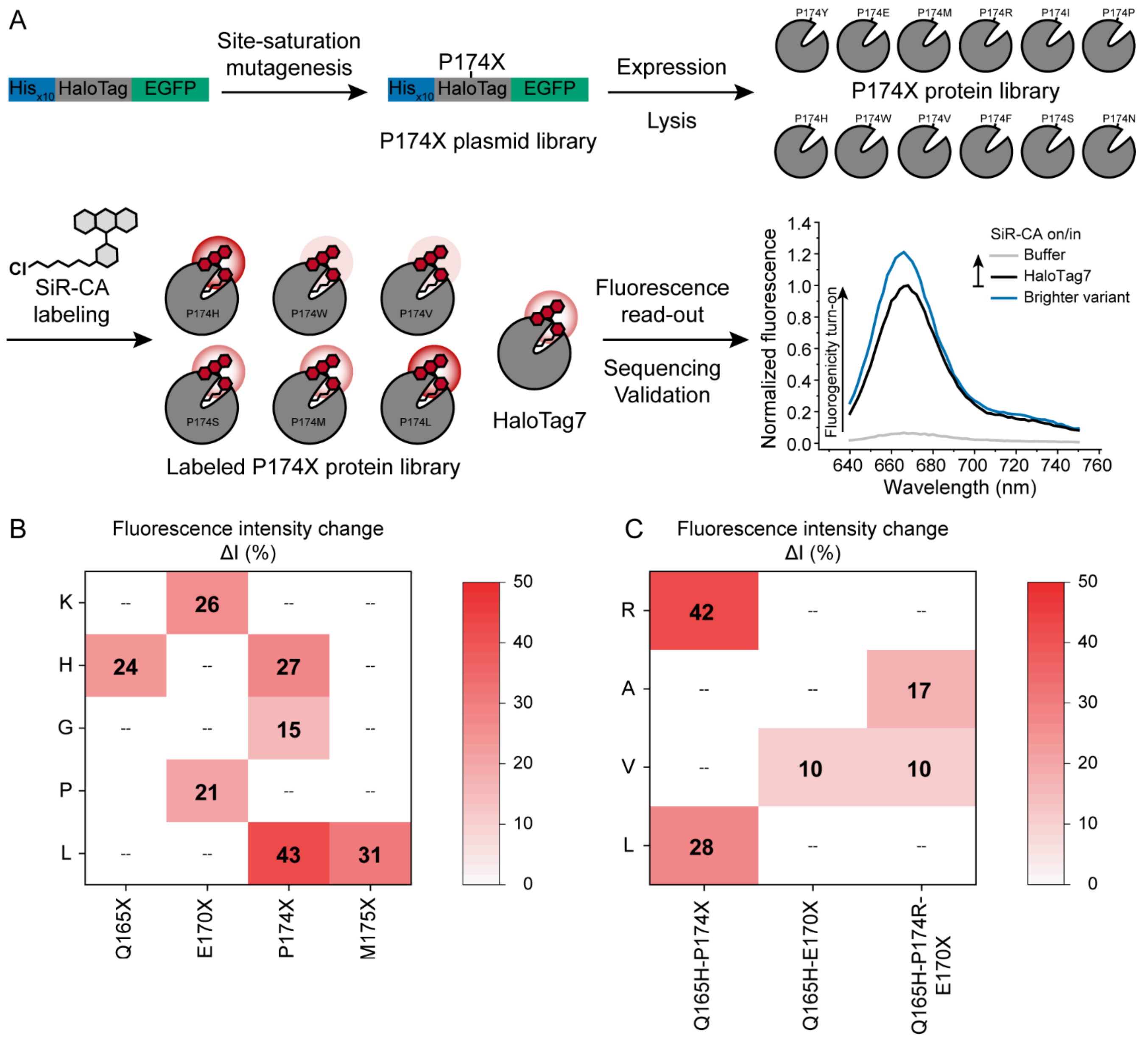

**Supplementary Figure S2:** HaloTag7 engineering strategy and results. **A** Schematic representation of the *in vitro* engineering of HaloTag7. Site saturation mutagenesis was performed onto the plasmid containing His$_{x10}$-HaloTag7-EGFP using degenerate primers and Gibson cloning. The plasmid library (e.g. P174X) was transformed in *E. coli* for protein production and extraction. The cell lysate was labeled with a limiting amount of SiR-CA and screened for increases in fluorescence intensity compared to HaloTag7. Brighter hits (increased fluorescence intensity of SiR on variants in comparison to SiR on HaloTag7) were sequenced and validated in a separate fluorescence assay using purified protein. For visibility EGFP is not shown in the expressed and labeled protein libraries. **B** Outcome of the first round of screening. Validated hits are given with their fluorescence intensity change ($\Delta I_{var\text{-}EGFP} = (I_{var} - I_{HT7}) \cdot I_{HT7}^{-1}$, mean, $N$ = 3 samples). **C** Outcome of the second and third round of screening. Validated hits are given with their fluorescence intensity change ($\Delta I_{var\text{-}EGFP}$, mean, $N$ = 3 samples).

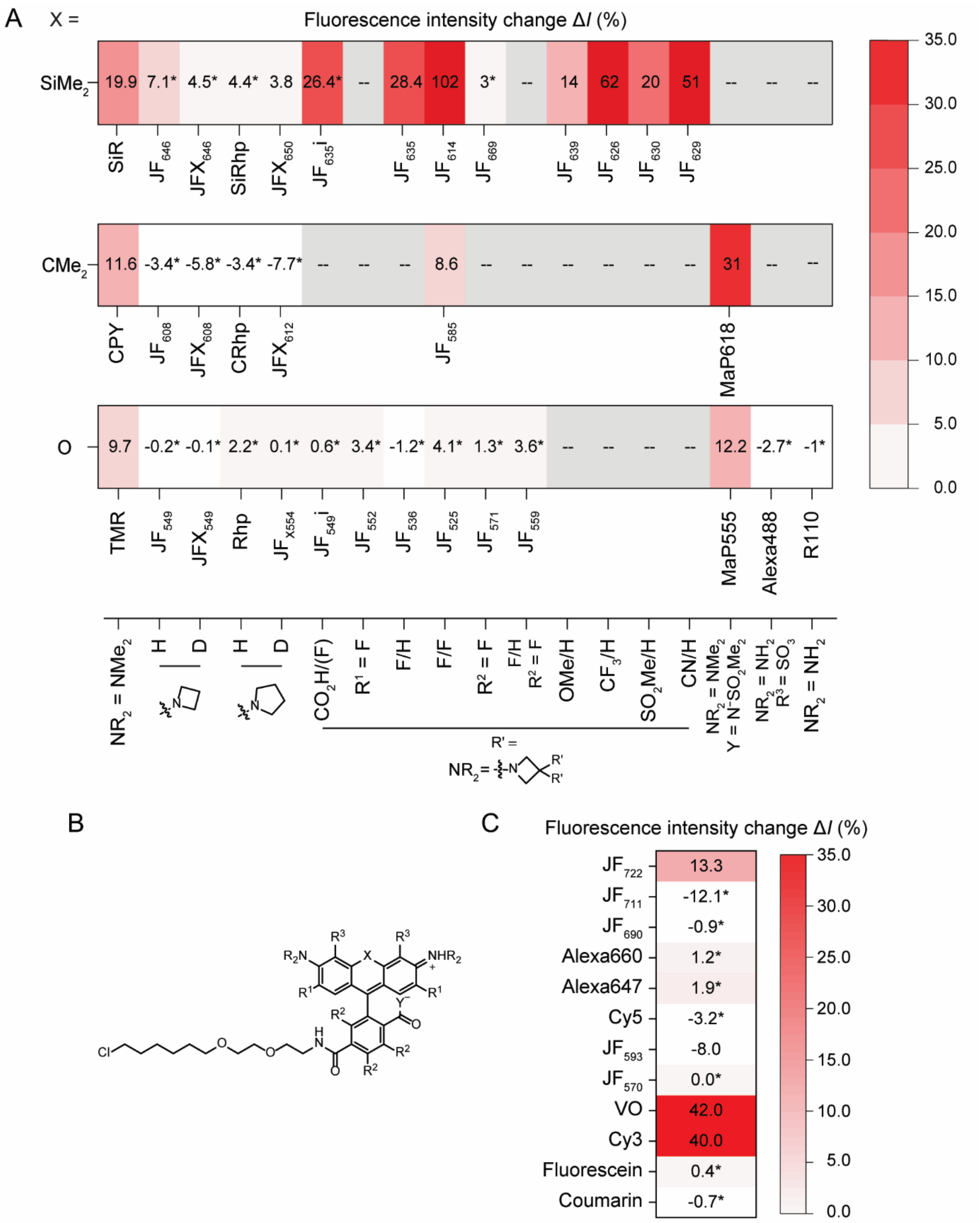

**Supplementary Figure S3:** Comparison of the fluorescence brightness between HaloTag7 and HaloTag9. **A** Fluorescence intensity change ($\Delta I = (I_{HT9} - I_{HT7}) \cdot I_{HT7}^{-1}$) of rhodamine based fluorophores reacted with HaloTag9 compared to HaloTag7. Fluorophore structures can be found in Supplementary Table S3 or can be inferred using the legend and the generic rhodamine structure in (**B**). Unless otherwise stated $R^1$, $R^2$, and $R^3$ = H, $Y^-$ = $O^-$, and R' = H. **B** Chemical structure of generic rhodamine. **C** Fluorescence intensity changes for non-traditional rhodamines (e.g. X = S ($JF_{570}$ and $JF_{593}$) or X = P=O(OH)/P=OPh ($JF_{690}$, $JF_{711}$, and $JF_{722}$)) as well as fluoresceins (fluorescein and Virginia Orange (VO)), cyanines (Cyanine3, Cyanine5, and Alexa647), and coumarin when reacted with HaloTag9 in comparison to HaloTag7. Fluorophore structures can be found in Supplementary Table S3. Mean, $N$ = 2 replicates each 4 samples. * Not significant. – Fluorophore not measured.

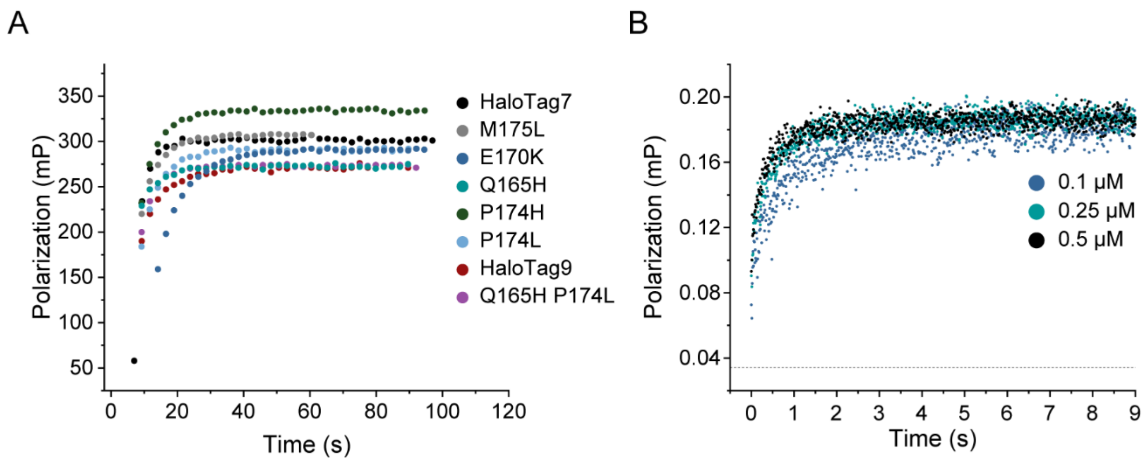

**Supplementary Figure S4:** Labeling kinetics of HaloTag7 variants with TMR-CA. **A** Plots of labeling kinetics of seven HaloTag7 variants compared to HaloTag7. Kinetic measurements were performed by fluorescence polarization on a plate reader. Proteins (80 nM) were reacted with limiting amounts of TMR-CA (20 nM) to allow complete labeling of fluorophores. The data was fitted with a mono-exponential function and the apparent first order rate constants $k_{1app}$ and the polarization values reached after reaction completion were compared (Supplementary Table S5). Only HaloTag7-E170K showed slower labeling kinetics than HaloTag7. Representative measurements of three replicates are displayed. **B** Labeling kinetics of HaloTag9. Kinetic measurements were performed by stop flow measurements. HaloTag9 was reacted with an equimolar amount of TMR-CA and the change in fluorescence polarization was followed. The measurement was performed with three different concentrations (0.1, 0.25 and 0.5 µM). The data was globally fit to a reaction model taking an initial association equilibrium (P + S ⬄ PS*) followed by an irreversible labeling reaction (PS* -> PS) into account. From this $K_D$, and $k_{app}$ were calculated (Supplementary Table S5). Similar measurements were performed for HaloTag7. Averaged measurements from at least 3 measurements are shown. The dashed line at 0.034 mP corresponds to the lower bound measured by a sample of free TMR-CA in solution.

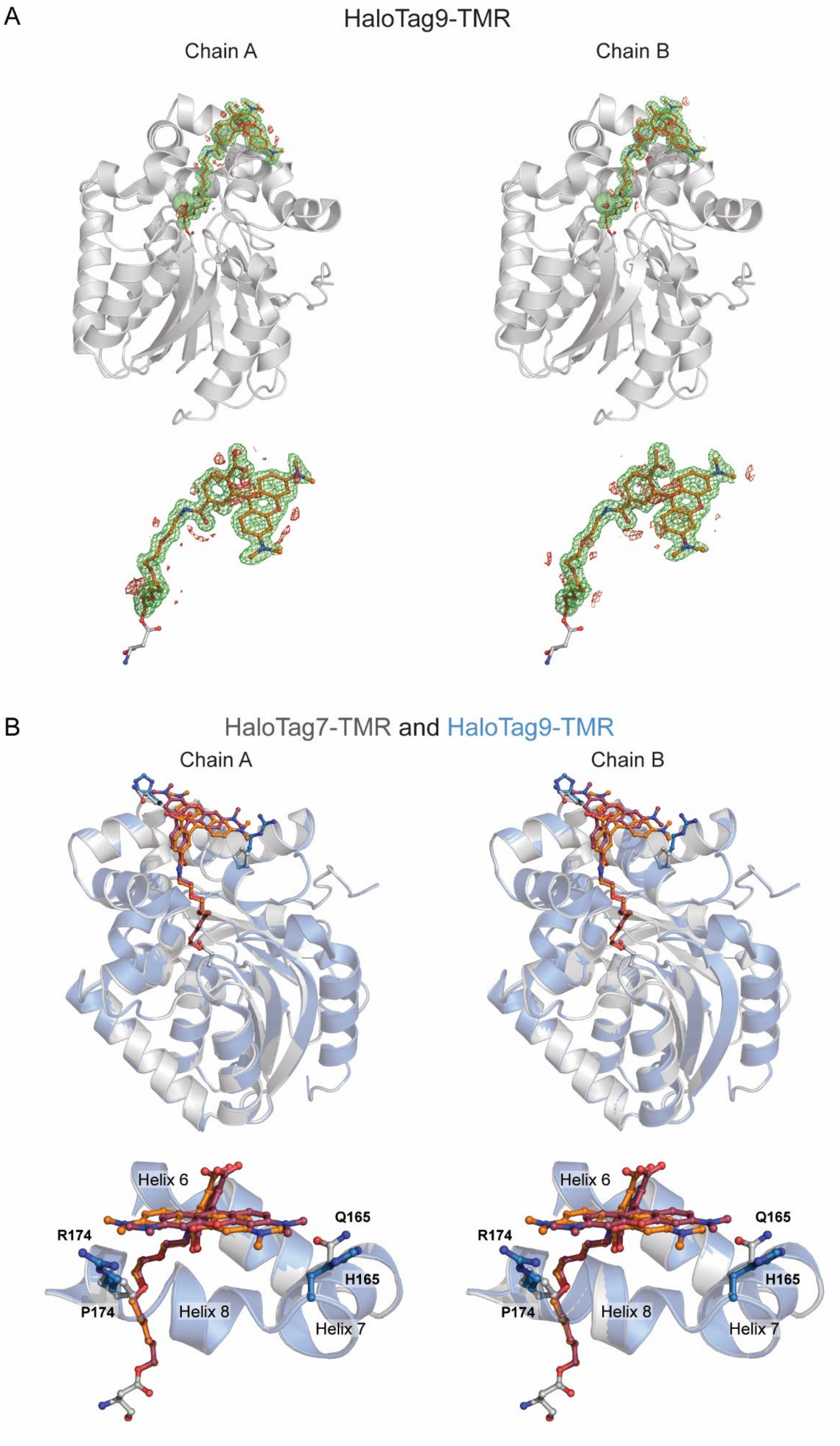
A
HaloTag9-TMR
Chain A
Chain B
B
HaloTag7-TMR and HaloTag9-TMR
Chain A
Chain B
Helix 6
Q165
R174
H165
P174
Helix 8
Helix 7
Helix 6
Q165
R174
H165
P174
Helix 8
Helix 7

**Supplementary Figure S5:** Crystal structure of HaloTag9-TMR. **A** OMIT-map of TMR from the HaloTag9-TMR structure (PDB ID: 6ZVY, 1.4 Å, Supplementary Table S12). The protein of each chain is represented as a cartoon (grey) while TMR is represented as sticks (orange). The chlorine atom is represented as a green sphere. The OMIT electron-density map for TMR is contoured at ±3.0 σ and represented as green (positive) and red (negative) mesh. In addition, a zoom onto the TMR ligand is given. **B** Structural comparison of HaloTag9-TMR (PDB ID: 6ZVY, 1.4 Å) with HaloTag7-TMR (PDB ID: 6Y7A, 1.4 Å, only chain A). The proteins are given in cartoon representation: HaloTag7 in grey and HaloTag9 in blue. The bound TMR is given in stick representation in orange (HaloTag7) and violet (HaloTag9). In addition, a zoom onto helices 6–8 with the proteins in cartoon representation and the TMR ligand as well as the mutated amino-acids as sticks is given. The two proteins show very similar protein structures ($RMSD_{\alpha C}$ = 0.204±0.015 Å; mean±s.d., *N* = 2 monomers; Supplementary Table S11).

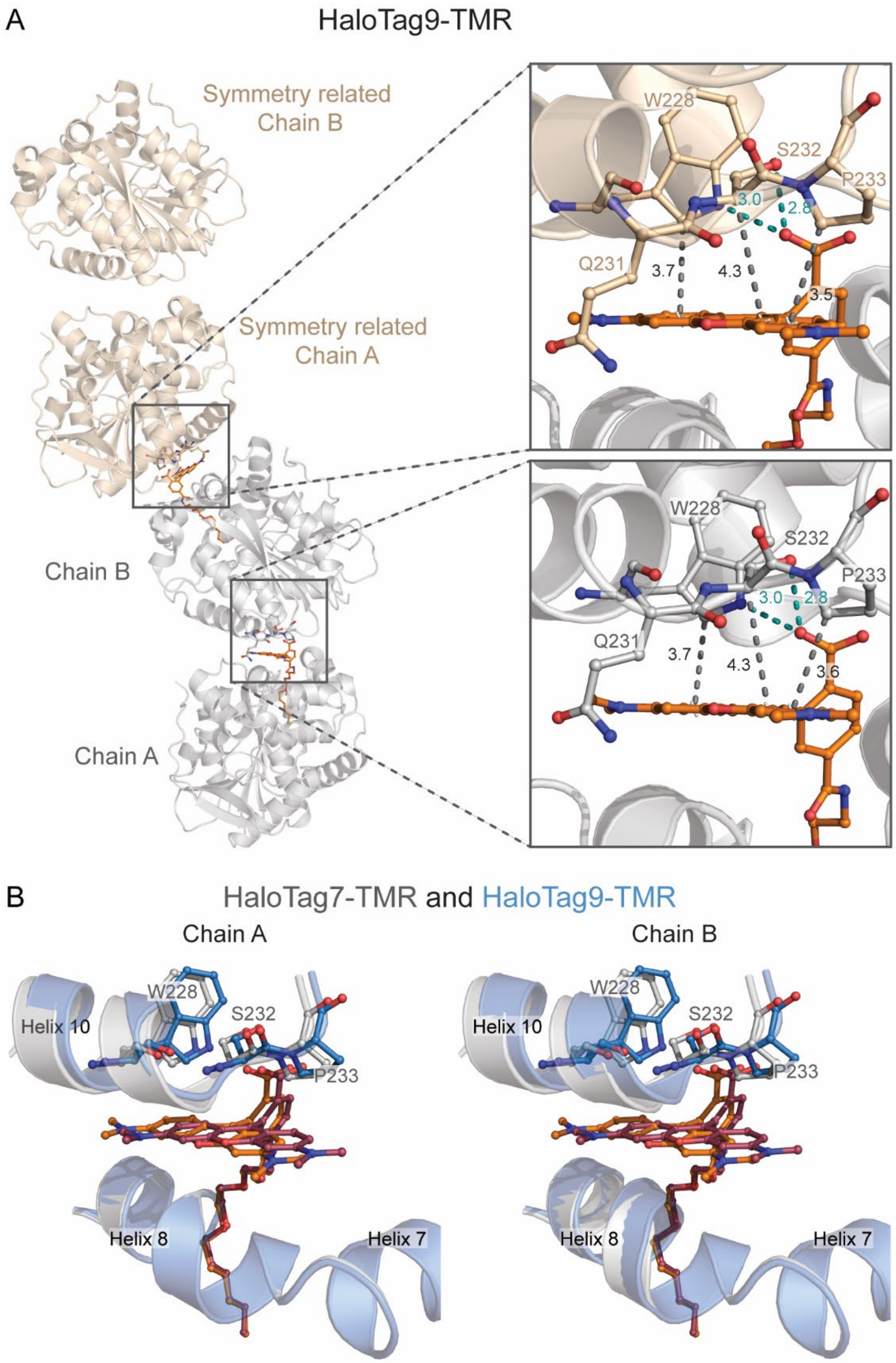

**Supplementary Figure S6:** Structural analysis of the interactions of TMR at the crystal packing interface. **A** HaloTag9-TMR (PDB ID: 6ZVY, 1.4 Å) crystal structure with the indicated zoom regions. The proteins are represented as cartoon (grey (asymmetric unit) or beige (symmetry related)). The bound TMR at the interface is represented as orange sticks. Important helix residues of the neighboring protein are represented as sticks together with characteristic distances, which are indicated in Ångström (e.g. hydrogen bonds: carboxylic acid of

TMR with S232 and W228 (teal dashes); distances from the xanthene ring to W228 (Cδ), S232 (Cα), and P233 (Cδ) (grey dashes)). In comparison to the structure of HaloTag7, the distances of the hydrogen bonds of the carboxylic acid do not change (Supplementary Table S10). The van der Waals distances increase for W228 (Cδ), remain constant for S232 (Cα), and decrease for P233 (Cδ) and the xanthene plane. This demonstrates that the fluorophore shifts from W288 towards S232 albeit the general interface remaining relatively similar. **B** Structural comparison of both chains of HaloTag9-TMR (**D**, PDB ID: 6ZVY, 1.4 Å) with HaloTag7-TMR at the packing interface. The HaloTag7 and HaloTag9 proteins are represented as cartoons in grey and blue, respectively. The bound TMR is given in stick representation in orange (HaloTag7) and violet (HaloTag9). Helices 7–8 of one monomer are represented with the end of helix 10 (224–234) of the adjacent monomer. Amino acids in close proximity to TMR are represented as sticks. The shift of the xanthene core of the fluorophore toward helix 7 is accompanied by a shift of the neighboring protein as indicated by the $RMSD_{xanth}$ compared to the $RMSD_{pack}$ of the W228, S232, and P233 residues (Supplementary Table 11; due to the two possible confirmations of Q231 in the HaloTag7 crystal structure this residue was omitted for RMSD calculations). Overall, the crystal packing of HaloTag7 and HaloTag9 is similar allowing the comparison of the relative fluorophore orientation.

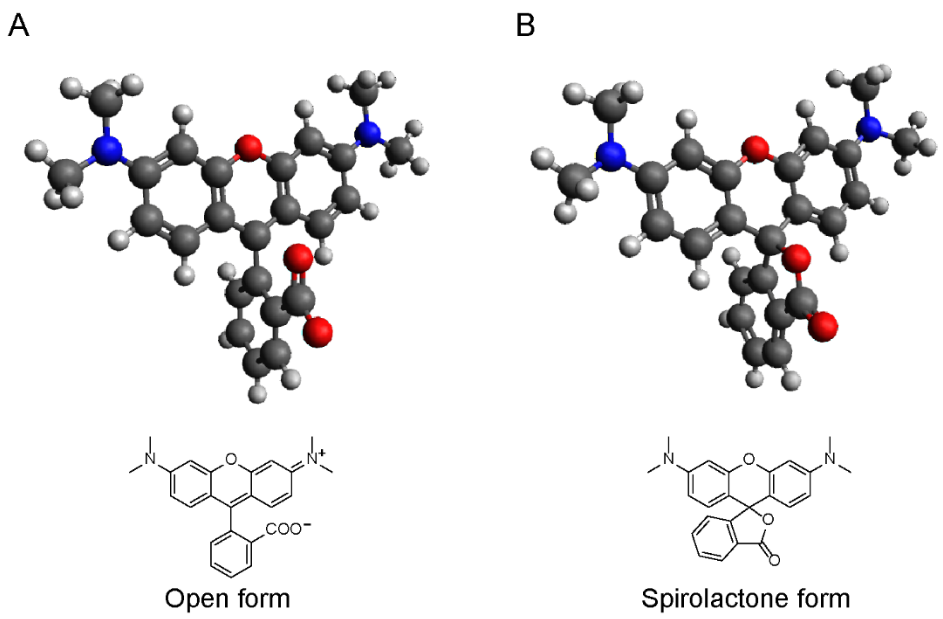

**Supplementary Figure S7:** Modeled structures of the open (**A**) and the spirolactone form (**B**) of TMR in water together with their chemical structures. The calculated angels $\varphi_{\text{Ar-Ar}}$ and $\gamma$ correspond to $\varphi_{\text{Ar-Ar}}$ = 94.3° and $\gamma$ = 5.6° for the open form and $\varphi_{\text{Ar-Ar}}$ = 115.1° and $\gamma$ = 37.8° for the closed form.

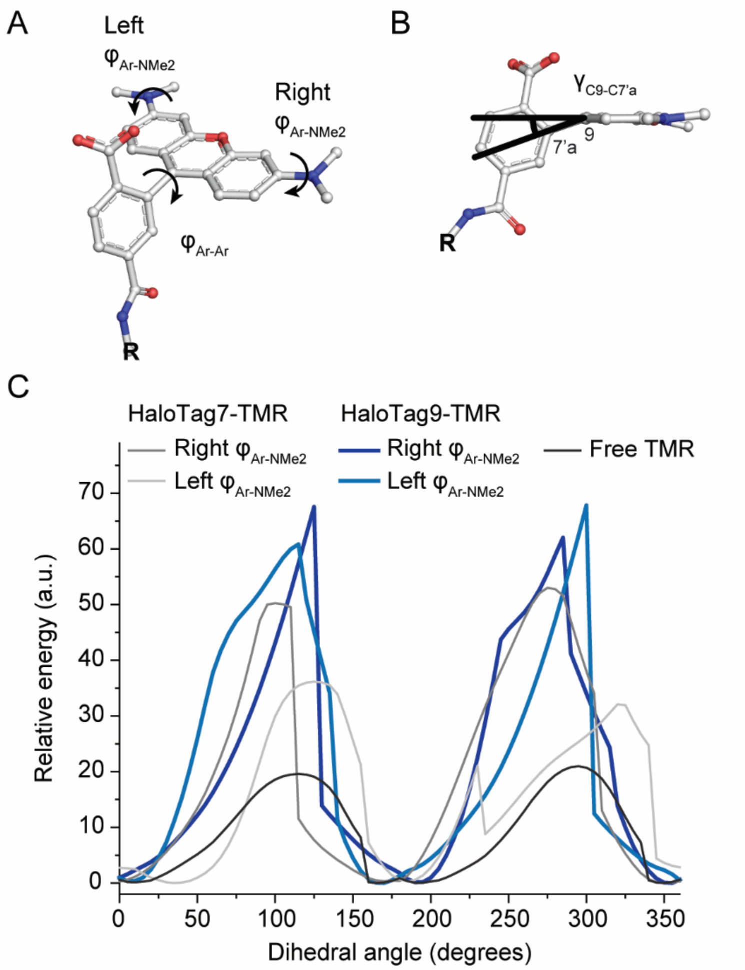

**Supplementary Figure S8:** Structural analysis of HaloTag7- or HaloTag9-TMR. **A** Dihedral angles $\varphi_{Ar\text{-}Ar}$ and $\varphi_{Ar\text{-}NMe2}$ left and right of TMR. The dihedral angle $\varphi_{Ar\text{-}Ar}$ is influenced by the protein, as it positions the xanthene core with respect to the chloroalkane, which is hold in place by two hydrogen bonds formed between the amid bond and T148 and T172. **B** Angle γ by which the C9-C7'a bond is tilted out of the xanthene plane. TMR on HaloTag9 and HaloTag7 showed angles of γ = 4.1° and 8.7°, respectively, which is in accordance with the calculated structures of the open form of TMR (5.6°) in water (Supplementary Fig. S7). A smaller angle in HaloTag9-TMR might indicate a greater $sp^2$ character of C9 and therefore a higher contribution of its remaining p orbital to the molecular orbitals relevant to the chromophore. This could positively influence the intrinsic extinction coefficient. **C** Relative energies of TMR when rotating the aniline group around $\varphi_{Ar\text{-}NMe2}$ (right or left). The rotational barrier is about three times (two times) as high for TMR on HaloTag9 (HaloTag7) when compared to unbound TMR in free solution. Energies given were calculated by molecular modeling using the force field OPLS3e.

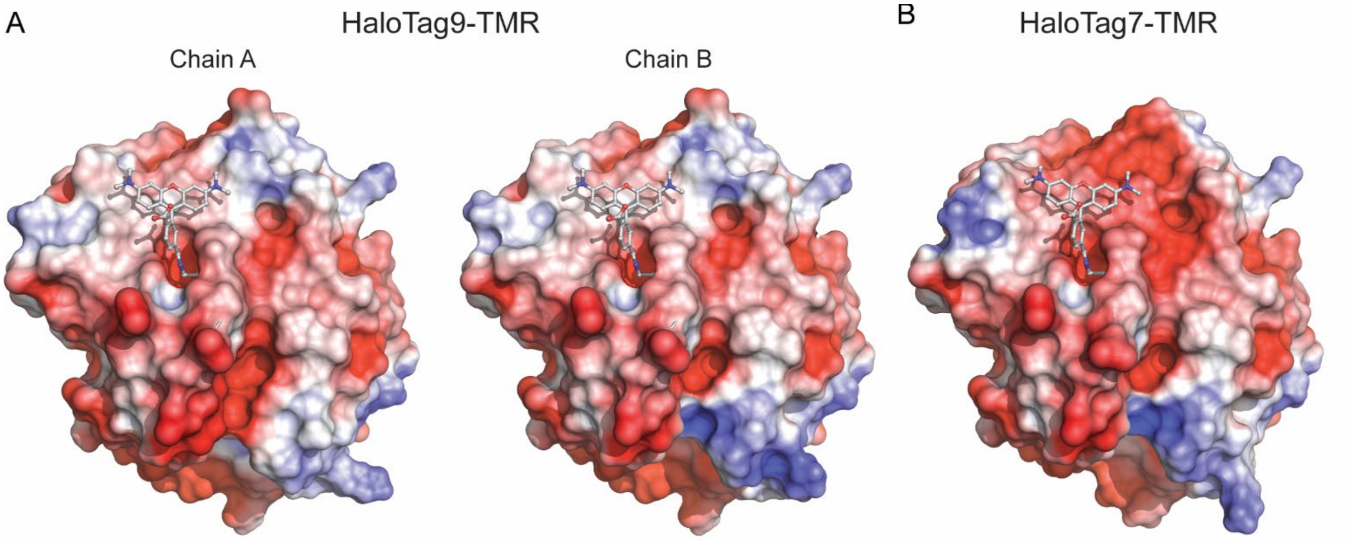

**Supplementary Figure S9:** Electrostatic potential of HaloTag9-TMR and HaloTag7-TMR mapped unto to their solvent excluded surface (Connolly surface). **A-B** The electrostatic surface potential of HaloTag9-TMR (PDB ID: 6ZVY, 1.4 Å, **A**) and HaloTag7-TMR (PDB ID: 6Y7A, 1.4 Å, **B**). The surface potential is more positive on HaloTag9 than for HaloTag7 especially around the fluorophore binding site. The potentials were obtained using the adaptive Poisson-Boltzmann Solver (APBS software) with standard parameters (0.15 M ionic strength in monovalent salt, 310.0 K, protein dielectric of 2, and solvent dielectric of 78.0). The surface potentials mapped unto the solvent excluded surface are on a [−3; 3] red–white–blue color map in units of $kJ \cdot mol^{-1} \cdot e^{-1}$. Calculations were performed using the APBS & PDB2PQR plug-in in Pymol[3].

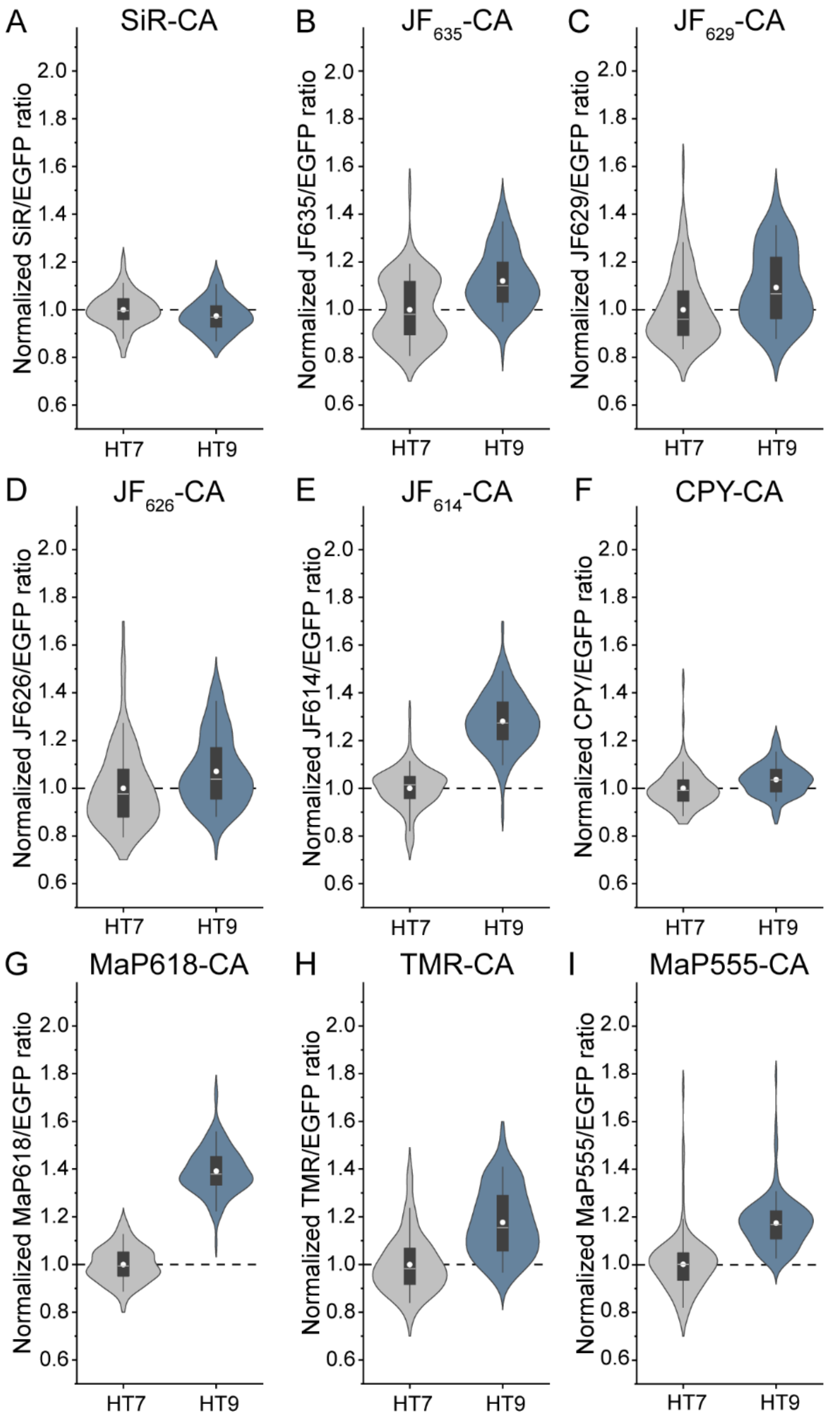

**Supplementary Figure S10:** Brightness comparison of HaloTag7 and HaloTag9 in mammalian cells by confocal microscopy. **A**-**I** Violin plots of normalized fluorescence intensities from living U-2 OS cells stably expressing either HaloTag7 or HaloTag9 in the cytosol. They were labeled with different fluorophores (1 µM, 3 h) and imaged by confocal microscopy. SiR-CA (**A**), $JF_{635}$-CA (**B**), $JF_{629}$-CA (**C**), $JF_{626}$-CA (**D**), $JF_{614}$-CA (**E**), CPY-CA (**F**), MaP618-CA (**G**), TMR-CA (**H**), and MaP555-CA (**I**). Distribution = light grey/blue, box = 25%–75% percentile, whiskers = 5%–95% percentile, white line = median, circle = mean, dashed line = mean of HaloTag7, *N* = 113–129 cells from three independent preparations (Supplementary Table S13).

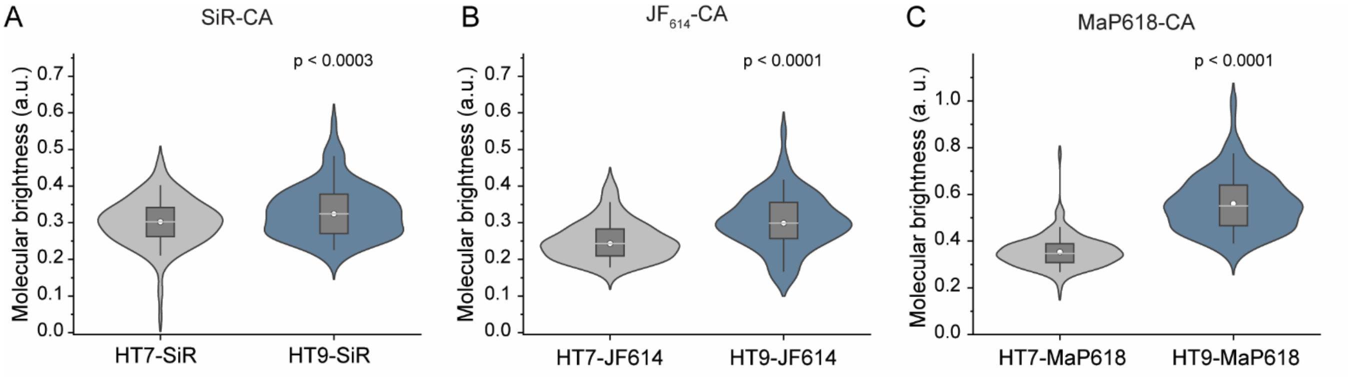

**Supplementary Figure S11:** Molecular brightness comparison of HaloTag7 and HaloTag9 in mammalian cells by FCS. **A-C** Violin plots of molecular brightness acquired by FCS in living U-2 OS cells expressing either HaloTag7 or HaloTag9 in the cytosol labeled with fluorophore (150 nM, 2 h, SiR-CA (**A**), $JF_{614}$-CA (**B**) and MaP618-CA (**C**)). 30 s long FCS traces were measured. Molecular brightness was then calculated using the amplitude of the autocorrelation curve as well as the mean fluorescence intensity. SiR-CA labeled HaloTag9 showed a 9±3% (mean±s.e.m.) higher brightness than HaloTag7, for $JF_{614}$-CA the change was 21±3%, and for MaP618-CA 59±4%. p-Values are given based one-sided t-tests (α = 5%, r = 0.05, 0.18, 0.27, Degrees of freedom (DF) = 296, 291, 323). Distribution = light grey/blue, box = 25%–75% percentile, whiskers = 5%–95% percentile, white line = median, circle = mean. *N* = SiR-CA: 141, 157; $JF_{614}$-CA: 148, 145; MaP618-CA: 155, 170 traces from 36 cells from three independent preparations (HaloTag7, HaloTag9).

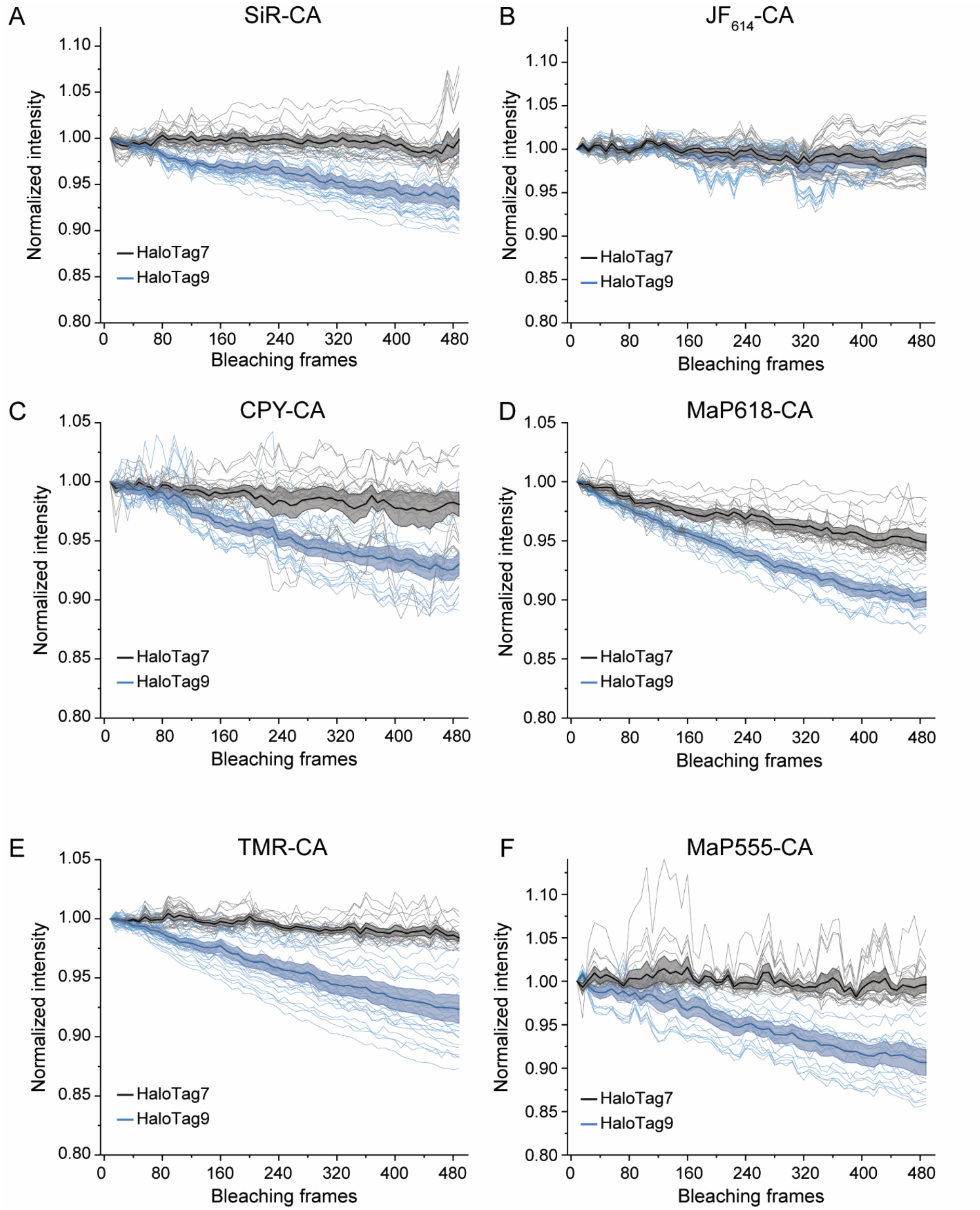

**Supplementary Figure S12:** Photostability of different fluorophores on HaloTag7 and HaloTag9 assessed by confocal microscopy. **A-F** Plots of normalized fluorescence intensity over time from living U-2 OS cells. U-2 OS cells stably expressing H2B-HaloTag7 or H2B-HaloTag9 were labeled with SiR-CA (**A**), $JF_{614}$-CA (**B**), CPY-CA (**C**), MaP618-CA (**D**), TMR-CA (**E**), and MaP555-CA (**F**) (1 µM, 3 h) and imaged by confocal microscopy over 480 bleaching cycles.

Photobleaching was induced by irradiation at maximal laser power and a z-stack to read out the fluorescence intensity was acquired every 8 bleaching cycles (thin lines = individual measurements; thick line and shaded area = mean±95% confidence interval). HaloTag9 was less photostable than HaloTag7 in all the experiments performed. The difference in fluorescence intensity in percentage after 480 photobleaching frames was SiR-CA: −6.7±1.5% (*N* = 25 and 30 nuclei); $JF_{614}$-CA: NS (*N* = 29 and 28 nuclei); CPY-CA: −5.1±1.4% (*N* = 26 and 26 nuclei); MaP618-CA: −5.0±1.0% (*N* = 24 and 23 nuclei); TMR-CA: −6.1±1.2% (*N* = 27 and 30 nuclei); MaP555-CA: −9.0±1.7% (*N* = 22 and 21 nuclei, all from three independent preparations). Albeit this difference being statistically significant (one sided t test, α = 5%, DF = 53, 55, 50, 45, 55, 41), its biological relevance is negligible. Indeed, in order to see 5% of photobleaching for HaloTag9 as compared to HaloTag7 one could measure 1,600 (555 nm), 2,600 (614 nm at 1.5%), 800 (614 nm at 5%), or 1,600 (630 nm at 2%) z-stacks. As the irradiation from 1 frame at maximal laser power (9.3 µW at 555 nm, 18.5 µW at 614 nm, and 17 µW at 630 nm) corresponds to 3.4x, 5.5x, 1.6x, and 3.4x higher energy deposition than for one z-stack in typical acquisition conditions (12 frames at 555 nm (2% = 0.23 µW), 614 nm (1.5% = 0.28 µW), 614 nm (5% = 0.95 µW), or 630 nm (2% = 0.42 µW)) for TMR-CA/MaP555-CA, CPY-CA/MaP618-CA, $JF_{614}$-CA, or SiR-CA.

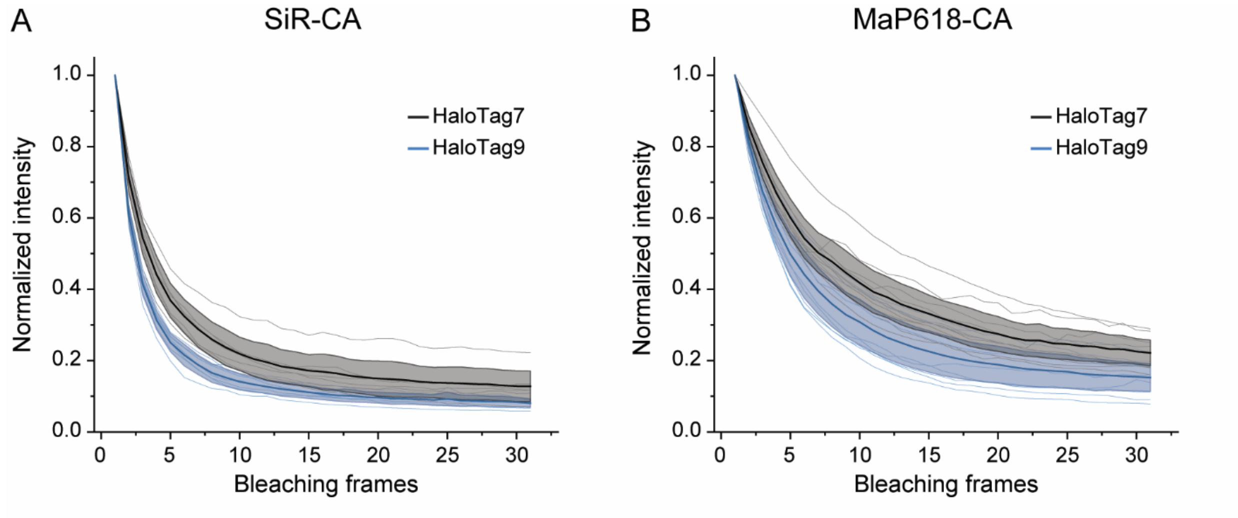

**Supplementary Figure S13:** Photostability of different fluorophores on HaloTag7 and HaloTag9 assessed by STED microscopy. **A-B** Plots of normalized fluorescence intensity over time from living U-2 OS cells. U-2 OS cells expressing Tomm20-HaloTag7 or Tomm20-HaloTag9 were labeled with SiR-CA (**A**) or MaP618-CA (**B**) (1 µM, 3 h) and imaged by STED microscopy for 30 consecutive frames (thin lines = individual measurements; thick line and shaded area = mean±95% confidence interval, *N* = 7 (SiR-CA) or 9 (MaP618-CA) measurements from three independent preparations). Fluorescence intensity is rapidly dropping and whereas there is a small difference between HaloTag7 and HaloTag9 labeled with SiR-CA during the first 10 frames there is no difference for the later 20 frames or for the experiment using MaP618-CA. We therefore conclude that the small difference in photostability found by confocal microscopy (Supplementary Fig. S12) has little biological relevance and no implications for experiments such as the STED microscopy presented here.

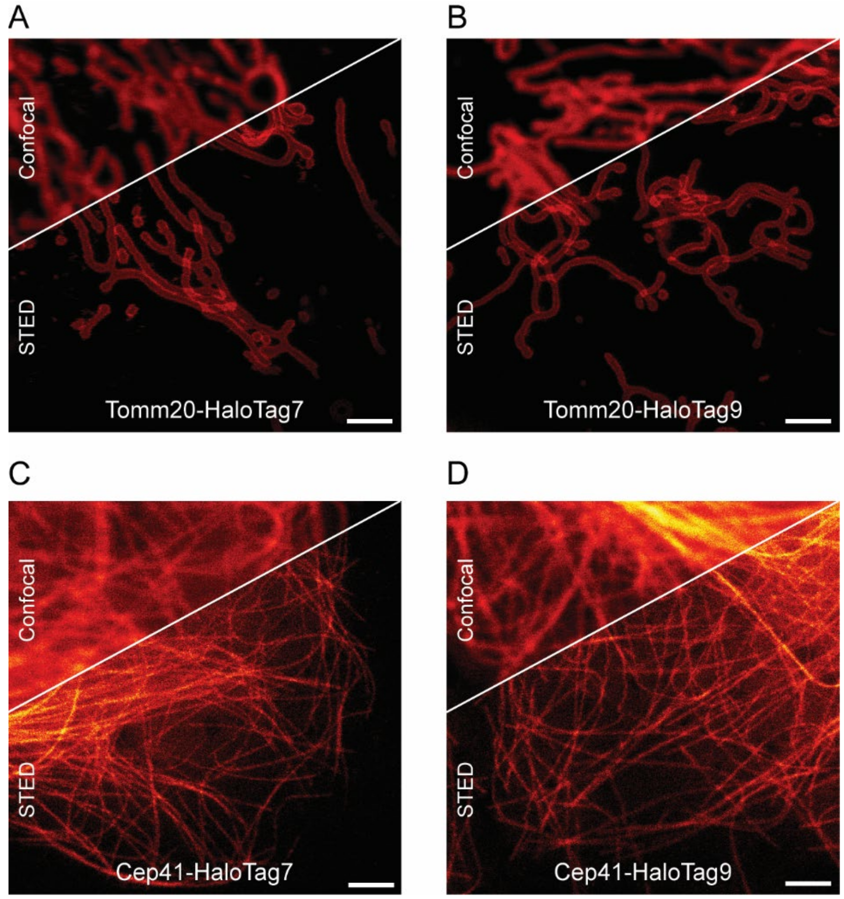

**Supplementary Figure S14:** Confocal and STED fluorescence microscopy images of mitochondria or microtubules labeled via HaloTag7 or HaloTag9. **A-B** U-2 OS cells stably expressing the outer mitochondrial membrane protein Tomm20 as a fusion with HaloTag7 (**A**) or HaloTag9 (**B**) labeled with MaP618-CA (1 µM, 3 h). **C-D** U-2 OS cells stably expressing the microtubule marker Cep41 as a fusion of HaloTag7 (**A**) or HaloTag9 (**B**) labeled with MaP618-CA (1 µM, 3 h). Scale bars, 2 µm.

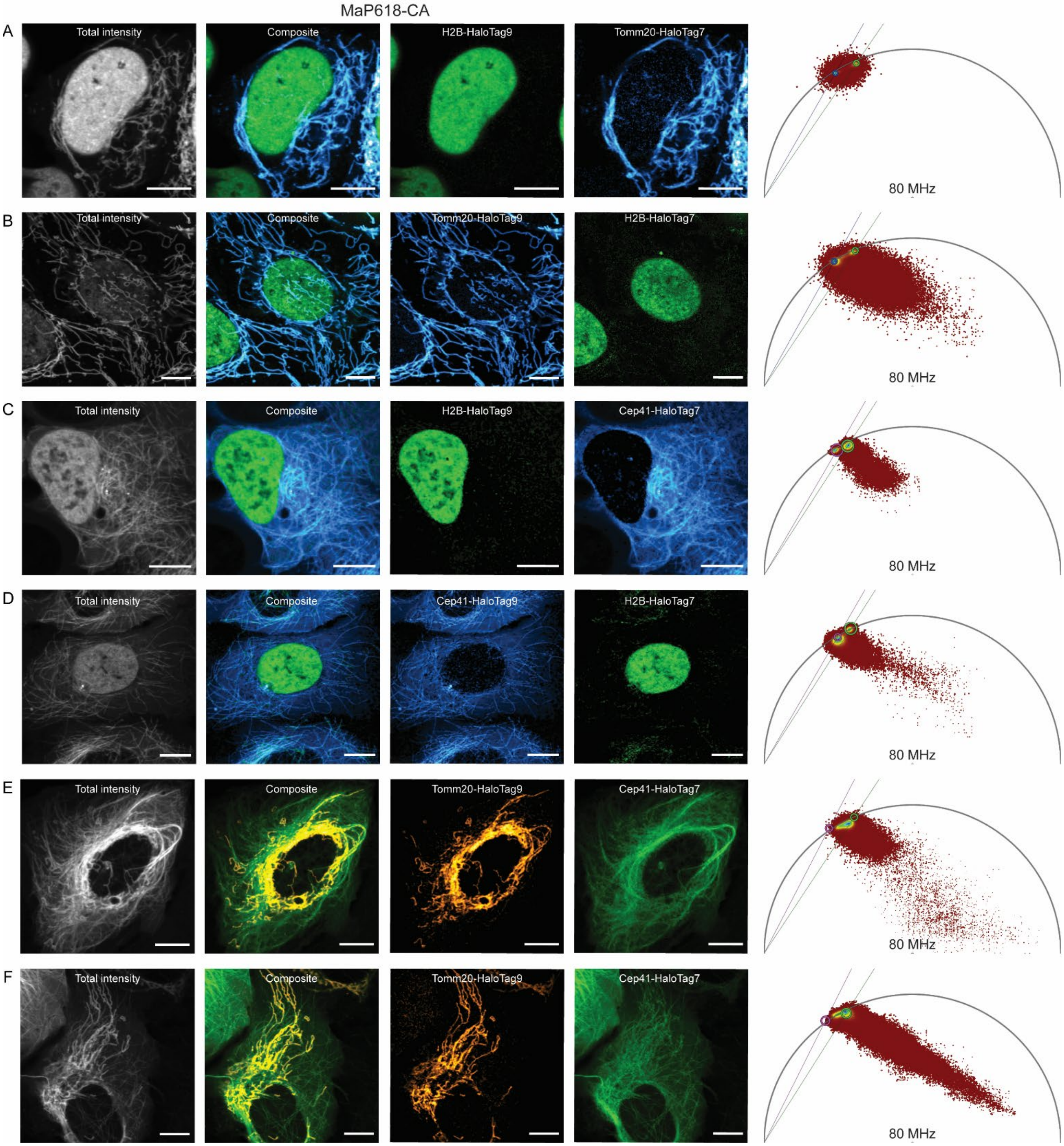

**Supplementary Figure S15:** Live-cell fluorescence lifetime multiplexing of different combinations of HaloTag7 and HaloTag9 fusion proteins labeled with a single fluorophore. **A** U-2 OS cells stably expressing H2B-HaloTag9 and transiently Tomm20-HaloTag7 labeled with MaP618-CA (1 µM, 3 h). **B** Inverse combination: Tomm20-HaloTag9 (stable) and H2B-HaloTag7 (transient) labeled with MaP618-CA (1 µM, 3 h). **C-D** Cep41-HaloTag7 (**C**) or HaloTag9 (**D**) stably expressed in U-2 OS cells and additionally transiently expressing H2B-HaloTag9 (**C**) or HaloTag7 (**D**) labeled with MaP618-CA (1 µM, 3 h). Fusion with Cep41 shifted the fluorescence lifetime of MaP618-CA on

HaloTag7 or HaloTag9 to slightly longer/shorter fluorescence lifetimes as compared to H2B or Tomm20 fusions (H2B-HaloTag7 = 3.0 ns and H2B-HaloTag9 = 3.6 ns vs. Cep41-HaloTag7 = 3.1 ns and Cep41-HaloTag9 = 3.4 ns). The phasor plots reveal that the clusters for Cep41 move away from the universal circle indicating a multi-exponential fluorescence lifetime decay. This and the associated change in fluorescence lifetime might originate from interactions of the fluorophore with the microtubule environment. **E-F** Cep41-HaloTag7 stably expressed in U-2 OS cells and additionally transiently expressing Tomm20-HaloTag9 labeled with MaP618-CA (1 µM, 3 h). Separation of the inverse combination Cep41-HaloTag9 and Tomm20-HaloTag7 was hindered by significantly different levels of expression of the target proteins, which prevented identifying different populations on the phasor plot. The fluorescence intensity, the composite, the two individual separated species as well as the corresponding wavelet-filtered phasor plot used for separation are given. Species separation was achieved using the phasor approach. Scale bars, 10 µm.

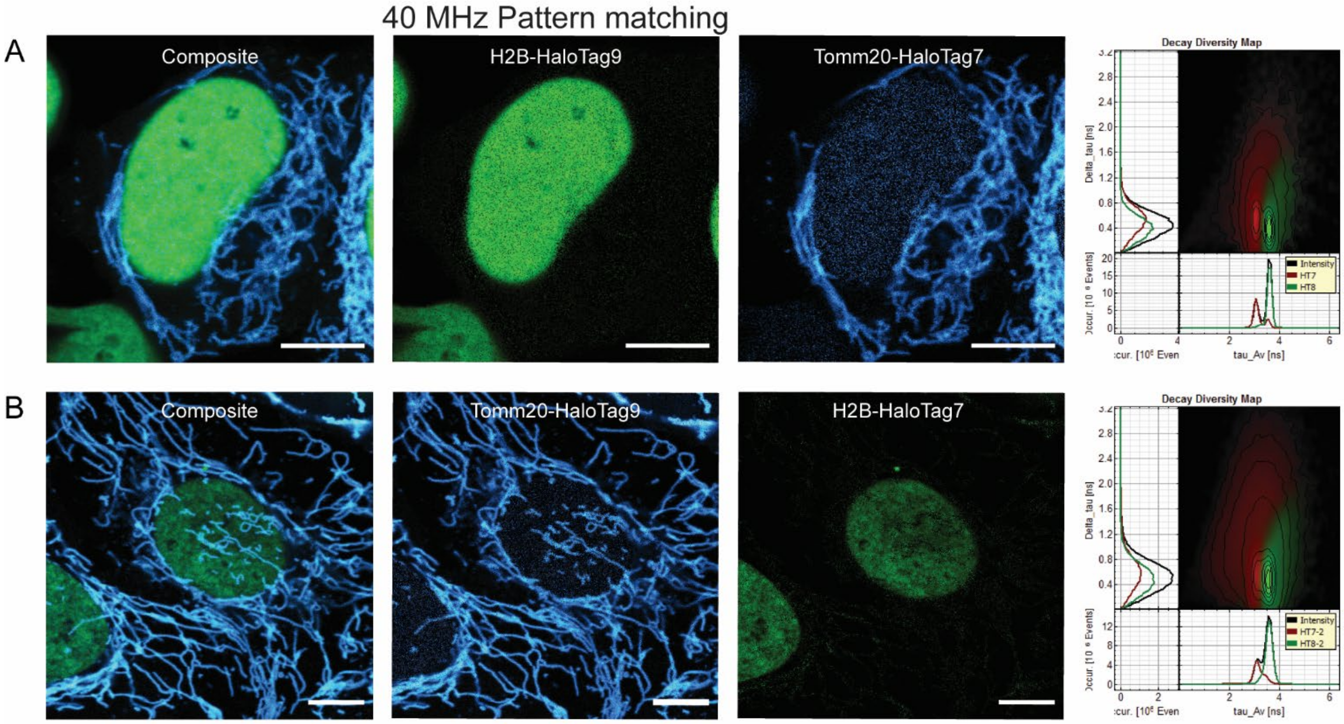

**Supplementary Figure S16:** Live-cell fluorescence lifetime multiplexing using pattern matching analysis to separate the two components. **A-B** Analysis of the FLIM images given in Supplementary Fig. S15A-B via pattern matching gave similar results as the separation using the phasor plot. However, only images acquired at 40 MHz instead of 80 MHz could be separated reliably using pattern matching. The composite, the two individual separated components as well as the corresponding decay diversity map used for separation are given. The presented images were acquired under the same conditions as the images in Supplementary Fig. S15A-B apart from the laser repetition rate, which was set to 40 MHz instead of 80 MHz. Scale bars, 10 µm.

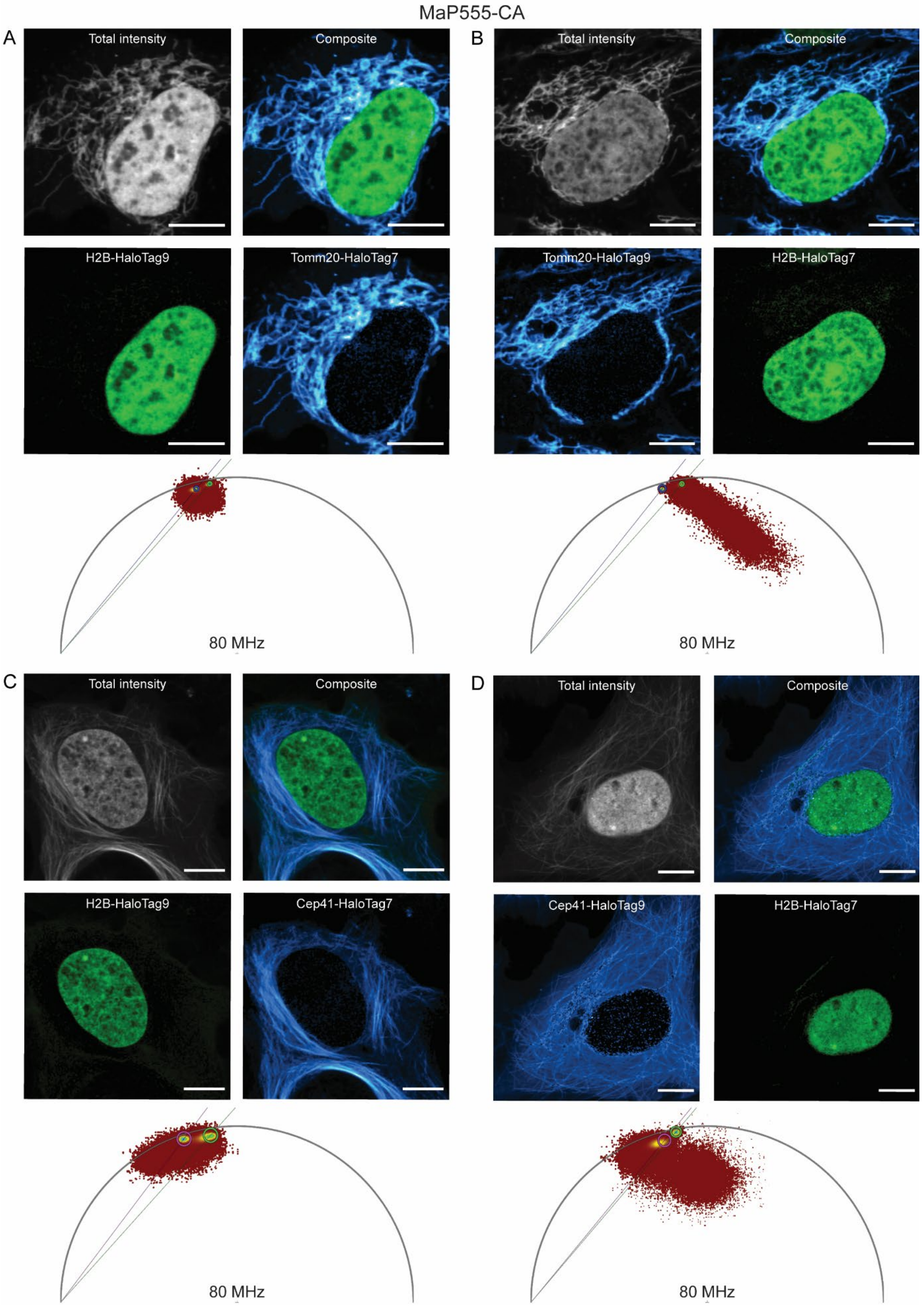

MaP555-CA
A
Total intensity
Composite
H2B-HaloTag9
Tomm20-HaloTag7
80 MHz
B
Total intensity
Composite
Tomm20-HaloTag9
H2B-HaloTag7
80 MHz
C
Total intensity
Composite
H2B-HaloTag9
Cep41-HaloTag7
80 MHz
D
Total intensity
Composite
Cep41-HaloTag9
H2B-HaloTag7
80 MHz

**Supplementary Figure S17:** Live-cell fluorescence lifetime multiplexing using MaP555-CA instead of MaP618-CA. **A-B** U-2 OS cells stably expressing H2B-HaloTag9 and transiently Tomm20-HaloTag7 (**A**) or the inverse combination (Tomm20-HaloTag9 (stable) and H2B-HaloTag7 (transient), **B**) labeled with MaP555-CA (1 µM, 3 h). **C-D** Cep41-HaloTag7 (**C**) or HaloTag9 (**D**) stably expressed in U-2 OS cells and additionally transiently expressing H2B-HaloTag9 (**C**) or HaloTag7 (**D**) labeled with MaP555-CA (1 µM, 3 h). The Cep41-fusions show altered fluorescence lifetime in comparison to the H2B or Tomm20 fusions. The fluorescence intensity, the composite, the two individual separated species as well as the corresponding wavelet-filtered phasor plot used for separation are given. Species separation was achieved using the phasor approach. Scale bars, 10 µm.

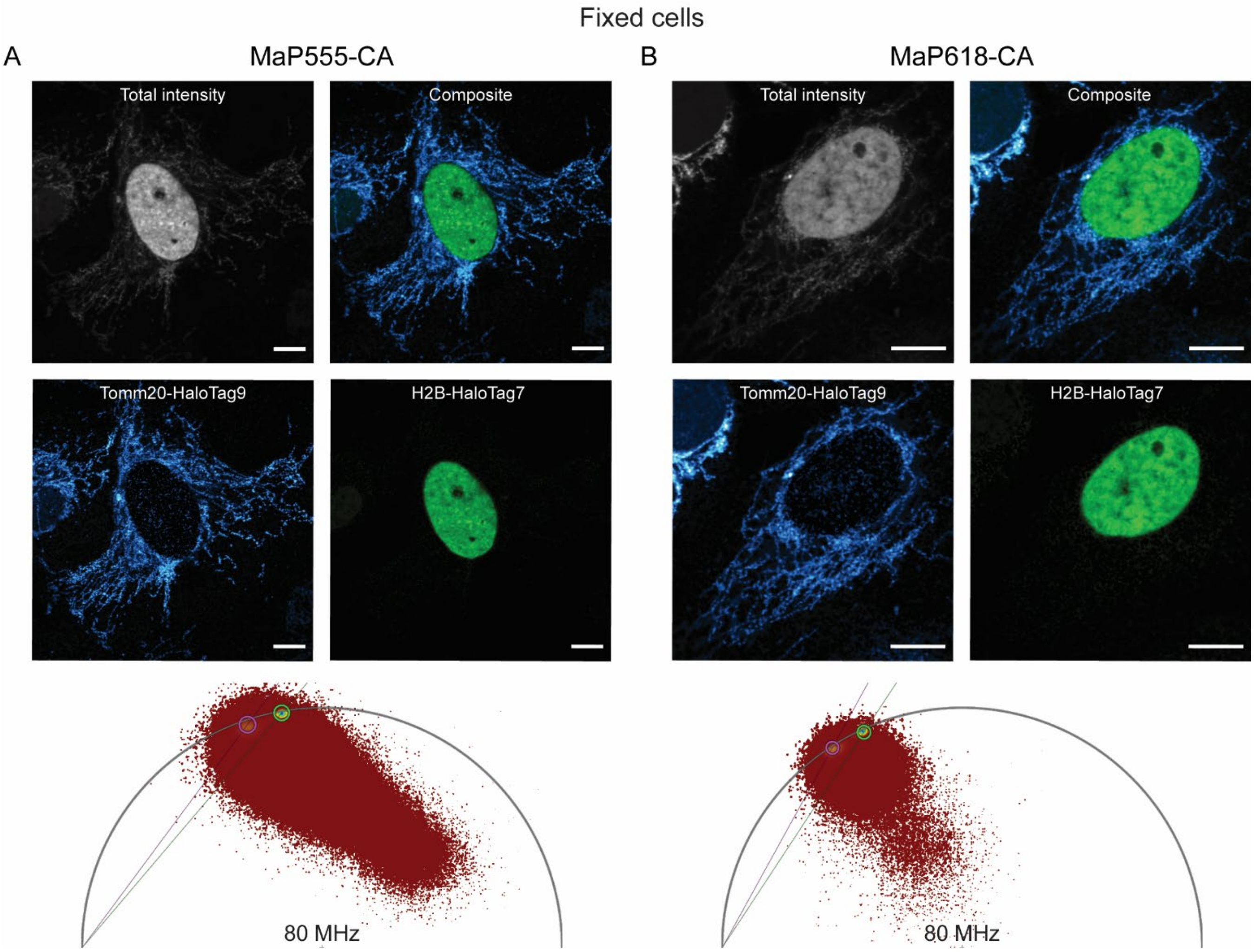

**Supplementary Figure 18:** Fluorescence lifetime multiplexing in fixed cells. **A-B** U-2 OS cells stably expressing Tomm20-HaloTag9 and transiently H2B-HaloTag7 were fixed with PFA and labeled with MaP618-CA (1 µM, 3 h, **A**) or MaP555-CA (1 µM, 3 h, **B**). The identification of both structures also works in fixed cells. The fluorescence intensity, the composite, the two individual separated species as well as the corresponding wavelet-filtered phasor plot used for separation are given. Species separation was achieved using the phasor approach. Scale bars, 10 µm.

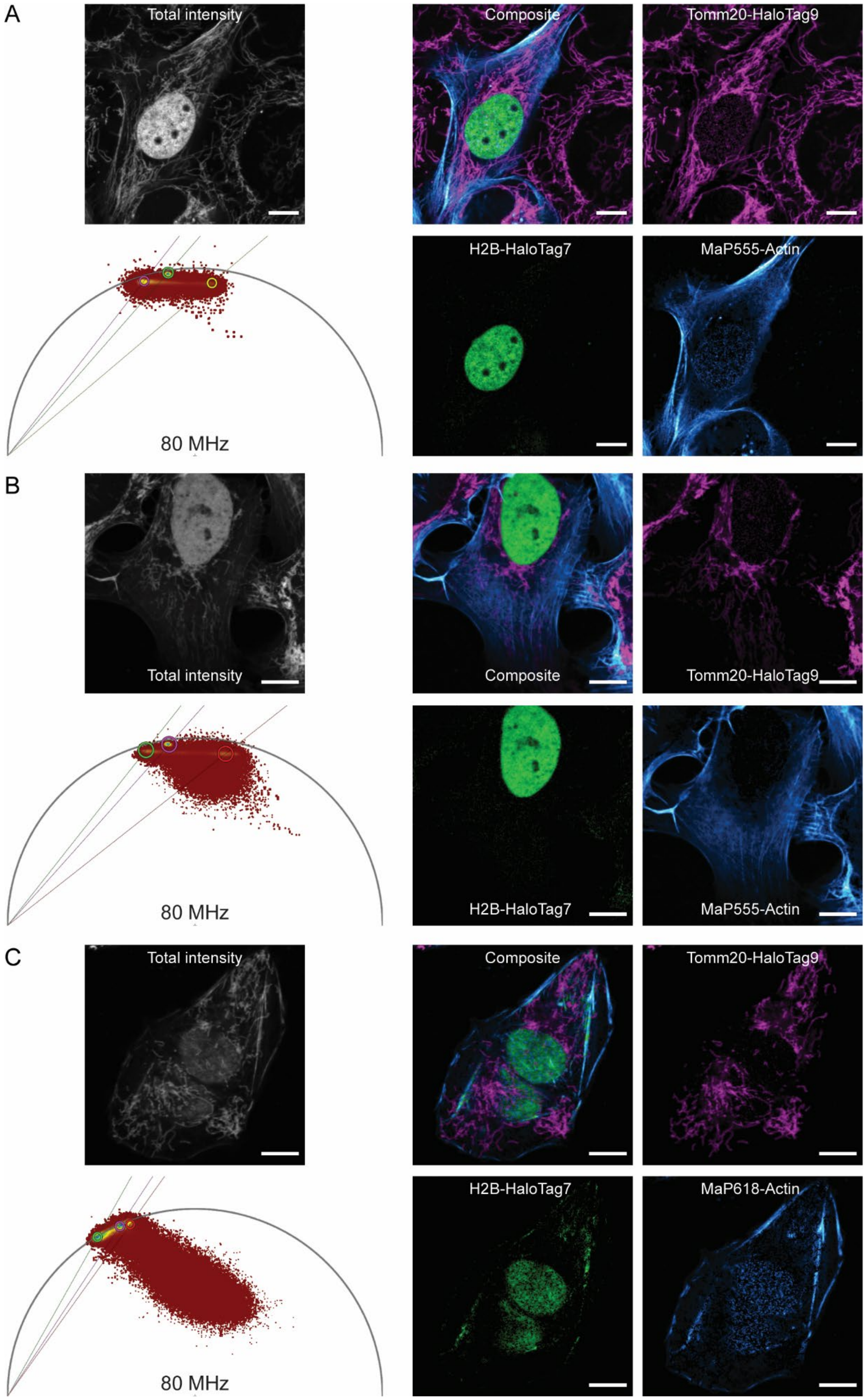
A
Total intensity
Composite
Tomm20-HaloTag9
80 MHz
H2B-HaloTag7
MaP555-Actin
B
Total intensity
Composite
Tomm20-HaloTag9
80 MHz
H2B-HaloTag7
MaP555-Actin
C
Total intensity
Composite
Tomm20-HaloTag9
80 MHz
H2B-HaloTag7
MaP618-Actin

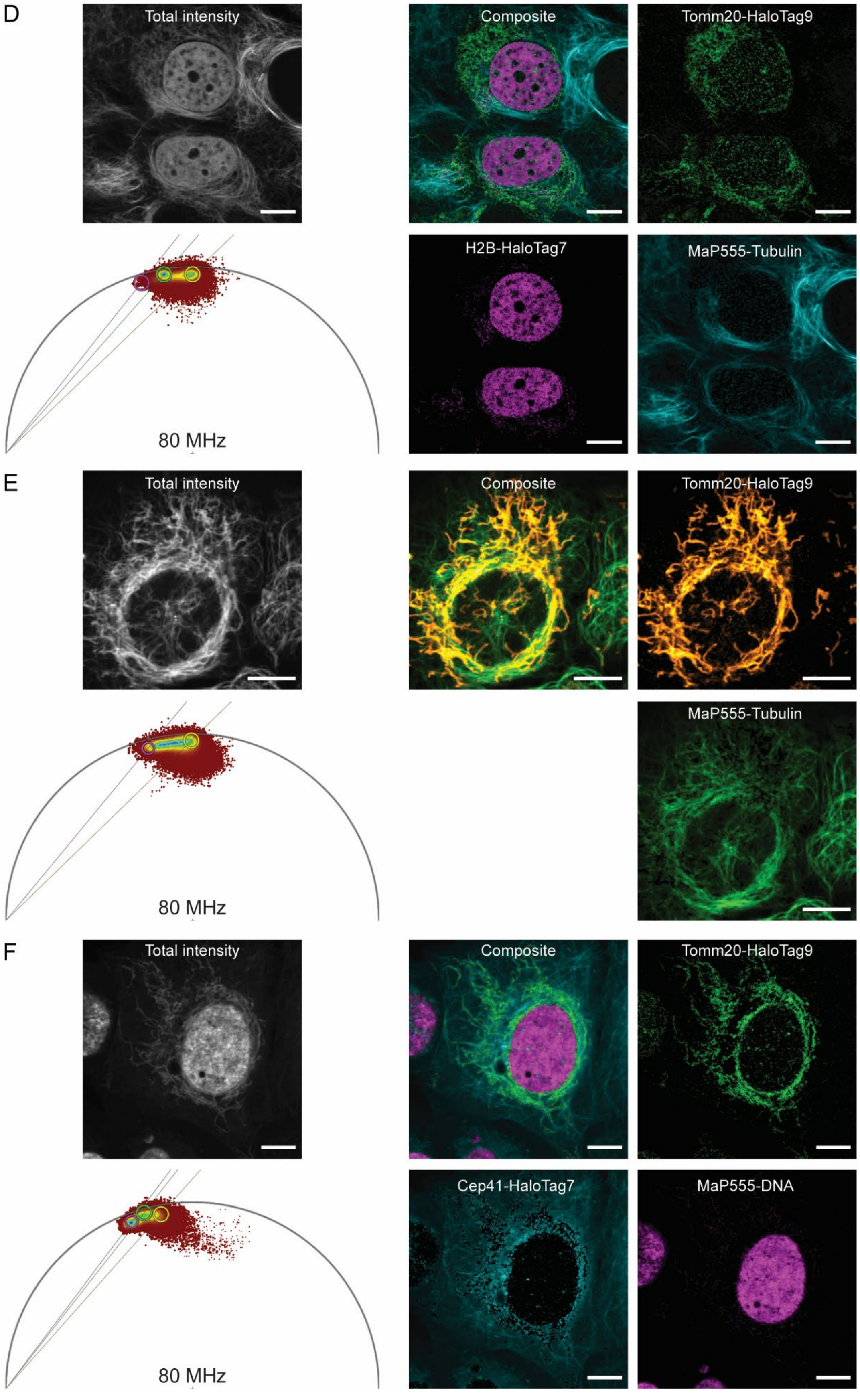
D
Total intensity
Composite
Tomm20-HaloTag9
H2B-HaloTag7
MaP555-Tubulin
80 MHz
E
Total intensity
Composite
Tomm20-HaloTag9
MaP555-Tubulin
80 MHz
F
Total intensity
Composite
Tomm20-HaloTag9
Cep41-HaloTag7
MaP555-DNA
80 MHz

**Supplementary Figure S19:** Live-cell fluorescence lifetime multiplexing of three instead of two species. **A-B** Live U-2 OS cells stably expressing Tomm20-HaloTag9 and transiently H2B-HaloTag7 were labeled with MaP555-CA (1 µM, 3 h) and MaP555-Actin (2 µM, 3 h). The three structures can be clearly separated. **C** Live U-2 OS cells stably expressing Tomm20-HaloTag9 and transiently H2B-HaloTag7 were labeled with MaP618-CA (1 µM, 3 h0 and MaP618-Actin (2 µM, 3 h). Separation was more challenging as the fluorescence lifetime of MaP618-Actin was very similar to HaloTag7-MaP618 and the three clusters were lying almost on a line and not forming a triangle in the phasor space. **D** U-2 OS cells stably expressing Tomm20-HaloTag9 and transiently H2B-HaloTag7 were labeled with MaP555-CA (1 µM, 3 h) as well as MaP555-Tubulin (2 µM, 3 h). MaP555-Tubulin showed a fluorescence lifetime of 2.0 ns and can therefore be separated from the fluorescence lifetimes of MaP555-CA. However, the separation from HaloTag9 (2.6 ns) is performed more easily than from HaloTag7 (2.3 ns). **E** U-2 OS cells expressing only Tomm20-HaloTag9 labeled with MaP555-CA (1 µM, 3 h) as well as MaP555-Tubulin (2 µM, 3 h), demonstrating that separation of these two species works very well. **F** U-2 OS cells stably expressing Cep41-HaloTag7 and transiently Tomm20-HaloTag9 labeled with MaP555-CA(1 µM, 3 h) and MaP555-DNA (1 µM, 3 h). The fluorescence lifetime of MaP555-DNA (2.7 ns) is very close to the fluorescence lifetime of HaloTag9-MaP555 (2.6 ns) and separation was therefore complex. Similar experiments using MaP618-CA and MaP618-DNA were performed but the images acquired were not separable as the fluorescence lifetime of HaloTag9-MaP618 and MaP618-DNA were too similar. The fluorescence intensity, the composite, the two individual separated species as well as the corresponding wavelet-filtered phasor plot used for separation are given. Species separation was achieved using the phasor approach. Scale bars, 10 µm.

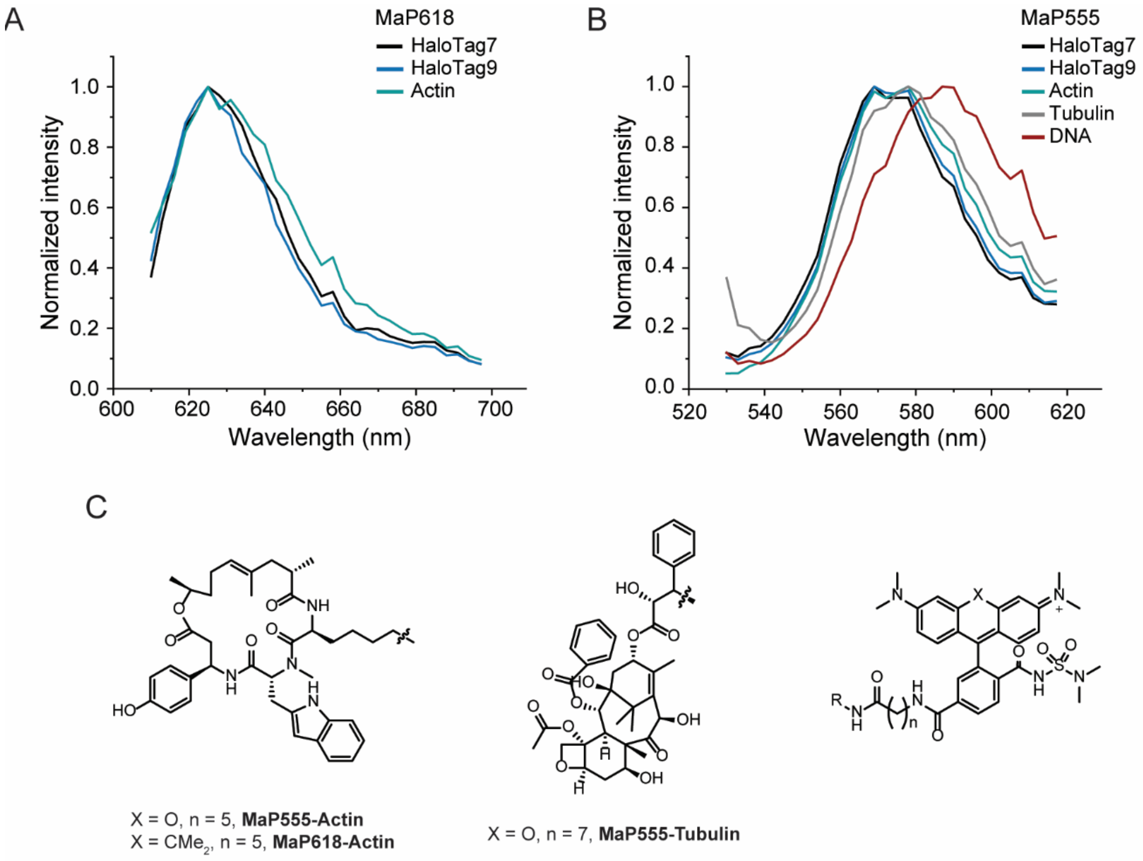

**Supplementary Figure S20:** Spectral characterization of MaP555 and MaP618 based fluorescent probes. **A-B** Emission spectra of MaP618 (**A**) and MaP555 (**B**) fluorophores on different targets as measured by confocal microscopy. Only the emission spectrum of MaP555-DNA showed a bathochromic shift compared to the other probes. **C** Chemical structures of MaP555-Actin, MaP618-Actin, and MaP555-Tubulin used for three target FLIM imaging. The structure for MaP555-DNA is unknown.

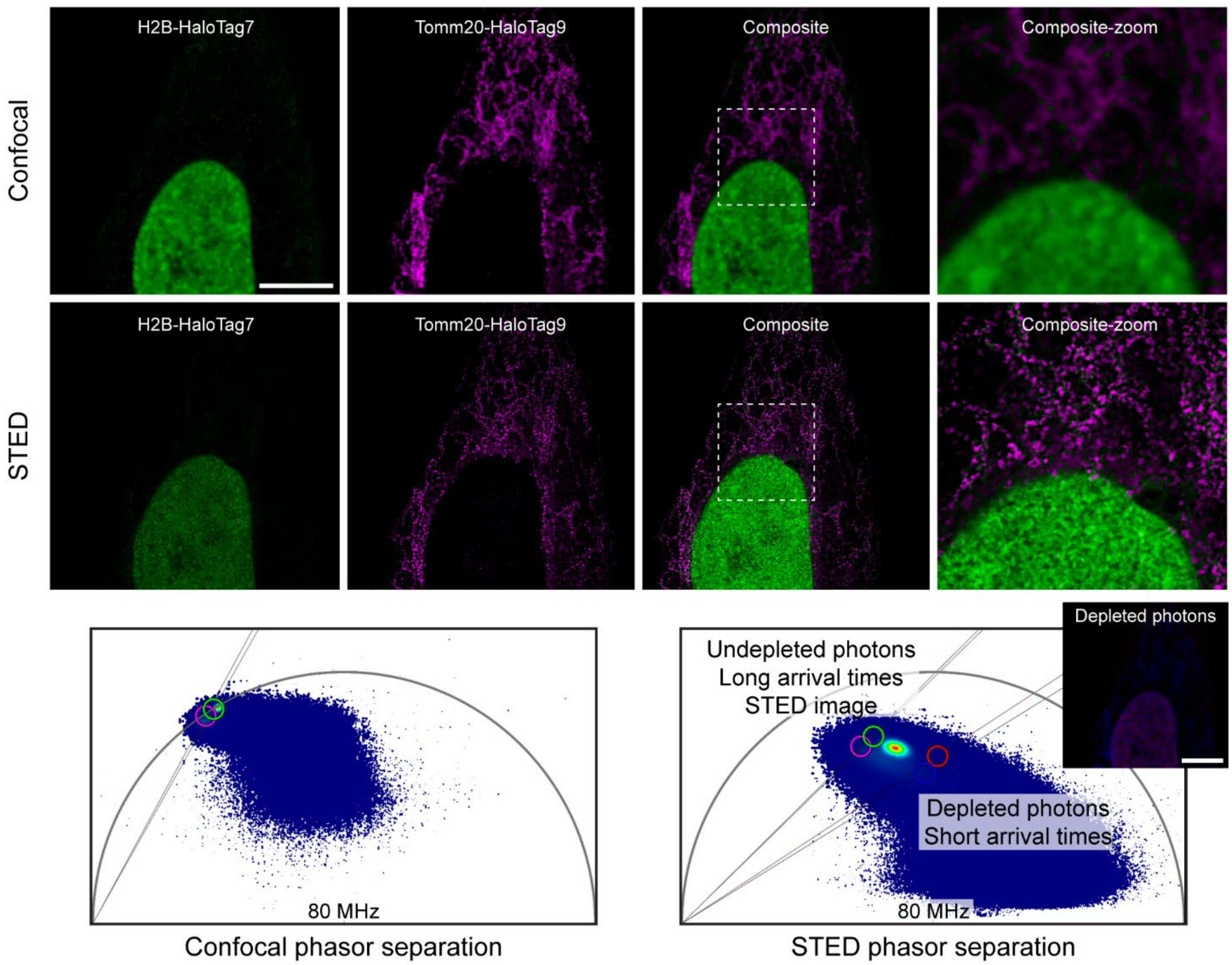

**Supplementary Figure S21:** Fixed cell fluorescence lifetime multiplexing in STED microscopy. U-2 OS cells stably expressing Tomm20-HaloTag9 and transiently H2B-HaloTag7 were fixed with PFA and labeled with MaP618-CA (1 µM, 3 h). Confocal and STED-FLIM images were simultaneously acquired (line sequential) and species separation performed using the phasor approach. The two individual separated species, the composite, a zoom-in and the corresponding wavelet-filtered phasor plots used for separation both in confocal and STED (magenta and green regions) are given. Additionally, the composite image generated using the depleted photons (blue and red region in the STED phasor plot) is given. Scale bars, 5 µm.

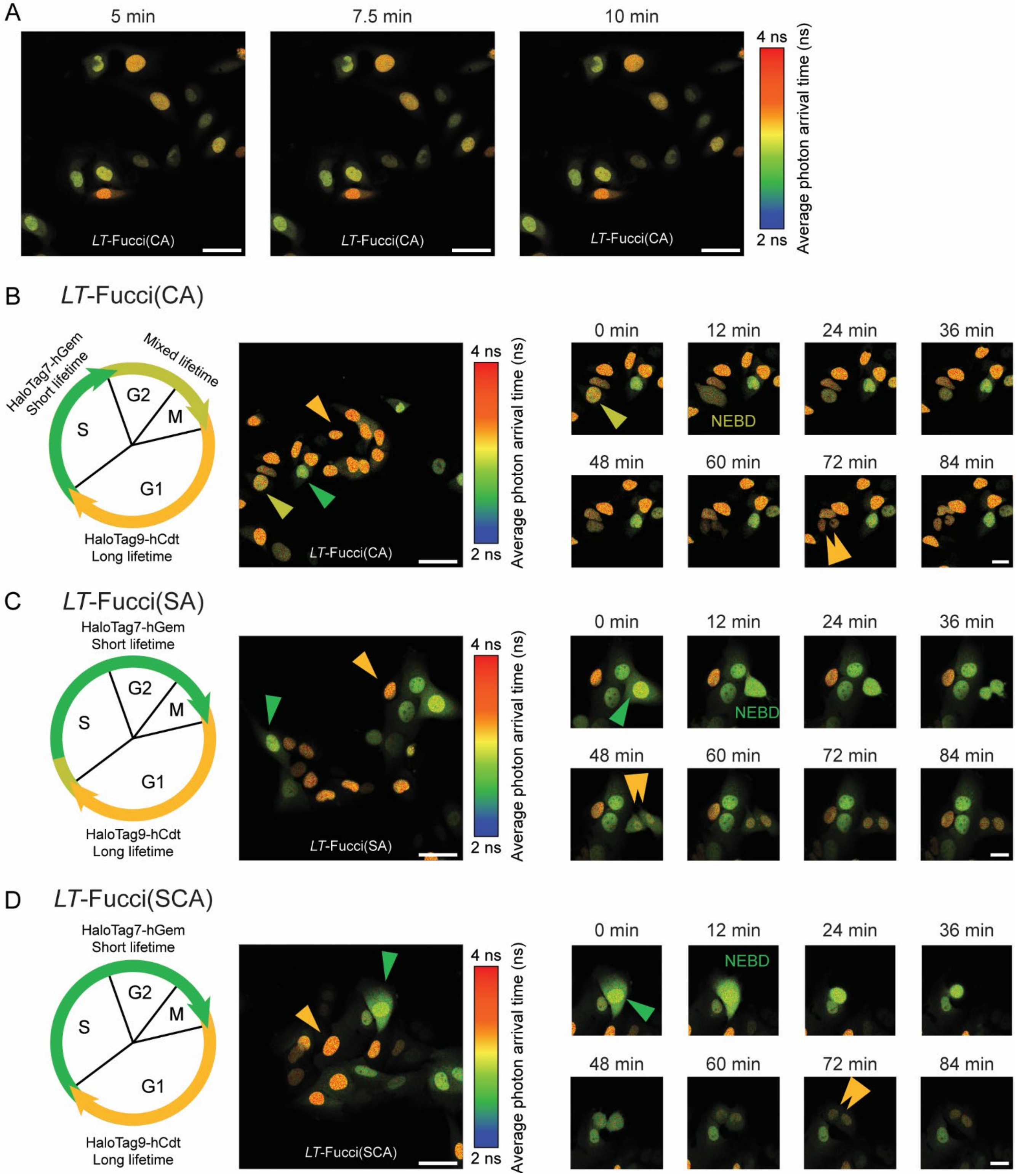

**Supplementary Figure S22:** Characterization of the *LT*-Fucci biosensors. **A** Assessment of the labeling speed of *LT*-Fucci(CA) with MaP618-CA (1 µM). Within 5 min the biosensor was sufficiently labeled. **B** *LT*-Fucci(CA). **C** *LT*-Fucci(SA). **D** *LT*-Fucci(SCA). For each biosensor a schematic overview of the predominant species present in each phase of the cell cycle is given. For all biosensors HaloTag9-hCdt is present during G1 and the nuclei will therefore present long average photon arrival times (MaP618-CA: 3.7 ns, orange). During S, G2 and M phase HaloTag7-hGem or a mixture of HaloTag7-hGem and HaloTag9-hCdt is present, resulting in short average photon arrival times (MaP618-CA: 3.1 ns, green) or a gradient of medium average photon arrival times (MaP618-CA: ~3.4 ns, light-green). A representative FastFLIM image of U-2 OS cells stably expressing the lifetime-biosensors labeled with MaP618-CA (1 µM) is given. Colored arrowheads indicate cells in the respective cell cycle phases. Providing 1 µM of MaP618-CA was sufficient to follow the division of cells over 24 h (Supplementary videos 1-3). Representative images of individual cells dividing for each cell line are given. The cells dividing ((light)-green arrowheads), the two daughter cells (orange arrowheads) as well as the moment of nuclear envelope breakdown (NEBD) are indicated. Scale bars, 50 µm and 25 µm.

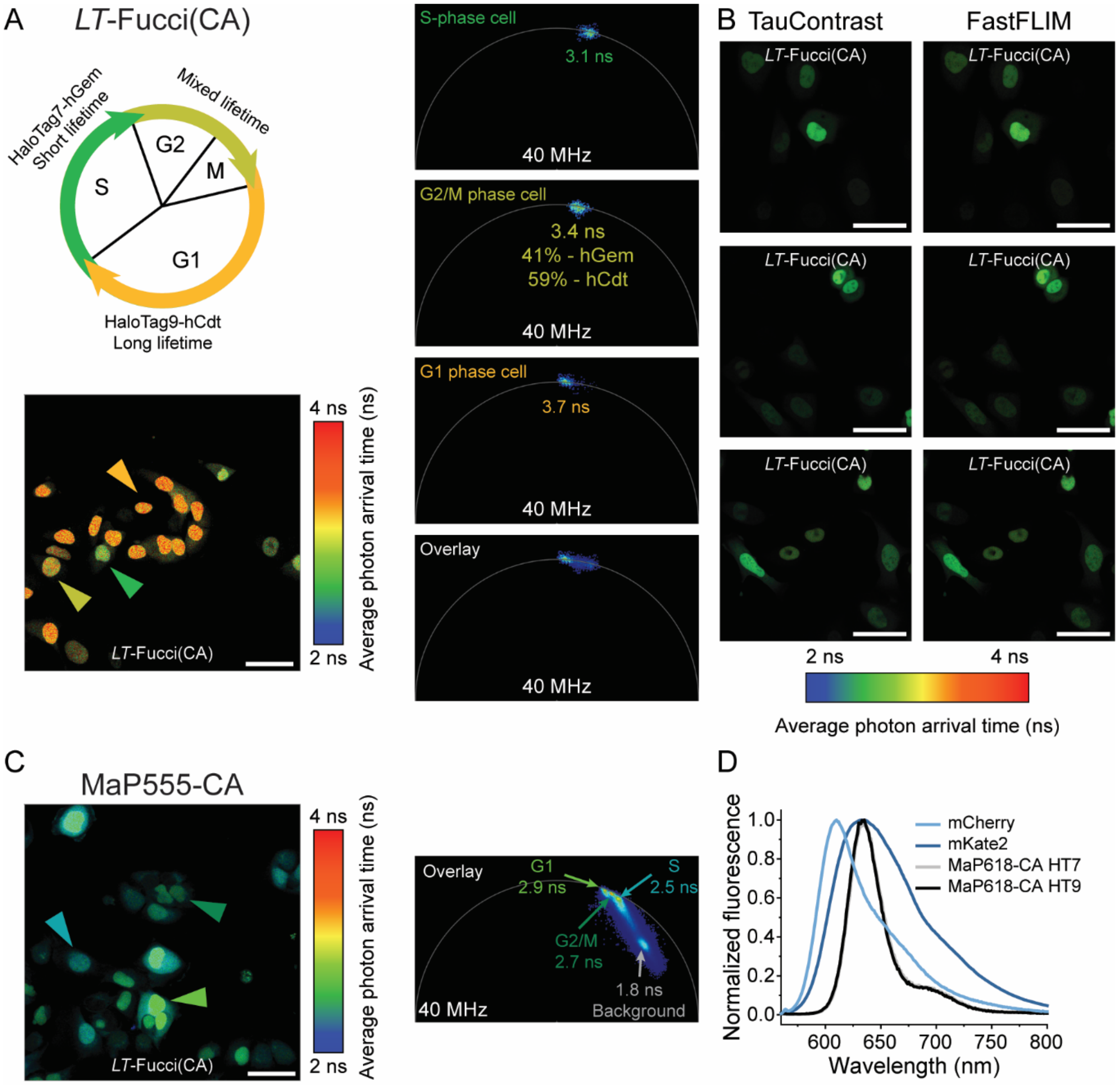

**Supplementary Figure S23:** Phasor analysis of the *LT*-Fucci(CA) biosensor. **A** Fluorescence lifetime analysis of three nuclei from the image of *LT*-Fucci(CA) (Fig. 2H-I). The average fluorescence lifetime of the nuclei indicated was evaluated using phasor analysis and the respective clusters are given in the phasor plots. In addition, the overlay of all three phasor plots was used to evaluate the percentages of hGem and hCdt present in the G2/M phase cell. As the fluorescence of a G2/M phase cell shows contributions from the two individual components HaloTag7-hGem and HaloTag9-hCdt, the law of linear addition in phasor space can be applied. Scale bars, 50 µm. **B** Lifetime-based biosensing on a confocal microscope without FLIM module. The average photon arrival time of *LT*-Fucci(CA) labeled with MaP618-CA (1 µM) was measured using TauContrast on a STELLARIS microscope[4]. This allows for a direct, lifetime-based readout of the biosensor, without the need for dedicated and complex FLIM instrumentation. TauContrast and FastFLIM images (for comparison) of three field of views are given. Scale bars, 50 µm. **C** *LT*-Fucci(CA) labeled with MaP555-CA (200 nM) generating a different color variant of the biosensor. The fluorescence lifetimes found for the different cell populations by phasor analysis were 2.9 ns (G1), 2.7 ns (G2/M) and 2.5 ns (S). Due to the lower fluorogenicity of MaP555-CA compared to MaP618-CA there was also a significant contribution of background fluorescence. Scale bars, 50 µm. **D** *In vitro* emission spectra of mCherry, mKate2[5], and MaP618-CA reacted with HaloTag7 or HaloTag9. The spectra of mCherry and mKate2, which compose FUCCI-Red[6], are broader especially toward the far-red region than the spectra of MaP618-CA on HaloTag7 or HaloTag9.

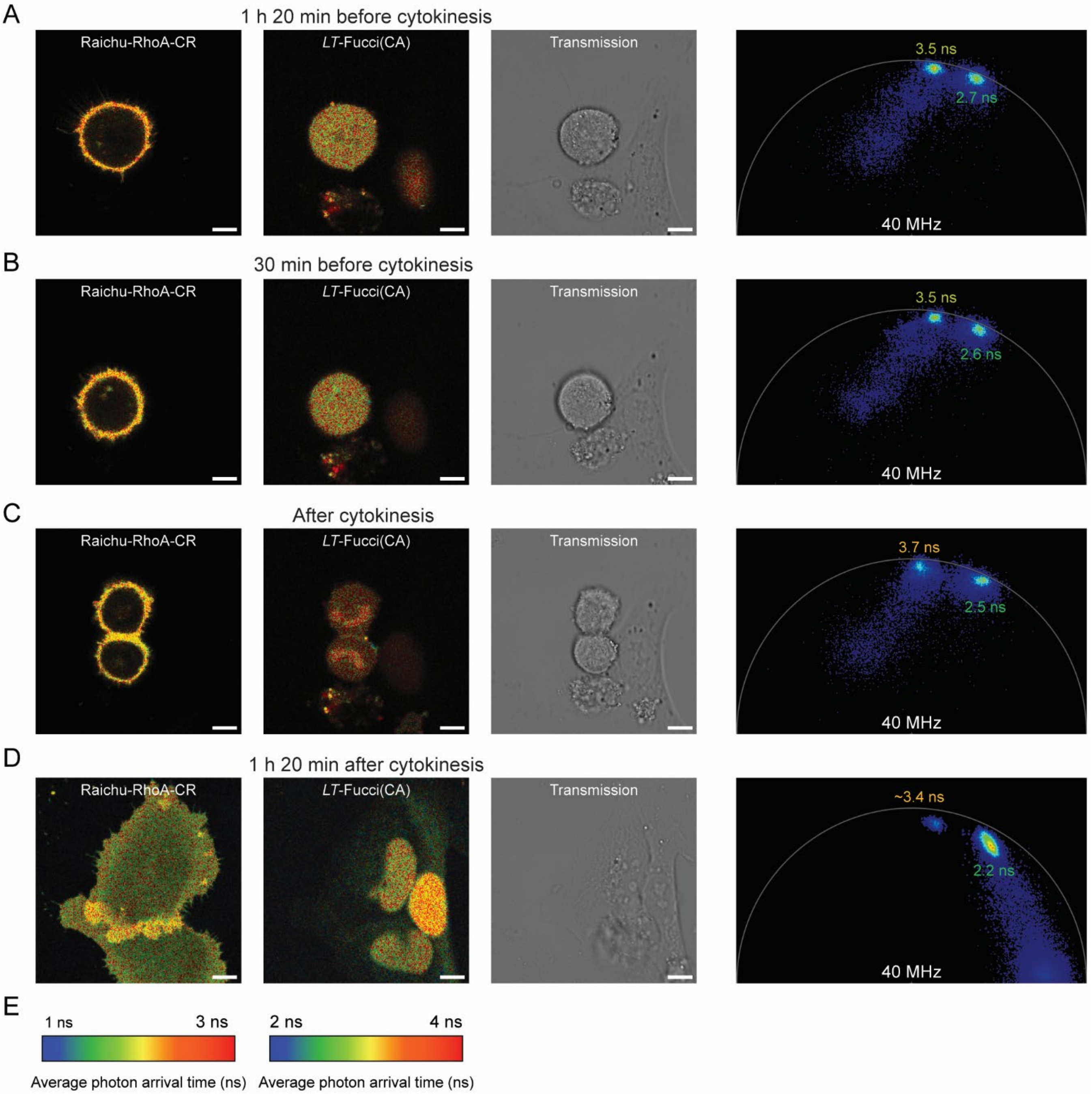

**Supplementary Figure S24:** Multiplexed FLIM of the *LT*-Fucci(CA) and Raichu-RhoA-CR biosensors. **A-D** The activity of the GTPase RhoA as well as the cell cycle stage were followed over time for dividing cells. FLIM images are given for both Raichu-RhoA-CR and *LT*-Fucci(CA) labeled with MaP618-CA (1 µM) along with a transmission light image and the phasor plot 1 h 20 min before cytokinesis (**A**), 30 min before cytokinesis (**B**), shortly after cytokinesis (**C**), and 1 h 20 min after cytokinesis (**D**). The FLIM images are color-coded according to the scales in (**E**). The two biosensors could be easily multiplexed measuring fluorescence lifetime in the GFP channel (Clover-Raichu-RhoA-CR) and the CPY channel (*LT*-Fucci(CA)-MaP618). As expected mitotic cells initially showed low RhoA activity indicated by long donor fluorescence lifetimes (2.7 ns). However, closer to cytokinesis RhoA activity started to increase (2.6 ns) until a donor fluorescence lifetime of 2.5 ns was reached directly after cytokinesis. After division the cells reached even higher RhoA activity (2.2 ns)[7]. Scale bar, 10 µm.

### Supplementary Tables

**Supplementary Table S1:** Summary of HaloTag7 engineering. The number of mutations found that increase the fluorescence intensity of SiR-CA compared to parental HaloTag7 (first round: $I_{Halo\text{-}EGFP}$±3·s.d.$_{Halo}$, second and third round: $I_{Halo\text{-}EGFP}$±2·s.d.$_{Halo}$, $N$ = 5 HaloTag7 samples) are given. The identified hits were picked for sequencing and further analyzed. The mutations are listed together with the ones that were later validated using purified proteins. The second round was started from the five brightest first round variants (M175L, E170K, Q165H, P174H, and P174L) randomizing the remaining three positions individually. Similarly the third round was performed starting from the two brightest second round variants (Q165H-P174R and Q165H-P174L) randomizing the remaining two positions (E170X and M175X) individually.

| | Library | Hits found (picked) | Mutations (redundancy) | Validated |
|---|---|---|---|---|
| First round | M175X | 6 (6) | L(4), M, Q | L |
| | V167X | 6 (5) | V(5) | - |
| | E170X | 1 (1) | K | K, P |
| | T148X | 1 (1) | I | - |
| | L161X | 0 | - | - |
| | Q165X | 1 (1) | H | H |
| | T172X | 0 | - | - |
| | F144X | 0 | - | - |
| | G171X | 0 | - | - |
| | P174X | 4 (4) | H, L, G, Y | H, L, G |
| Second round | E170K, M175X | 2 (2) | M(2) | - |
| | E170X, M175L | 12 (12) | E(5), G, A(2), Q, K, L, V | - |
| | Q165H, M175X | 2 (2) | Y, M | - |
| | Q165X, M175L | 4 (4) | Q, V, L, A | - |
| | Q165H, P174X | 4 (4) | R(2) = HaloTag9, F, L | R, L |
| | Q165X, P174H | 2 (2) | V, Q | - |
| | Q165X, P174L | 1 (1) | V | - |
| | P174X, M175L | 3 (3) | L, H, V | - |
| | P174H, M175X | 0 | - | - |
| | P174L, M175X | 6 (6) | L(3), M(3) | - |
| | Q165X, E170K | 0 | - | - |
| | Q165H, E170X | 2 (2) | V, Q | V |
| | E170K, P174X | 0 | - | - |
| | E170X, P174H | 7 (7) | E(4), K, Q, N | - |
| | E170X, P174L | 4 (4) | E(2), Q, A | - |
| Third round | Q165H, P174R, M175X | 1 (1) | V | - |
| | Q165H, E170X, P174R | 5 (5) | G(2), V, T, A | V, A |
| | Q165H, P174L, M175X | 3 (3) | L, V(2) | - |
| | Q165H, E170X, P174L | 10 (10) | E(5), H, V, G, A, Q | - |

**Supplementary Table S2:** Validation of the engineered variants. SiR changes in fluorescence intensity form the variant-EGFP assay are given compared to the parental variant ($\Delta I_{Par\text{-}EGFP} = (I_{var} - I_{par}) \cdot I_{par}^{-1}$, $N$ = 3 samples) or to HaloTag7 ($\Delta I_{HT7\text{-}EGFP} = (I_{var} - I_{HT7}) \cdot I_{HT7}^{-1}$, $N$ = 3 samples). Selection criteria: $\Delta I_{HT7\text{-}EGFP} > 23$. * Variants selected for further analysis. s.e.m.: standard error of the mean.

| Round | Variant | $\Delta I_{Par\text{-}EGFP}$ [%] | s.e.m. [%] | $\Delta I_{HT7\text{-}EGFP}$ [%] | s.e.m. [%] |
|---|---|---|---|---|---|
| First round | M175L* | - | - | 31 | 7 |
| | E170K* | - | - | 26 | 5 |
| | E170P | - | - | 21 | 5 |
| | Q165H* | - | - | 24 | 5 |
| | P174H* | - | - | 27 | 5 |
| | P174L* | - | - | 43 | 11 |
| | P174G | - | - | 15 | 5 |
| Second round | Q165H-P174R* = HaloTag9 | 32 | 4 | 42 | 7 |
| | Q165H-P174L* | 19 | 3 | 28 | 6 |
| | E170V, Q165H | 10 | 3 | 10 | 3 |
| Third round | Q165H-E170V-P174R | 5 | 3 | 10 | 3 |
| | Q165H-E170A-P174R | 12 | 3 | 17 | 2 |

**Supplementary Table S3:** Fluorophores used for HaloTag9 characterization. Their chemical structures, names, excitation and emission maxima, the settings used during plate reader assays ($Ex_{used}$, $Em_{used}$, and BW = bandwidth), as well as a reference, the vendor, or the distributor are given. R = CA. [a]HaloTag ligand; [b]NHS ester purchases and coupled to $CA\text{-}NH_2$ – see methods.

| # | Structure | Name | $Ex_{max}$ $Ex_{used}$/BW [nm] | $Em_{max}$ $Em_{used}$/BW [nm] | Reference, Vendor or Distributor |
|---|---|---|---|---|---|
| 1 | 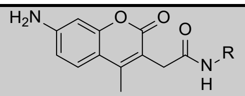 | Coumarin-CA | 353<br>400/25 | 434<br>485/20 | Promega |
| 2 | 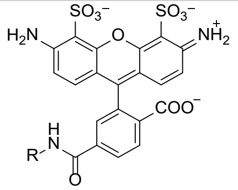 | Alexa 488-CA | 494<br>490/25 | 517<br>595/35 | Promega |
| 3 | 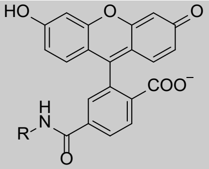 | Fluorescein-CA | 492<br>490/25 | 525<br>595/35 | [8] |
| 4 | 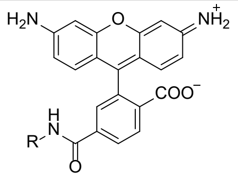 | R110-CA | 502<br>490/25 | 527<br>595/35 | Promega |
| 5 | 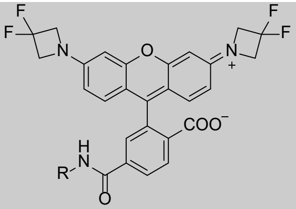 | $JF_{525}$-CA | 525<br>490/25 | 549<br>595/35 | [9] |
| 6 | 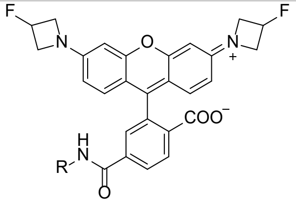 | $JF_{536}$-CA | 536<br>490/25 | 5605<br>95/35 | [9] |
| 7 | 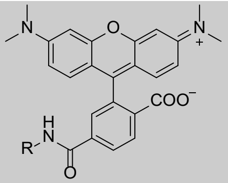 | TMR-CA | 548<br>535/25 | 572<br>595/35 | [10] |
| 8 | 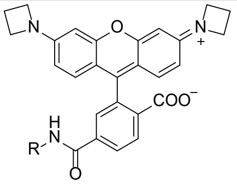 | $JF_{549}$-CA | 549<br>535/25 | 571<br>595/35 | [10] |

| | | | | | |
|---|---|---|---|---|---|
| **9** | 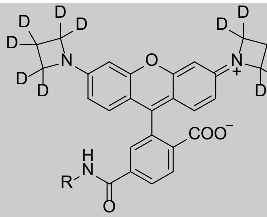 | $JFX_{549}$-CA | 548<br>535/25 | 570<br>595/35 | [11]<br>Janelia |
| **10** | 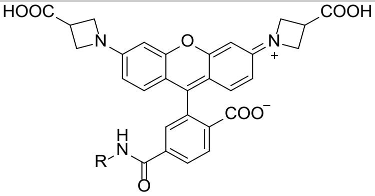 | $JF_{549}$i-CA | 545<br>535/25 | 568<br>595/35 | [9,12]<br>Janelia |
| **11** | 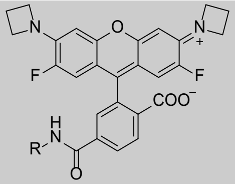 | $JF_{552}$-CA | 552<br>490/25 | 575<br>595/35 | [13]<br>Janelia |
| **12** | 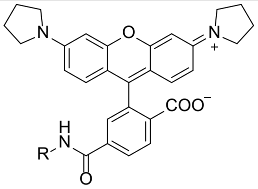 | Rhp-CA | 553<br>535/25 | 576<br>595/35 | [11]<br>Janelia |
| **13** | 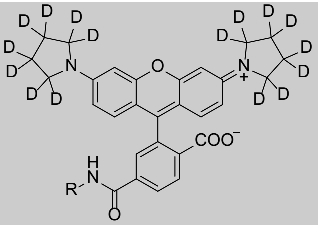 | $JFX_{554}$-CA | 554<br>535/25 | 576<br>595/35 | [11]<br>Janelia |
| **14** | 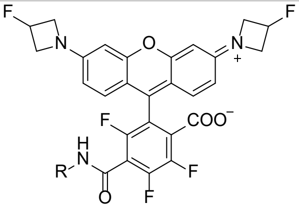 | $JF_{559}$-CA | 559<br>565/25 | 579<br>625/35 | [14]<br>Janelia |
| **15** | 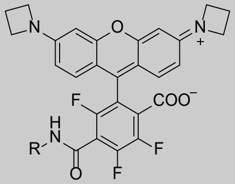 | $JF_{571}$-CA | 571<br>565/25 | 590<br>625/35 | [14]<br>Janelia |
| **16** | 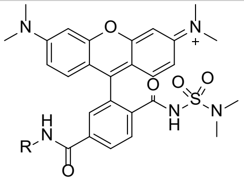 | MaP555-CA | 558[a]<br>535/25 | 578[a]<br>595/35 | [15] |

| | | | | | |
|---|---|---|---|---|---|
| **17** | 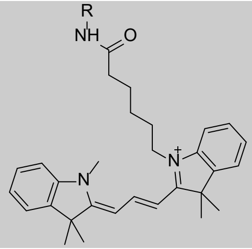 | Cyanine3-CA | 554<br>490/25 | 568<br>595/35 | See methods |
| **18** | 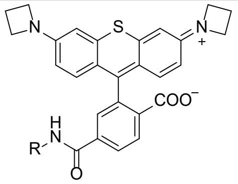 | JF$_{570}$-CA | 570<br>565/25 | 593<br>625/35 | [14]<br>Janelia |
| **19** | 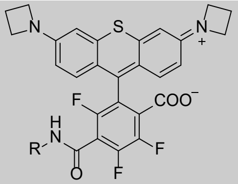 | JF$_{593}$-CA | 593<br>565/25 | 612<br>625/35 | [14]<br>Janelia |
| **20** | 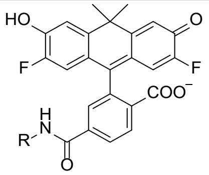 | VO-CA | 555<br>490/25 | 581<br>595/35 | [16]<br>Janelia |
| **21** | 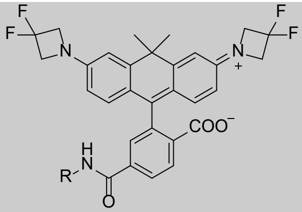 | JF$_{585}$-CA | 585<br>565/25 | 609<br>625/35 | [9] |
| **22** | 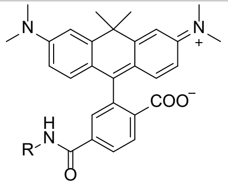 | CPY-CA | 606<br>610/20 | 626<br>680/30 | [17] |
| **23** | 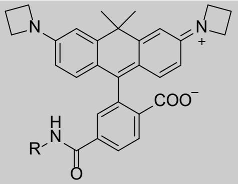 | JF$_{608}$-CA | 608<br>610/20 | 631<br>680/30 | [10] |
| **24** | 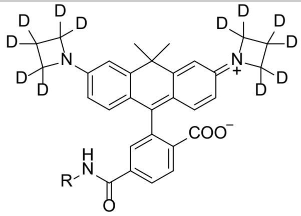 | JFX$_{608}$-CA | 608<br>610/20 | 628<br>680/30 | [11]<br>Janelia |

| | | | | | |
|---|---|---|---|---|---|
| **25** | 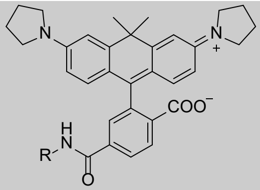 | CRhp-CA | 613<br>610/20 | 633<br>680/30 | [11]<br>Janelia |
| **26** | 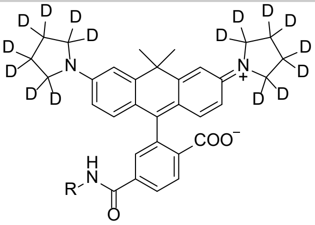 | JFX$_{612}$-CA | 612<br>610/20 | 633<br>680/30 | [11]<br>Janelia |
| **27** | 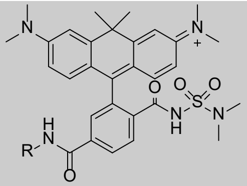 | MaP618-CA | 618[a]<br>610/20 | 635[a]<br>680/30 | [15] |
| **28** | 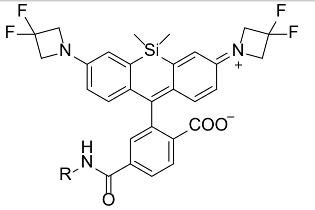 | JF$_{614}$-CA | 622[a]<br>620/20 | 640[a]<br>680/30 | [18] |
| **29** | 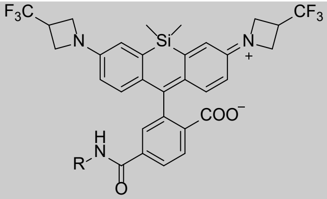 | JF$_{626}$-CA | 634[a]<br>620/20 | 647[a]<br>680/30 | [18]<br>Janelia |
| **30** | 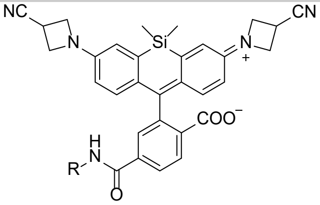 | JF$_{629}$-CA | 638[a]<br>620/20 | 655[a]<br>680/30 | [18]<br>Janelia |
| **31** | 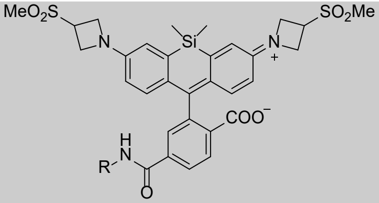 | JF$_{630}$-CA | 633 [a]<br>620/20 | 657[a]<br>680/30 | [18]<br>Janelia |
| **32** | 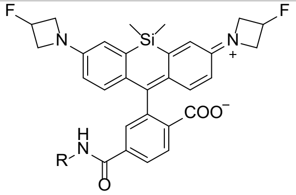 | JF$_{635}$-CA | 635<br>620/20 | 652<br>680/30 | [9] |

| | | | | | |
|---|---|---|---|---|---|
| **33** | 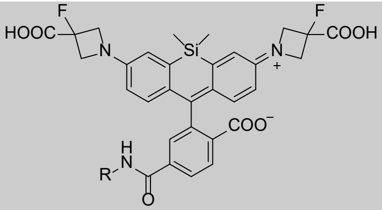 | $JF_{635}$i-CA | 640[a]<br>620/20 | 656[a]<br>680/30 | [19]<br>Janelia |
| **34** | 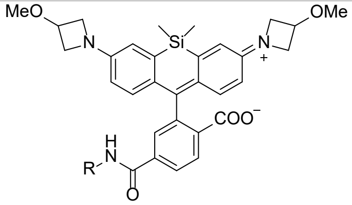 | $JF_{639}$-CA | 645<br>620/20 | 658<br>680/30 | [18]<br>Janelia |
| **35** | 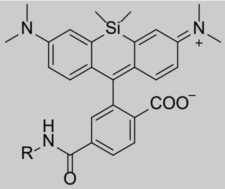 | SiR-CA | 643<br>620/20 | 662<br>680/30 | [20] |
| **36** | 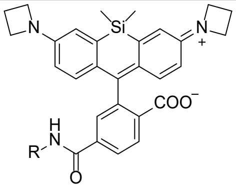 | $JF_{646}$-CA | 646<br>620/20 | 664<br>680/30 | [10] |
| **37** | 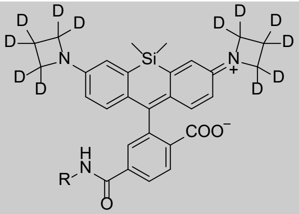 | $JFX_{646}$-CA | 645<br>620/20 | 662<br>680/30 | [11]<br>Janelia |
| **38** | 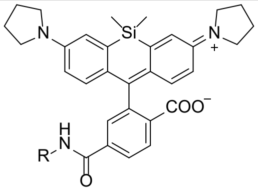 | SiRhp-CA | 652<br>620/20 | 668<br>680/30 | [11]<br>Janelia |
| **39** | 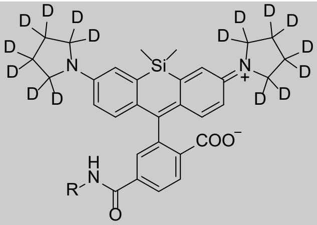 | $JFX_{650}$-CA | 650<br>620/20 | 667<br>680/30 | [11]<br>Janelia |
| **40** | 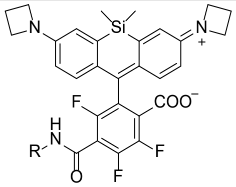 | $JF_{669}$-CA | 669<br>640/30 | 682<br>740/30 | [14]<br>Janelia |

| | | | | | |
|---|---|---|---|---|---|
| **41** | 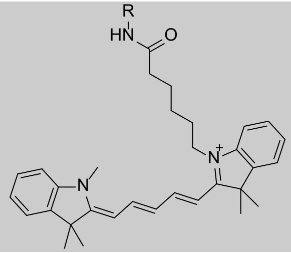 | Cyanine5-CA | 649<br>640/30 | 666<br>740/30 | See methods |
| **42** | 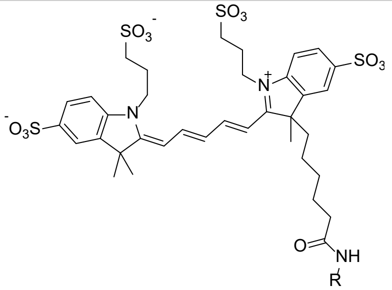 | Alexa 647-CA | 651<br>640/30 | 672<br>740/30 | ThermoFisher[b] |
| **43** | No structure given | Alexa 660-CA | 663<br>640/30 | 690<br>740/30 | Promega |
| **44** | 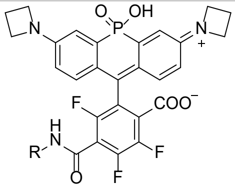 | JF$_{690}$-CA | 690<br>640/30 | 707<br>740/30 | [14]<br>Janelia |
| **45** | 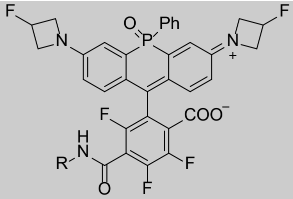 | JF$_{711}$-CA | 711<br>640/30 | 732<br>740/30 | [14]<br>Janelia |
| **46** | 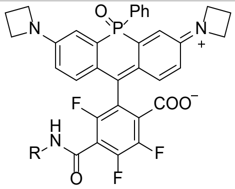 | JF$_{722}$-CA | 722<br>640/30 | 743<br>740/30 | [14]<br>Janelia |

**Supplementary Table S4:** Comparison of the fluorescence intensity of different fluorophores on HaloTag9 relative to HaloTag7. Fluorescence intensity changes are given in % in comparison to HaloTag7 ($\Delta I = (I_{\mathrm{var}} - I_{\mathrm{HT7}}) \cdot I_{\mathrm{HT7}}^{-1}$, mean±s.d., $N$ = 2 replicates each 4 samples unless otherwise stated). $D_{50}$ values are given for selected fluorophores (mean±s.e.m., $N$ = 11 samples). $K_{L\text{-}Z}$ literature values and their reference are given. * not significant (one sided t-test, α = 5%, DF = 6). NA not applicable.

| **Fluorophore** | **Δ*I* [%]** | **s.d. [%]** | **$D_{50}$** | **$K_{L\text{-}Z}$** |
|---|---|---|---|---|
| **Coumarin-CA** | −0.7* | 2.5 | NA | NA |
| **Alexa 488-CA** | −2.7* | 1.5 | | |
| **Fluorescein-CA** | 0.4* | 1.6 | NA | NA |
| **R110-CA** | −1* | 6 | | |
| **JF$_{525}$-CA** | 4.1* | 2.4 | 38.67±0.23 | 0.068 [9] |
| **JF$_{536}$-CA** | −1.2* | 0.3 | 22.3±0.3 | 1.0 [9] |
| **TMR-CA** | 9.7 | 0.6 | 12.5±0.5 | |
| **JF$_{549}$-CA** | −0.2* | 0.5 | | 3.5 [9] |
| **JFX$_{549}$-CA** | −0.1* | 0.4 | | |
| **JF$_{549}$i-CA** | 0.6* | 1.0 | | |
| **JF$_{552}$-CA** | 3.4* | 2.5 | | 0.70 [14] |
| **Rhp-CA** | 2.2* | 0.5 | | |
| **JFX$_{554}$-CA** | 0.1* | 0.5 | | |
| **JF$_{559}$-CA** | 3.6* | 1.5 | | 6.22 [14] |
| **JF$_{571}$-CA** | 1.3* | 1.1 | | 7.93 [14] |
| **MaP555-CA** | 12.2 | 1.7 | 55.8±0.3 | |
| **Cy3-CA** | 40 | 10 | NA | NA |
| **JF$_{570}$-CA** | −0* | 6 | | 2.24 [14] |
| **JF$_{593}$-CA** | −8 | 4 | | 6.06 [14] |
| **VO-CA** | 42 | 6 | NA | NA |
| **JF$_{585}$-CA** | 8.6 | 0.8 | 56.3±1.5 | <0.0001 [9] |
| **CPY-CA** | 11.6 | 0.4 | 35.0±0.5 | |
| **JF$_{608}$-CA** | −3.4* | 1.6 | | 0.091 [9] |
| **JFX$_{608}$-CA** | −5.8* | 0.9 | | |
| **CRhp-CA** | −3.4* | 1.0 | | |
| **JFX$_{612}$-CA** | −7.7* | 1.0 | | |
| **MaP618-CA** | 31 | 4 | 61±7 | |
| **JF$_{614}$-CA** | 102 | 11 | >78 | |
| **JF$_{626}$-CA** | 62 | 14 | | |
| **JF$_{629}$-CA** | 51 | 8 | | |
| **JF$_{630}$-CA** | 20 | 4 | | |
| **JF$_{635}$-CA (*N* = 4)** | 28.4 | 2.5 | >78 | <0.0001 [9] |
| **JF$_{635}$i-CA** | 26.4 | 1.5 | | |
| **JF$_{639}$-CA** | 14 | 6 | | |
| **SiR-CA (*N* = 4)** | 19.9 | 0.5 | 60.5±0.8 | 0.0034 [21] |
| **JF$_{646}$-CA (*N* = 4)** | 7.1* | 0.8 | 55.2±0.3 | 0.0012 [9] |
| **JFX$_{646}$-CA** | 4.5* | 0.3 | | |
| **SiRp-CA** | 4.4* | 2.5 | | |
| **JFX$_{650}$-CA** | 3.8* | 0.6 | | |
| **JF$_{669}$-CA** | 3* | 3 | | 0.262 [14] |
| **Cy5-CA (*N* = 4)** | −3.2* | 1.2 | NA | NA |
| **Alexa 647-CA (*N* = 4)** | 1.9* | 1.2 | NA | NA |
| **Alexa 660-CA (*N* = 4)** | 1.2* | 1.3 | NA | NA |
| **JF$_{690}$-CA** | −0.9* | 0.6 | | 2.90 [14] |
| **JF$_{711}$-CA** | −12.1* | 1.7 | | <0.001 [14] |
| **JF$_{722}$-CA** | 13.3 | 1.0 | | 0.11 [14] |

**Supplementary Table S5:** Characterization of different HaloTag7 variants. Comparison of SiR-CA fluorescence intensity changes of all selected variants without EGFP marker ($\Delta I$, $N$ = 4 replicates each $N$ = 4 samples) as well as the variants' labeling kinetics characterization. The apparent second order rate constant $k_{app}$ measured by either a plate reader or stopped flow, the maximal fluorescence polarization value reached $y_0$, and the dissociation constant $K_D$ are given ($N$ = 3 samples, * $N$ = 2 samples, stop flow $N$ = 3 curves from 14 measurements). In addition, the melting temperature ($N$ = 2 samples) is given. s.d.: standard deviation. ND not determined.

| | Plate reader | | | | | | | | Stop flow | | Thermostability | |
|---|---|---|---|---|---|---|---|---|---|---|---|---|
| **Variant** | **$\Delta I$ [%]** | **s.d. [%]** | **$k_{app}$ [$s^{-1}$ $M^{-1}$]** | **s.d. [$s^{-1}$ $M^{-1}$]** | **$y_0$ [mP]** | **s.d. [mP]** | **$K_D$ [µM]** | **s.d. [µM]** | **$k_{app}$ [$s^{-1}$ $M^{-1}$]** | **s.d. [$s^{-1}$ $M^{-1}$]** | **$T_m$ [°C]** | **s.d. [°C]** |
| **M175L** | 9 | 3 | $8.6 \cdot 10^6$* | $1.7 \cdot 10^6$* | 306.7* | 3* | ND | ND | ND | ND | ND | ND |
| **E170K** | 12 | 5 | $4.4 \cdot 10^6$ | $0.7 \cdot 10^6$ | 287.74 | 0.19 | ND | ND | ND | ND | ND | ND |
| **Q165H** | 10 | 6 | $10 \cdot 10^6$* | $3 \cdot 10^6$* | 273* | 4* | ND | ND | ND | ND | ND | ND |
| **P174H** | 16.8 | 2.0 | $7.7 \cdot 10^6$ | $0.4 \cdot 10^6$ | 331.46 | 0.14 | ND | ND | ND | ND | ND | ND |
| **P174L** | 16 | 5 | $5.70 \cdot 10^6$ | $0.01 \cdot 10^6$ | 288.9 | 0.6 | ND | ND | ND | ND | ND | ND |
| **Q165H-P174L** | 15 | 4 | $9.3 \cdot 10^6$ | $1.1 \cdot 10^6$ | 271.8 | 0.8 | ND | ND | ND | ND | ND | ND |
| **HaloTag9** | 19.9 | 0.5 | $11 \cdot 10^6$ | $5 \cdot 10^6$ | 271 | 3 | 0.123 | 0.024 | $4.4 \cdot 10^7$ | $0.8 \cdot 10^7$ | 60.3 | 0.1 |
| **HaloTag7** | 0 | - | $9.3 \cdot 10^6$ | $0.3 \cdot 10^6$ | 300.3 | 0.7 | 0.23 | 0.07 | $2.4 \cdot 10^7$ | $0.7 \cdot 10^7$ | 61.0 | 0.4 |

**Supplementary Table S6:** Comparison of quantum yields (ϕ) of fluorophores on HaloTag9 relative to HaloTag7. Quantum yields of different fluorophores reacted with HaloTag7, HaloTag9, or in activity buffer are given (mean±s.e.m., *N* = 3 samples). ND not determined. * Low absorbance.

| **Fluorophore** | **HaloTag9 [%]** | | **HaloTag7 [%]** | | **Buffer [%]** | |
|---|---|---|---|---|---|---|
| | **ϕ** | **s.e.m.** | **ϕ** | **s.e.m.** | **ϕ** | **s.e.m.** |
| **$JF_{722}$-CA** | 12.37 | 0.18 | 13.07 | 0.11 | 11.50 | 0.22 |
| **SiR-CA** | 55.3 | 0.5 | 51.7 | 0.3 | 32.6 | 0.5 |
| **$JF_{646}$-CA** | 61.83 | 0.24 | 60.93 | 0.17 | 61* | 3* |
| **$JFX_{646}$-CA** | 72.53 | 0.17 | 70.47 | 0.05 | 62* | 3* |
| **$JF_{635}$-CA** | 67.50 | 0.17 | 67.00 | 0.12 | 53* | 8* |
| **$JF_{635i}$-CA** | 65.57 | 0.11 | 65.83 | 0.05 | 58.5 | 0.5 |
| **$JF_{614}$-CA** | 75.63 | 0.28 | 73.4 | 1.0 | ND | ND |
| **CPY-CA** | 71.67 | 0.22 | 64.17 | 0.23 | 49.4 | 1.2 |
| **$JF_{608}$-CA** | 79.47 | 0.10 | 77.30 | 0.17 | 66.07 | 0.10 |
| **$JFX_{608}$-CA** | 70.97 | 0.10 | 70.07 | 0.19 | 64.4 | 0.3 |
| **MaP618-CA** | 68.07 | 0.24 | 58.50 | 0.09 | 0.7 | 0.6 |
| **$JF_{585}$-CA** | 85.23 | 0.07 | 80.23 | 0.24 | 79.7 | 1.0 |
| **VO-CA** | 75.2 | 0.4 | 71.6 | 0.3 | 39.60 | 0.17 |
| **TMR-CA** | 61.2 | 0.4 | 56.00 | 0.08 | 43.43 | 0.10 |
| **$JF_{549}$-CA** | 86.27 | 0.12 | 84.60 | 0.16 | 84.80 | 0.9 |
| **$JFX_{549}$-CA** | 88.57 | 0.19 | 86.63 | 0.24 | 88.6 | 0.3 |
| **MaP555-CA** | 61.17 | 0.24 | 54.13 | 0.21 | 45* | 5* |
| **$JF_{525}$-CA** | 91.2 | 0.3 | 80.30 | 0.19 | 87.7* | 2.2* |
| **Cy3-CA** | 18.87 | 0.05 | 17.60 | 0.08 | 3.93 | 0.03 |

**Supplementary Table S7:** Comparison of extinction coefficients (ε) of fluorophores on HaloTag9 relative to HaloTag7. Extinction coefficients of different fluorophores reacted with HaloTag7, HaloTag9, or in activity buffer with 0.1% SDS (mean±s.e.m., *N* = 6 samples). ND not determined. *Literature values from references [10,15,18,20].

| Fluorophore | HaloTag9 [$M^{-1}$ $cm^{-1}$] | | HaloTag7 [$M^{-1}$ $cm^{-1}$] | | Buffer+ SDS [$M^{-1}$ $cm^{-1}$] | |
|---|---|---|---|---|---|---|
| | ε | s.e.m. | ε | s.e.m. | ε | s.e.m. |
| **SiR-CA** | 167,000 | 3,000 | 167,300 | 2,200 | 100,000* | 7,000 |
| **$JF_{614}$-CA** | 19,360 | 260 | 7,000* | 400 | ND | ND |
| **CPY-CA** | 124,000 | 800 | 122,000 | 1,000 | 121,000* | 500 |
| **MaP618-CA** | 127,900 | 1,600 | 107,000* | 800 | ND | ND |
| **TMR-CA** | 82,700 | 300 | 81,500 | 500 | 87,000* | 300 |
| **MaP555-CA** | 96,500 | 500 | 94,100 | 600 | 92,000* | 1,100 |

**Supplementary Table S8:** Comparison of fluorescence lifetimes (τ) of fluorophores on HaloTag9 relative to HaloTag7. Fluorescence lifetimes of different fluorophores reacted with HaloTag9 and HaloTag7 as measured by FLIM (mean±s.e.m., *N* = 6-8 FOVs from 3 biological replicates).

| Variant | HaloTag7 [ns] | | | HaloTag9 [ns] | | |
|---|---|---|---|---|---|---|
| | τ | s.e.m. | *N* | τ | s.e.m. | *N* |
| **SiR-CA** | 3.26 | 0.01 | 6 | 3.45 | 0.01 | 6 |
| **$JF_{614}$-CA** | 3.93 | 0.02 | 6 | 3.94 | 0.03 | 8 |
| **CPY-CA** | 3.19 | 0.00 | 8 | 3.66 | 0.02 | 7 |
| **MaP618-CA** | 3.05 | 0.01 | 6 | 3.71 | 0.01 | 7 |
| **TMR-CA** | 2.41 | 0.01 | 6 | 2.80 | 0.01 | 6 |
| **MaP555-CA** | 2.33 | 0.01 | 7 | 2.78 | 0.01 | 6 |

**Supplementary Table S9:** Comparison of fluorophore properties on HaloTag9 and HaloTag7. Changes in fluorescence intensity ($\Delta I = (I_{var} - I_{HT7}) \cdot I_{HT7}^{-1}$, $N$ = 2 replicates each 4 samples, unless otherwise stated) together with the changes in quantum yield ($\Delta\phi = (\phi_{var} - \phi_{HT7}) \cdot \phi_{HT7}^{-1}$, $N$ = 3 samples), changes in extinction coefficient ($\Delta\varepsilon = (\varepsilon_{var} - \varepsilon_{HT7}) \cdot \varepsilon_{HT7}^{-1}$, $N$ = 6 samples), and changes in fluorescence lifetime ($\Delta\tau = (\tau_{var} - \tau_{HT7}) \cdot \tau_{HT7}^{-1}$, $N$ = 6–8 samples). ND not determined, * not significant (one sided t-test, α = 5%, DF = 6, 4, 10, 10-12).

| **Fluorophore** | **Δ*I*±s.d. [%]** | **Δϕ±s.e.m. [%]** | **Δε±s.e.m. [%]** | **Δτ±s.e.m. [%]** |
|---|---|---|---|---|
| **$JF_{722}$-CA** | 13.3±1.0 | −5.4±1.6* | ND | ND |
| **SiR-CA (*N* = 4)** | 19.9±0.5 | 7.0±1.2 | −0.1±2.3* | 5.8±0.5 |
| **$JF_{646}$-CA (*N* = 4)** | 7.1±0.8* | 1.5±0.5* | ND | ND |
| **$JFX_{646}$-CA** | 4.5±0.3* | 2.93±0.25 | ND | ND |
| **$JF_{635}$-CA (*N* = 4)** | 28.4±2.5 | 0.7±0.3* | ND | ND |
| **$JF_{635}$i-CA** | 26.4±1.5 | −0.4±0.4* | ND | ND |
| **$JF_{614}$-CA** | 102±11 | 3.0±1.4* | 177±17 | 0.0±0.8* |
| **CPY-CA** | 11.6±0.4 | 11.7±0.5 | 1.3±1.0* | 14.5±0.5 |
| **$JF_{608}$-CA** | −3.4±1.6* | 2.8±0.3 | ND | ND |
| **$JFX_{608}$-CA** | −5.8±0.9* | 1.3±0.3 | ND | ND |
| **MaP618-CA** | 31±4 | 16.4±0.4 | 19.5±1.8 | 21.4±0.6 |
| **$JF_{585}$-CA** | 8.6±0.8 | 6.2±0.3 | ND | ND |
| **VO-CA** | 42±6 | 5.1±0.6 | ND | ND |
| **TMR-CA** | 9.7±0.6 | 9.2±0.7 | 1.5±0.7* | 16.1±0.8 |
| **$JF_{549}$-CA** | −0.2±0.5* | 1.97±0.24 | ND | ND |
| **$JFX_{549}$-CA** | −0.1±0.4* | 2.2±0.4 | ND | ND |
| **MaP555-CA** | 12.2±1.7 | 13.0±0.6 | 2.6±0.8* | 19.3±0.6 |
| **$JF_{525}$-CA** | 4.1±2.4* | 13.6±0.5 | ND | ND |
| **Cy3-CA** | 40±10 | 7.2±0.6 | ND | ND |

**Supplementary Table S10:** Comparison of the packing interface in HaloTag7-TMR and HaloTag9-TMR. Distances between different amino acid residues and TMR at the interface between two proteins as indicated in Supplementary Fig. S1 and S6. The lengths of the hydrogen bonds $COO^-$–W228(NH) and $COO^-$–S232(OH) as well as the distance from the xanthene core to S232 barley change between HaloTag7 and HaloTag9. The distances between the amino acids W228 and P233 and the xanthene core however changed going from HaloTag7 to HaloTag9. Mean±s.d., $N$ = 2 monomers.

| Variant | $COO^-$–W228(NH) [Å] | $COO^-$–S232(OH) [Å] | Xanthene–W228(Cδ) [Å] | Xanthene–S232(Cα) [Å] | Xanthene–P233(Cδ) [Å] |
|---|---|---|---|---|---|
| **HaloTag9** | 3.0±0.0 | 2.8±0.0 | 3.7±0.0 | 4.3±0.0 | 3.55±0.07 |
| **HaloTag7** | 2.9 | 2.8 | 3.3 | 4.3 | 3.9 |

**Supplementary Table S11:** Structural analysis of HaloTag7-TMR and HaloTag9-TMR. The following parameters are given: the (average) root-mean-square deviation of the alpha carbons of ($RMSD_{\alpha C}$), the (average) root-mean square displacement of the xanthene core ($RMSD_{xanth}$) and the rhodamine excluding the amid bond ($RMSD_{rhod}$), the (average) root-mean square displacement of three residues of the neighboring monomer (W228, S232, and P233) ($RMSD_{pack}$) comparing HaloTag9 with HaloTag7, the (average) dihedral angle between the xanthene and the appended aromatic ring ($\varphi_{Ar\text{-}Ar}$), the (average) tilt angle of the C9–C7'a bond out of the xanthene plane (γ, Supplementary Fig. S8). Mean±s.d., *N* = 2 monomers. [a] Conformation 1. [b] Conformation 2 (Supplementary Fig. S1).

| Variant | $RMSD_{\alpha C}$ [Å] | $RMSD_{xanth}$ [Å] | $RMSD_{rhod}$ [Å] | $RMSD_{pack}$ [Å] | $\varphi_{Ar\text{-}Ar}$ [°] | γ [°] |
|---|---|---|---|---|---|---|
| **HaloTag9** | 0.204±0.015 | 1.82±0.16 | 1.57±0.14 | 0.85±0.13 | 122.32±0.21 | 4.45±0.11 |
| **HaloTag7** | - | - | - | - | 119.0 | 8.7 |
| **6U32** | 0.266 | - | - | - | 104.4[a]<br>100.9[b] | 15.2[a]<br>1.3[b] |
| **TMR open** | - | - | - | - | 94.3 | 5.6 |
| **TMR closed** | - | - | - | - | 115.1 | 37.8 |

**Supplementary Table S12**. Data collection and refinement statistics for the crystal structure of HaloTag9-TMR. Values in parentheses are for the highest resolution shell.

| | **HaloTag7-Q165H-P174R-TMR 6ZVY** |
|---|---|
| **Data collection** | |
| Space group | *P*1 |
| Unit-cell parameters | |
| *a, b, c* (Å) | 44.15, 47.34, 78.47 |
| $\alpha, \beta, \gamma$ (°) | 97.67, 90.19, 113.54 |
| Radiation source | PXII-X10SA, SLS |
| Wavelength (Å) | 0.99988 |
| Temperature (K) | 100 |
| Resolution range (Å) | 50-1.40 (1.50-1.40) |
| No. of observed reflections | 189598 (25609) |
| No. of unique reflections | 99388 (14156) |
| Multiplicity | 1.91 (1.81) |
| Completeness (%) | 87.5 (66.7) |
| $R_{merge}$ (%) | 2.6 (12.5) |
| $<I/\sigma(I)>$ | 17.82 (5.57) |
| $CC_{1/2}$ (%)[#] | 99.9 (97.1) |
| Wilson B ( Å$^2$) | 20.83 |
| **Refinement** | |
| Molecules per a.u. | 2 |
| No. of reflections | 99379 |
| No. of reflections in test set | 4970 |
| Resolution range (Å) | 40.40-1.40 |
| No. of non-hydrogen atoms | |
| Protein | 4730 |
| Ligand/ion | 114 |
| Water | 648 |
| Total | 5492 |
| *R* (%) | 16.21 |
| $R_{free}$ (%) | 18.00 |
| RMS deviations from ideal | |
| bonds (Å) | 0.006 |
| angles (°) | 0.882 |
| *B*-factors (Å$^2$) | |
| Protein | 15.59 |
| Ligand/ion | 16.65 |
| Water | 26.66 |
| Average | 16.92 |
| Ramachandran statistics (%) | |
| favored regions | 96.2 |
| allowed regions | 3.8 |
| disallowed regions | 0 |
| Clashscore | 1.16 |

**Supplementary Table S13:** Brightness comparison of different fluorophores on HaloTag9 relative to HaloTag7 *in cellulo*. Live-cell confocal microscopy was performed on U-2 OS cells expressing HaloTag9 or HaloTag7 co-translationally with EGFP (T2A)[22] labeled with SiR-CA, $JF_{635}$-CA, $JF_{629}$-CA, $JF_{626}$-CA, $JF_{614}$-CA, CPY-CA, MaP618-CA, TMR-CA, and MaP555-CA. Sums of z-stacks were analyzed with regard to their fluorophore and EGFP fluorescence intensities. Mean±s.e.m., *N* = number of total cells analyzed, from three independent preparations.

| Fluorophore | Variant | *N* | Δ*I* [%] | s.e.m. [%] |
|---|---|---|---|---|
| **SiR-CA** | **HaloTag7** | 120 | 0.0 | 0.7 |
| | **HaloTag9** | 120 | −2.6 | 0.6 |
| **$JF_{635}$-CA** | **HaloTag7** | 120 | 0.0 | 1.2 |
| | **HaloTag9** | 120 | 12.1 | 1.2 |
| **$JF_{629}$-CA** | **HaloTag7** | 120 | 0.0 | 1.4 |
| | **HaloTag9** | 120 | 9.3 | 1.5 |
| **$JF_{626}$-CA** | **HaloTag7** | 120 | 0.0 | 1.5 |
| | **HaloTag9** | 123 | 7.1 | 1.3 |
| **$JF_{614}$-CA** | **HaloTag7** | 120 | 0.0 | 0.8 |
| | **HaloTag9** | 120 | 28.0 | 1.1 |
| **CPY-CA** | **HaloTag7** | 117 | 0.0 | 0.7 |
| | **HaloTag9** | 120 | 3.6 | 0.6 |
| **MaP618-CA** | **HaloTag7** | 120 | 0.0 | 0.7 |
| | **HaloTag9** | 119 | 39.2 | 0.9 |
| **TMR-CA** | **HaloTag7** | 122 | 0.0 | 1.1 |
| | **HaloTag9** | 120 | 17.7 | 1.3 |
| **MaP555-CA** | **HaloTag7** | 119 | 0.2 | 1.2 |
| | **HaloTag9** | 129 | 17.4 | 1.0 |

**Supplementary Table S14:** Plasmids used and generated in this work as well as the stable cell lines derived thereof. * Plasmid published in [23].

| Name | Addgene# | Plasmid | Gene | Entry Plasmid(s) Addgene# | Stable cell lines |
|---|---|---|---|---|---|
| **pET51b(+)_HaloTag9-EGFP** | - | pET51b(+) | HaloTag9 | pET51b(+)_HaloTag7 | - |
| **pET51b(+)_HaloTag9** | 169324 | pET51b(+) | HaloTag9 | pET51b(+)_HaloTag7 | - |
| **pCDNA5/FRT/TO_HaloTag7_T2A_EGFP** | 169325 | pCDNA5/FRT/TO | HaloTag7 and EGFP | 135444[23] | U-2 OS Flp-In TREx |
| **pCDNA5/FRT/TO_HaloTag9_T2A_EGFP** | 169326 | pCDNA5/FRT/TO | HaloTag9 and EGFP | 135444[23] | U-2 OS Flp-In TREx |
| **pCDNA5/FRT/TO_CEP41-HaloTag7_T2A_EGFP** | 169327 | pCDNA5/FRT/TO | CEP41-HaloTag7 and EGFP | 135446[23] | - |
| **pCDNA5/FRT/TO_CEP41-HaloTag9_T2A_EGFP** | 169328 | pCDNA5/FRT/TO | CEP41-HaloTag9 and EGFP | 135446[23] | - |
| **pCDNA5/FRT/TO_H2B-HaloTag7_T2A_EGFP*** | - | pCDNA5/FRT/TO | H2B-HaloTag9 and EGFP | - | U-2 OS Flp-In TREx |
| **pCDNA5/FRT/TO_H2B-HaloTag9_T2A_EGFP** | 169332 | pCDNA5/FRT/TO | H2B-HaloTag9 and EGFP | 135444[23] | U-2 OS Flp-In TREx |
| **pCDNA5/FRT/TO_TOMM20-HaloTag7_T2A_EGFP*** | - | pCDNA5/FRT/TO | TOMM20-HaloTag9 and EGFP | - | - |
| **pCDNA5/FRT/TO_TOMM20-HaloTag9_T2A_EGFP** | 169333 | pCDNA5/FRT/TO | TOMM20-HaloTag9 and EGFP | 135443[23] | - |
| **pCDNA5/FRT/TO_CEP41-HaloTag7*** | - | pCDNA5/FRT/TO | CEP41-HaloTag7 | - | U-2 OS Flp-In TREx |
| **pCDNA5/FRT/TO_CEP41-HaloTag9** | 169331 | pCDNA5/FRT/TO | CEP41-HaloTag9 | 135446[23] | U-2 OS Flp-In TREx |
| **pCDNA5/FRT/TO_H2B-HaloTag7** | 169329 | pCDNA5/FRT/TO | H2B-HaloTag7 | 135444[23] | U-2 OS Flp-In TREx |
| **pCDNA5/FRT/TO_H2B-HaloTag9** | 169334 | pCDNA5/FRT/TO | H2B-HaloTag9 | 135444[23] | U-2 OS Flp-In TREx |
| **pCDNA5/FRT/TO_TOMM20-HaloTag7** | 169330 | pCDNA5/FRT/TO | TOMM20-HaloTag7 | 135443[23] | U-2 OS Flp-In TREx |
| **pCDNA5/FRT/TO_TOMM20-HaloTag9** | 169335 | pCDNA5/FRT/TO | TOMM20-HaloTag9 | 135443[23] | U-2 OS Flp-In TREx |
| **pCDNA5/FRT_Fucci(CA)** | 169338 | pCDNA5/FRT | HaloTag7-Geminin(1/110) and HaloTag9-Cdt(1-100)Cy- | 83841[24] 80007[25] | U-2 OS Flp-In TREx |
| **pCDNA5/FRT_Fucci(SA)** | 169336 | pCDNA5/FRT | HaloTag7-Geminin(1/110) and HaloTag9-Cdt(30-120) | 83841[24] | U-2 OS Flp-In TREx |
| **pCDNA5/FRT_Fucci(SCA)** | 169337 | pCDNA5/FRT | HaloTag7-Geminin(1/110) and HaloTag9-Cdt(1-100)Cy+ | 83841[24] 80007[25] | U-2 OS Flp-In TREx |
| **pCAGGS-Raichu-RhoA-CR** | - | pCAGGS | Raichu-RhoA-CR | 40258[26] | - |

**Supplementary Table S15:** Fluorescence microscopy data acquisition parameters. Fluorophores refer to CA analogues unless otherwise stated. *See methods for details.

| Image | Label | Ligand | Microscope | Excitation [nm] | Pixel dwell time [µs] | Pinhole [Airy Units] | Objective | Pixel size [nm] | Size [pixels] | Emission [nm] | Comment |
|---|---|---|---|---|---|---|---|---|---|---|---|
| **Fig 1D** | HT7 + HT9 | SiR-CA | SP8-FALCON | 631 | 2.09 | 1 | 40x1.10 water | 569 | 512x512 | 680-700 | 80 MHz |
| **Fig 1D** | HT7 + HT9 | $JF_{614}$-CA | SP8-FALCON | 605 | 2.09 | 1 | 40x1.10 water | 569 | 512x512 | 640-700 | 80 MHz |
| **Fig 1D** | HT7 + HT9 | CPY-CA | SP8-FALCON | 595 | 2.09 | 1 | 40x1.10 water | 569 | 512x512 | 640-660 | 80 MHz |
| **Fig 1D** | HT7 + HT9 | MaP618-CA | SP8-FALCON | 595 | 2.09 | 1 | 40x1.10 water | 569 | 512x512 | 640-660 | 80 MHz |
| **Fig 1D** | HT7 + HT9 | TMR-CA | SP8-FALCON | 550 | 2.09 | 1 | 40x1.10 water | 569 | 512x512 | 570-600 | 80 MHz |
| **Fig 1D** | HT7 + HT9 | MaP555-CA | SP8-FALCON | 550 | 2.09 | 1 | 40x1.10 water | 569 | 512x512 | 570-600 | 80 MHz |
| **Fig 2A** | HT7 + HT9 | MaP618-CA | SP8 | 595 | 1.58 | 1 | 20x0.75 dry | 569 | 1024x1024 | 640-660 | Sum projection |
| **Fig 2A** | EGFP | - | SP8 | 489 | 1.58 | 1 | 20x0.75 dry | 569 | 1024x1024 | 510-530 | Sum projection |
| **Fig 2B** | HT7 or HT9 | MaP618-CA | SP8-FALCON | 615 | - | 1 | 40x1.10 water | - | - | 630-700 | FCS |
| **Fig 2C** | Cep41-HT7 | MaP618-CA | SP8 | 595 | 10.36 | 1 | 40x1.10 water | 110 | 760x760 | 640-660 | Sum projection |
| **Fig 2C** | EGFP | - | SP8 | 489 | 10.36 | 1 | 40x1.10 water | 110 | 760x760 | 510-530 | Sum projection |
| **Fig 2D** | Cep41-HT9 | MaP618-CA | SP8 | 595 | 10.36 | 1 | 40x1.10 water | 110 | 760x760 | 640-660 | Sum projection |
| **Fig 2D** | EGFP | - | SP8 | 489 | 10.36 | 1 | 40x1.10 water | 110 | 760x760 | 510-530 | Sum projection |
| **Fig 2F** | H2B-HT7 and Tomm20-HT9 | MaP555-CA MaP555-Actin | SP8-FALCON | 550 | 13.68 | 1 | 40x1.10 water | 101 | 576x576 | 570-620 | 80 MHz 10 line accumu. |
| **Fig 2H-I** | *LT*-Fucci(CA) | MaP618-CA | SP8-FALCON | 615 | 1.58 | 5 | 40x1.10 water | 284 | 1024x1024 | 640-700 | 40 MHz 4 line accumu. 2 pixel binning 24 h every 12 min |
| **S10A** | HT7 or HT9 | SiR-CA | SP8 | 631 | 1.58 | 1 | 20x0.75 dry | 569 | 1024x1024 | 680-700 | Sum projection |
| **S10A** | EGFP | - | SP8 | 489 | 1.58 | 1 | 20x0.75 dry | 569 | 1024x1024 | 510-530 | Sum projection |
| **S10B** | HT7 or HT9 | $JF_{635}$-CA | SP8 | 605 | 1.58 | 1 | 20x0.75 dry | 569 | 1024x1024 | 640-700 | Sum projection |
| **S10B** | EGFP | - | SP8 | 489 | 1.58 | 1 | 20x0.75 dry | 569 | 1024x1024 | 510-530 | Sum projection |
| **S10C** | HT7 or HT9 | $JF_{629}$-CA | SP8 | 605 | 1.58 | 1 | 20x0.75 dry | 569 | 1024x1024 | 640-700 | Sum projection |
| **S10C** | EGFP | - | SP8 | 489 | 1.58 | 1 | 20x0.75 dry | 569 | 1024x1024 | 510-530 | Sum projection |

| | | | | | | | | | | | |
|---|---|---|---|---|---|---|---|---|---|---|---|
| **S10D** | HT7 or HT9 | $JF_{626}$-CA | SP8 | 605 | 1.58 | 1 | 20x0.75 dry | 569 | 1024x1024 | 640-700 | Sum projection |
| **S10D** | EGFP | - | SP8 | 489 | 1.58 | 1 | 20x0.75 dry | 569 | 1024x1024 | 510-530 | Sum projection |
| **S10E** | HT7 or HT9 | $JF_{614}$-CA | SP8 | 605 | 1.58 | 1 | 20x0.75 dry | 569 | 1024x1024 | 640-700 | Sum projection |
| **S10E** | EGFP | - | SP8 | 489 | 1.58 | 1 | 20x0.75 dry | 569 | 1024x1024 | 510-530 | Sum projection |
| **S10F** | HT7 or HT9 | CPY-CA | SP8 | 595 | 1.58 | 1 | 20x0.75 dry | 569 | 1024x1024 | 640-660 | Sum projection |
| **S10F** | EGFP | - | SP8 | 489 | 1.58 | 1 | 20x0.75 dry | 569 | 1024x1024 | 510-530 | Sum projection |
| **S10G** | HT7 or HT9 | MaP618-CA | SP8 | 595 | 1.58 | 1 | 20x0.75 dry | 569 | 1024x1024 | 640-660 | Sum projection |
| **S10G** | EGFP | - | SP8 | 489 | 1.58 | 1 | 20x0.75 dry | 569 | 1024x1024 | 510-530 | Sum projection |
| **S10H** | HT7 or HT9 | TMR-CA | SP8 | 550 | 1.58 | 1 | 20x0.75 dry | 569 | 1024x1024 | 570-600 | Sum projection |
| **S10H** | EGFP | - | SP8 | 489 | 1.58 | 1 | 20x0.75 dry | 569 | 1024x1024 | 510-530 | Sum projection |
| **S10I** | HT7 or HT9 | MaP555-CA | SP8 | 550 | 1.58 | 1 | 20x0.75 dry | 569 | 1024x1024 | 570-600 | Sum projection |
| **S10I** | EGFP | - | SP8 | 489 | 1.58 | 1 | 20x0.75 dry | 569 | 1024x1024 | 510-530 | Sum projection |
| **S10J** | HT7 or HT9 | $JF_{525}$-CA | SP8 | 525 | 1.58 | 1 | 20x0.75 dry | 569 | 1024x1024 | 560-580 | Sum projection |
| **S10J** | EGFP | - | SP8 | 489 | 1.58 | 1 | 20x0.75 dry | 569 | 1024x1024 | 505-515 | Sum projection |
| **S11A** | HT7 or HT9 | SiR-CA | SP8-FALCON | 630 | - | 1 | 40x1.10 water | - | - | 650-700 | FCS |
| **S11B** | HT7 or HT9 | $JF_{614}$-CA | SP8-FALCON | 615 | - | 1 | 40x1.10 water | - | - | 630-700 | FCS |
| **S11C** | HT7 or HT9 | MaP618-CA | SP8-FALCON | 615 | - | 1 | 40x1.10 water | - | - | 630-700 | FCS |
| **S12A** | H2B-HT7 or H2B-HT9 | SiR-CA | SP8 | 630 | 0.86 | 5 | 40x1.10 water | 284 | 1024x1024 | 650-700 | Sum projection Bleaching |
| **S12B** | H2B-HT7 or H2B-HT9 | $JF_{614}$-CA | SP8 | 614 | 0.86 | 5 | 40x1.10 water | 284 | 1024x1024 | 630-750 | Sum projection Bleaching |
| **S12C** | H2B-HT7 or H2B-HT9 | CPY-CA | SP8 | 614 | 0.86 | 5 | 40x1.10 water | 284 | 1024x1024 | 630-750 | Sum projection Bleaching |
| **S12D** | H2B-HT7 or H2B-HT9 | MaP618-CA | SP8 | 614 | 0.86 | 5 | 40x1.10 water | 284 | 1024x1024 | 630-750 | Sum projection Bleaching |
| **S12E** | H2B-HT7 or H2B-HT9 | TMR-CA | SP8 | 555 | 0.86 | 5 | 40x1.10 water | 284 | 1024x1024 | 570-620 | Sum projection Bleaching |
| **S12F** | H2B-HT7 or H2B-HT9 | MaP555-CA | SP8 | 555 | 0.86 | 5 | 40x1.10 water | 284 | 1024x1024 | 570-620 | Sum projection Bleaching |
| **S13A** | Tomm20-HT7 or Tomm20-HT9 | SiR-CA | Abberior | 640 | 7.00 | 1 | 100x1.40 oil | 25 | 680x680 | 650-725 | 2 line accumu. STED 3 frame accumu. |
| **S13B** | Tomm20-HT7 or Tomm20-HT9 | MaP618-CA | Abberior | 640 | 7.00 | 1 | 100x1.40 oil | 25 | 680x680 | 650-725 | 2 line accumu. STED 3 frame accumu. |
| **S14A** | Tomm20-HT7 | MaP618-CA | Abberior | 640 | 7.00 | 1 | 100x1.40 oil | 25 | 680x680 | 650-725 | 2 line accumu. STED 3 frame accumu. |

| | | | | | | | | | | | |
|---|---|---|---|---|---|---|---|---|---|---|---|
| **S14B** | Tomm20-HT9 | MaP618-CA | Abberior | 640 | 7.00 | 1 | 100x1.40 oil | 25 | 680x680 | 650-725 | 2 line accumu. STED 3 frame accumu. |
| **S14C** | Cep41-HT7 | MaP618-CA | Abberior | 640 | 12.00 | 1 | 100x1.40 oil | 25 | 680x680 | 650-725 | 2 line accumu. STED 3 frame accumu. |
| **S14D** | Cep41-HT9 | MaP618-CA | Abberior | 640 | 12.00 | 1 | 100x1.40 oil | 25 | 680x680 | 650-725 | 2 line accumu. STED 3 frame accumu. |
| **S15A** | Tomm20-HT7 and H2B-HT9 | MaP618-CA | SP8-FALCON | 615 | 22.90 | 1 | 40x1.10 water | 113 | 344x344 | 630-700 | 80 MHz 10 line accumu. |
| **S15B** | H2B-HT7 and Tomm20-HT9 | MaP618-CA | SP8-FALCON | 615 | 21.40 | 1 | 40x1.10 water | 112 | 520x520 | 630-700 | 80 MHz 10 line accumu. |
| **S15C** | Cep41-HT7 and H2B-HT9 | MaP618-CA | SP8-FALCON | 615 | 10.70 | 1 | 40x1.10 water | 112 | 368x368 | 630-700 | 80 MHz 10 line accumu. |
| **S15D** | Cep41-HT9 and H2B-HT7 | MaP618-CA | SP8-FALCON | 615 | 8.06 | 1 | 40x1.10 water | 113 | 488x488 | 630-700 | 80 MHz 6 line accumu. |
| **S15E** | Cep41-HT7 and Tomm20-HT9 | MaP618-CA | SP8-FALCON | 615 | 7.58 | 1 | 40x1.10 water | 112 | 448x448 | 630-700 | 80 MHz 10 line accumu. |
| **S15F** | Cep41-HT7 and Tomm20-HT9 | MaP618-CA | SP8-FALCON | 615 | 8.79 | 1 | 40x1.10 water | 112 | 520x520 | 630-700 | 80 MHz 10 line accumu. |
| **S16A** | Tomm20-HT7 and H2B-HT9 | MaP618-CA | SP8-FALCON | 615 | 22.90 | 1 | 40x1.10 water | 113 | 344x344 | 630-700 | 40 MHz 10 line accumu. |
| **S16B** | H2B-HT7 and Tomm20-HT9 | MaP618-CA | SP8-FALCON | 615 | 21.40 | 1 | 40x1.10 water | 112 | 520x520 | 630-700 | 40 MHz 10 line accumu. |
| **S17A** | Tomm20-HT7 and H2B-HT9 | MaP555-CA | SP8-FALCON | 550 | 21.89 | 1 | 40x1.10 water | 101 | 360x360 | 570-620 | 80 MHz 10 line accumu. |
| **S17B** | H2B-HT7 and Tomm20-HT9 | MaP555-CA | SP8-FALCON | 550 | 17.59 | 1 | 40x1.10 water | 100 | 448x448 | 570-620 | 80 MHz 10 line accumu. |
| **S17C** | Cep41-HT7 and H2B-HT9 | MaP555-CA | SP8-FALCON | 550 | 7.94 | 1 | 40x1.10 water | 101 | 496x496 | 570-620 | 80 MHz 10 line accumu. |
| **S17D** | Cep41-HT9 and H2B-HT7 | MaP555-CA | SP8-FALCON | 550 | 6.92 | 1 | 40x1.10 water | 100 | 568x568 | 570-620 | 80 MHz 10 line accumu. |
| **S18A** | H2B-HT7 and Tomm20-HT9 | MaP555-CA | SP8-FALCON | 550 | 4.55 | 1 | 40x1.10 water | 101 | 864x864 | 570-620 | 80 MHz 12 line accumu. |
| **S18B** | H2B-HT7 and Tomm20-HT9 | MaP618-CA | SP8-FALCON | 615 | 8.64 | 1 | 40x1.10 water | 112 | 456x456 | 630-700 | 80 MHz 10 line accumu. |

| | | | | | | | | | | | |
|---|---|---|---|---|---|---|---|---|---|---|---|
| **S19A** | H2B-HT7 and Tomm20-HT9 | MaP555-CA MaP555-Actin | SP8-FALCON | 550 | 10.94 | 1 | 40x1.10 water | 101 | 720x720 | 570-620 | 80 MHz<br>10 line accumu. |
| **S19B** | H2B-HT7 and Tomm20-HT9 | MaP555-CA MaP555-Actin | SP8-FALCON | 550 | 13.68 | 1 | 40x1.10 water | 101 | 576x576 | 570-620 | 80 MHz<br>10 line accumu. |
| **S19C** | H2B-HT7 and Tomm20-HT9 | MaP618-CA MaP618-Actin | SP8-FALCON | 615 | 15.15 | 1 | 40x1.10 water | 112 | 520x520 | 630-700 | 80 MHz<br>10 line accumu. |
| **S19D** | H2B-HT7 and Tomm20-HT9 | MaP555-CA MaP555-Tubulin | SP8-FALCON | 550 | 6.39 | 1 | 40x1.10 water | 101 | 616x616 | 570-620 | 80 MHz<br>10 line accumu. |
| **S19E** | Tomm20-HT9 | MaP555-CA MaP555-Tubulin | SP8-FALCON | 550 | 8.64 | 1 | 40x1.10 water | 100 | 456x456 | 570-620 | 80 MHz<br>10 line accumu. |
| **S19F** | Cep41-HT7 and Tomm20-HT9 | MaP555-CA MaP555-DNA | SP8-FALCON | 550 | 6.31 | 1 | 40x1.10 water | 102 | 624x624 | 570-620 | 80 MHz<br>10 line accumu. |
| **S20A** | HT7 and HT9 | MaP618-CA | SP8 | * | 1.58 | 1 | 40x1.10 water | 284 | 1024x1024 | * | Spectra |
| **S20A** | - | MaP618-Actin | SP8 | * | 1.29 | 1 | 40x1.10 water | 116 | 1256x1256 | * | Spectra |
| **S20B** | HT7 and HT9 | MaP555-CA | SP8 | * | 1.58 | 1 | 40x1.10 water | 284 | 1024x1024 | * | Spectra |
| **S20B** | - | MaP555-Actin | SP8 | * | 1.29 | 1 | 40x1.10 water | 116 | 1256x1256 | * | Spectra |
| **S20B** | - | MaP555-Tubulin | SP8 | * | 3.84 | 1 | 40x1.10 water | 284 | 1024x1024 | * | Spectra<br>2 line accumu. |
| **S20B** | - | MaP555-DNA | SP8 | * | 1.58 | 1 | 40x1.10 water | 284 | 1024x1024 | * | Spectra |
| **S21** | H2B-HT7 and Tomm20-HT9 | MaP618-CA | SELLARIS 8 STED FALCON | 615 | 1.20 | 1 | 86x1.20 water | 20 | 1792x1792 | 630-760 | 80 MHz<br>10 line accumu.<br>2 pixel bin (confocal)<br>STED |
| **S22A** | *LT*-Fucci(CA) | MaP618-CA | SP8-FALCON | 615 | 3.88 | 1 | 40x1.10 water | 284 | 1024x1024 | 640-700 | 40 MHz<br>4 line accumu.<br>2 pixel binning |
| **S22B** | *LT*-Fucci(CA) | MaP618-CA | SP8-FALCON | 615 | 1.58 | 5 | 40x1.10 water | 284 | 1024x1024 | 640-700 | 40 MHz<br>4 line accumu.<br>2 pixel binning<br>24 h every 12 min |

| | | | | | | | | | | | |
|---|---|---|---|---|---|---|---|---|---|---|---|
| **S22C** | *LT*-Fucci(SA) | MaP618-CA | SP8-FALCON | 615 | 1.58 | 5 | 40x1.10 water | 284 | 1024x1024 | 640-700 | 40 MHz<br>4 line accumu.<br>2 pixel binning<br>24 h every 12 min |
| **S22D** | *LT*-Fucci(SCA) | MaP618-CA | SP8-FALCON | 615 | 1.58 | 5 | 40x1.10 water | 284 | 1024x1024 | 640-700 | 40 MHz<br>4 line accumu.<br>2 pixel binning<br>24 h every 12 min |
| **S23A** | *LT*-Fucci(CA) | MaP618-CA | SP8-FALCON | 615 | 1.58 | 5 | 40x1.10 water | 284 | 1024x1024 | 640-700 | 40 MHz<br>4 line accumu.<br>2 pixel binning<br>24 h every 12 min |
| **S23B** | *LT*-Fucci(CA) | MaP618-CA | SELLARIS 8 (FALCON) | 615 | 1.58 | 1 | 86x1.20 water | 284 | 1024x1024 | 630-700 | 80 MHz<br>10 line accumu.<br>TauContrast |
| **S23C** | *LT*-Fucci(CA) | MaP555-CA | SP8-FALCON | 550 | 7.69 | 5 | 40x1.10 water | 569 | 512x512 | 570-620 | 40 MHz<br>10 line accumu. |
| **S24A-D** | *LT*-Fucci(CA)<br>Raichu-RhoA-CR | MaP618-CA<br>- | SP8-FALCON | 615<br>495 | 7.69 | 1 | 40x1.10 water | 162 | 512x512 | 640-700<br>510-560 | 40 MHz<br>10 line accumu.<br>2 pixel binning<br>2.5 h every 5 min |

### Supplementary Videos

#### Supplementary Videos 1-3. *LT*-Fucci biosensors over 24 h in living U-2 OS cells.

FastFlim video of U-2 OS cells stably expressing *LT*-Fucci(CA) (**1**), *LT*-Fucci(SA) (**2**), or *LT*-Fucci(SCA) (**3**) after no-wash labeling with MaP618-CA (1 µM). For all biosensors HaloTag9-hCdt is present during G1 and the nuclei will therefore present long average photon arrival times (MaP618-CA: 3.7 ns, orange). During S, G2 and M phase HaloTag7-hGem or a mixture of HaloTag7-hGem and HaloTag9-hCdt is present, resulting in short average photon arrival times (MaP618-CA: 3.1 ns, green) or a gradient of medium average photon arrival times (MaP618-CA: ~3.4 ns, light-green). Scale bar, 50 µm.

### Supplementary Methods

**General considerations:** Fluorophores-CA were either prepared according to literature procedures by B. Matthes or D. Schmidt (MPI-MR), kindly provided by Dr. L. Lavis (Janelia Research Campus) or purchased from commercial vendors (Supplementary Table S3). The synthesis of Cyanine3-CA, Cyanine5-CA, and Alexa647-CA are described below. MaP618-Actin, MaP555-Actin, MaP555-DNA, and MaP555-Tubulin were purchased from Spirochrome. Fluorophores were prepared as stock solutions in dry DMSO and diluted in the respective buffer such that the final concentration of DMSO did not exceed 1% v/v. Activity buffer (50 mM HEPES, 150 mM NaCl, pH 7.2) was used in all experiments unless otherwise stated. 96-well plates (black, flat bottom, non-binding, (Corning)) were used unless otherwise stated. Fluorescence intensity was measured on a plate reader (Spark® 20M, Tecan) equipped with filters and a monochromator. Excitation and emission collection was performed as indicated in Supplementary Table S3. EGFP was excited at 485/20 nm and emission was collected at 535/25 nm.

**Synthesis:** All chemical reagents and anhydrous solvents for synthesis were purchased from commercial suppliers (Acros, Merck, Sigma-Aldrich, and TCI) and used without further purification. CA-NHBoc was synthesized according to literature procedures[8]. Alexa647-NHS was purchased from ThermoFisher. Composition of mixed solvents is given by volume ratio (v/v). High-resolution mass spectrometry (HRMS) was performed by the MS-facility of the Max Planck Institute for Medical Research on a Bruker maXis II™ ETD. Liquid chromatography coupled to mass spectrometry (LC-MS) was performed on a Shimadzu MS2020 connected to a Nexera UHPLC system equipped with a Supelco Titan C18 80 Å (1.9 µm, 2.1 x 50 mm). Buffer A: 0.05% HCOOH in $H_2O$ Buffer B: 0.05% HCOOH in ACN. Analytical gradient was from 10% to 90% B within 6 min with 0.5 mL min$^{-1}$ flow. Preparative reverse phase high-performance liquid chromatography (RP-HPLC) was either carried out on a Dionex system equipped with an UltiMate 3000 diode array detector for product visualization on a Supleco Ascentis® C18 column (5 µm, 10 x 250 mm) or on a Supleco Ascentis® C18 column (5 µm, 21.2 x 250 mm). Buffer A: 0.1% TFA in $H_2O$ Buffer B: ACN. Typical gradient was from 10% to 90% B within 32 min with 4 or 8 mL min$^{-1}$ flow or on a Shimadzu system equipped with an SPD-M20A diode array detector for product visualization and a LCMS-2020 for mass detection on a Shimadzu Shim-pack GIS C18 column (5 µm, 50 x 250 mm). Buffer A: 0.1% FA in $H_2O$ Buffer B: 0.1% FA in ACN. Typical gradient was from 10% to 90% B within 60 min with 50 mL min$^{-1}$ flow.

**Cyanine3-CA**

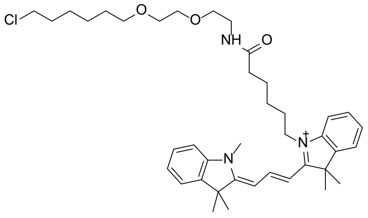

Cyanine3-CA

Cyanine3-COOH was synthesized according to Ueno et al. 2011.[27]

A solution of Cyanine3-COOH (50 mg, 219 µmol, 1.0 equiv.) in dry DMSO (1.0 mL) was treated with DIPEA (229 µL, 1.3 mmol, 6.0 equiv.) and TSTU (92.1 mg, 306 µmol, 1.4 equiv.). The mixture was stirred for 10 min at room temperature. In a separate vial a solution of CA-NHBoc (78.7 mg, 241 µmol, 1.1 eq.) in TFA (500 µL) was shaken for 20 min. The solution was evaporated and dried. The residue was taken up in DMSO (500 µL) and added to the other mixture together with DIPEA (76 µL, 438 mmol, 2.0 equiv.). The mixture was shaken for 30 min and then acidified with TFA (150 µL). RP-HPLC (50 mL min$^{-1}$, 10% to 90% B in 60 min) gave Cyanine3-CA (62 mg, 43%) as a red solid.

HRMS (ESI): calc. for $C_{40}H_{57}ClN_3O_3^+$ $[M]^+$ : 662.4083; found 661.4084.

**Cyanine5-CA**

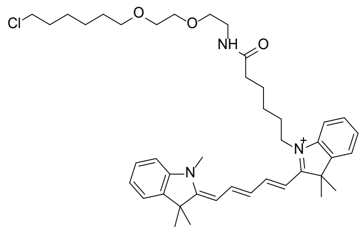

Cyanine5-CA

Cyanine5-COOH was synthesized according to Ueno et al. 2010.[27]

A solution of Cyanine5-COOH (100 mg, 207 µmol, 1.0 equiv.) in dry DMSO (2.0 mL) was treated with DIPEA (205 µL, 1.24 mmol, 6.0 equiv.) and TSTU (87.1 mg, 289 µmol, 1.4 equiv.). The mixture was stirred for 10 min at room temperature. In a separate vial a solution of CA-NHBoc (73.5 mg, 227 µmol, 1.1 eq.) in TFA (500 µL) was shaken for 20 min. The solution was evaporated and dried. The residue was taken up in DMSO (500 µL) and added to the other mixture together with DIPEA (68 µL, 414 mmol, 2.0 equiv.). The mixture was shaken for 30 min and then acidified with TFA (150 µL). RP-HPLC (50 mL min$^{-1}$, 10% to 90% B in 60 min) gave Cyanine5-CA (98 mg, 69%) as a blue solid.

HRMS (ESI): calc. for $C_{42}H_{59}ClN_3O_3^+$ $[M]^+$ : 688.4239; found 688.4239.

**Alexa647-CA**

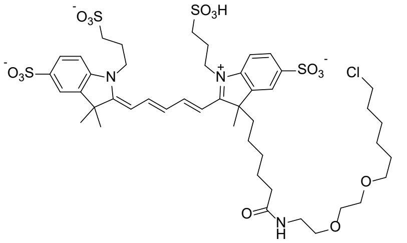

A solution of Alex647-NHS (0.3 mg, 0.3 µmol, 1.0 equiv.) in dry DMSO (150 µL) was treated with DIPEA (0.12 µL, 0.9 µmole, 6.0 equiv.). The mixture was stirred for 10 min at room temperature. In a separate vial a solution of CA-NHBoc (0.38 mg, 1.1 µmol, 3.6 eq.) in TFA (50 µL) was shaken for 20 min. The solution was evaporated and dried. The residue was taken up in DMSO (50 µL) and added to the other mixture together with DIPEA (68 µL, 414 mmol, 2.0 equiv.). The mixture was shaken for 30 min and then acidified with TFA (150 µL). RP-HPLC (8 mL $min^{-1}$, 5% to 60% B in 32 min) gave Alexa647-CA as a blue solid, which was taken up in 40 µL DMSO (1.98 mM, 25%).
HRMS (ESI): calc. for $C_{46}H_{64}ClN_3O_{15}S_4^{2-}$ $[M]^+$ : 530.6460; found 530.6449.

**Plasmids:** A pET51b(+) vector (Novagen) was used for protein production in *Escherichia coli*. Proteins were N-terminally tagged with $His_{x10}$, followed by a TEV cleavage site (ENLYFQ|G). For *in vitro* screening EGFP was fused to the C terminus of HaloTag7. A pcDNA5/FRT/TO or a pcDNA5/FRT vector (ThermoFisher Scientific) was used for transient expression as well as stable cell line establishment in mammalian cells. HaloTag7 and its variants were N-terminally fused to T2A-EGFP for expression in the cytosol. HaloTag7 and its variants were fused C-terminally to CEP41, H2B, and TOMM20 for expression in mammalian cells. In addition, T2A-EGFP was fused C-terminally of HaloTag7 and its variants. The Fucci sensors were constructed as follows: Fucci(SA): HaloTag7-Geminin(1-110)-P2A-HaloTag9-Cdt(30-120), Fucci(SCA): HaloTag7-Geminin(1-110)-P2A-HaloTag9-Cdt(1-100)Cy+, Fucci(CA): HaloTag7-Geminin(1-110)-P2A-HaloTag9-Cdt(1-100)Cy-[28], two different codon optimizations were used for HaloTag7 and HaloTag9, the later was obtained from gene synthesis (Eurofins). Cloning was performed by Gibson assembly[29]. DNA was subsequently electroporated in *E. cloni*10G (Lucigene) and plated on agar Lysogenic Broth plates with 100 µg/mL ampicillin and incubated at 37 °C overnight. EGFP, T2A-EGFP (Addgene plasmid # 135443)[23], CEP41 (Addgene plasmid # 135446)[23], TOMM20 (Addgene plasmid # 135443)[23], H2B (Addgene plasmid # 135444)[23], hGeminin(1-110) and hCdt(30-130): pLL3.7m-Clover-Geminin(1-110)-IRES-mKO2-Cdt(30-120) was a gift from Michael Lin (Addgene plasmid # 83841)[24], and hCdt(1-100): pCSII-EF-miRFP709-hCdt(1-100) was a gift from Vladislav Verkhusha (Addgene plasmid # 80007)[25] were used as template plasmids. pCAGGS-Raichu-RhoA-CR a gift from Michael Lin (Addgenen plasmid # 40258)[26] was directly used for transient transfection. For more information see Supplementary Table S14. Plasmids generated herein have been deposited on Addgene with accession codes listed in Supplementary Table S13.

**Protein production and purification:** Proteins were expressed in the *E. coli* strain BL21(DE3)-pLysS. Lysogen broth (LB) cultures were grown at 37 °C to an optical density at 600 nm ($OD_{600nm}$) of 0.8, induced by the addition of 0.5 mM isopropyl-β-D-thiogalactopyranoside (IPTG) and grown at 17 °C overnight in the presence of 1 mM $MgCl_2$. The cells were harvested by centrifugation (4,500 g, 10 min, 4 °C) and lysed by sonication (5 min, cycle 5, 70%, SonoPlus Bandelin). The cell lysate was cleared by centrifugation

(70,000 g, 20 min, 4 °C). Proteins were purified using affinity-tag Ni-NTA (Qiagen) leading to purity higher than 95% (verified by SDS-PAGE coomassie staining). Proteins were finally concentrated using an Ultra-0.5 mL centrifugal filter device (Amicon) with a molecular weight cut-off according to the protein size, followed by buffer exchange into activity buffer (<0.1 mM Imidazole). The proteins were stored in a glycerol 45% solution at −20 °C or flash frozen and stored at −80 °C. The protein's amino acid sequences are listed below.
For X-ray crystallography, proteins were produced as described above but purified using a HisTRAP FF affinity column (GE-Healthcare) on an ÄKTAPure M FPLC (GE-Healthcare). The proteins were concentrated using an Ultra-4 mL centrifugal filter device (Amicon) and were diluted to a final concentration of ~0.3 mg mL$^{-1}$ (ca. 40 mL) in TEV-cleavage buffer (25 mM $Na_2HPO_4$, 200 mM NaCl). β-mercaptoethanol (10 µL) and TEV protease (mass ratio substrate:TEV 30:1, TEV protease produced and purified in house by A. Bergner) were added and incubated at 30 °C overnight. The solution was filtered (0.22 µm) and the cleaved protein was harvested by reverse purification on a HisTRAP FF affinity column (GE-Healthcare), collecting the flow through. Proteins were concentrated using an Ultra-4 mL centrifugal filter device (Amicon) and further purified by size exclusion chromatography on a HiLoad 26/600 Superdex 75 pg column (GE-Healthcare) exchanging the buffer to activity buffer. Proteins were concentrated again and prepared to a final concentration of 5 µM in activity buffer. 3 mg of protein was incubated in presence of TMR-CA (10 µM) at room temperature overnight. The labeled protein was concentrated and an Illustra MicroSPin G-50 desalting column (GE-Healthcare) was employed to remove excess of unreacted fluorophore. The final protein concentration was adjusted to 13.0–15.0 mg mL$^{-1}$ using the absorbance at 280 nm, correcting the extinction coefficient of the protein by $\varepsilon_{280,\ TMR\text{-}CA}$ = 0.16.

**Protein sequences:**
>His-TEV-Halo-EGFP
MHHHHHHHHHHENLYFQGIGTGFPFDPHYVEVLGERMHYVDVGPRDGTPVLFLHGNPTSSYVWRNIIPHVAPTHRCI
APDLIGMGKSDKPDLGYFFDDHVRFMDAFIEALGLEEVVLVIHDWGSALGFHWAKRNPERVKGIAFMEFIRPIPTWDE
WPEFARETFQAFRTTDVGRKLIIDQNVFIEGTLPMGVVRPLTEVEMDHYREPFLNPVDREPLWRFPNELPIAGEPANIV
ALVEEYMDWLHQSPVPKLLFWGTPGVLIPPAEAARLAKSLPNCKAVDIGPGLNLLQEDNPDLIGSEIARWLSTLEIVSKG
EELFTGVVPILVELDGDVNGHKFSVSGEGEGDATYGKLTLKFICTTGKLPVPWPTLVTTLTYGVQCFSRYPDHMKQHDF
FKSAMPEGYVQERTIFFKDDGNYKTRAEVKFEGDTLVNRIELKGIDFKEDGNILGHKLEYNYNSHNVYIMADKQKNGIK
VNFKIRHNIEDGSVQLADHYQQNTPIGDGPVLLPDNHYLSTQSALSKDPNEKRDHMVLLEFVTAAGITLGMDELYK
Red: His-tag, Purple: TEV-cleavage site, Blue: HaloTag7, Black: Sites of mutations, Green: EGFP.

>His-TEV-Halo
MHHHHHHHHHHENLYFQGIGTGFPFDPHYVEVLGERMHYVDVGPRDGTPVLFLHGNPTSSYVWRNIIPHVAPTHRCI
APDLIGMGKSDKPDLGYFFDDHVRFMDAFIEALGLEEVVLVIHDWGSALGFHWAKRNPERVKGIAFMEFIRPIPTWDE
WPEFARETFQAFRTTDVGRKLIIDQNVFIEGTLPMGVVRPLTEVEMDHYREPFLNPVDREPLWRFPNELPIAGEPANIV
ALVEEYMDWLHQSPVPKLLFWGTPGVLIPPAEAARLAKSLPNCKAVDIGPGLNLLQEDNPDLIGSEIARWLSTLEI
Red: His-tag, Purple: TEV-cleavage site, Blue: HaloTag7, Black: Sites of mutations

**Protein crystallization:** Crystallization was performed at 20 °C using the vapor-diffusion method. HaloTag9 labeled with a TMR-CA fluorophore substrate, was concentrated to 13.0–15.0 mg mL$^{-1}$ in 50 mM

HEPES pH 7.3, 150 mM sodium chloride. Crystals of HaloTag9-TMR were grown by mixing equal volumes of protein solution and a reservoir solution containing 0.1 M MES pH 6.0, 1.0 M lithium chloride and 20% (m/v) PEG 6000. The crystals were briefly washed in cryoprotectant solution consisting of the reservoir solution with glycerol added to a final concentration of 20% (v/v), prior to flash-cooling in liquid nitrogen.

**X-ray diffraction data collection and structure determination:** Single crystal X-ray diffraction data was collected at 100 K on the X10SA beamline at the SLS (PSI, Villigen, Switzerland). Data was processed with XDS[30]. The structures of HaloTag9 labeled with TMR was determined by molecular replacement (MR) using Phaser[31] and HaloTag7-TMR coordinates (6Y7A) as a search model. Geometrical restraints for TMR were generated using Grade server[32]. The final model was optimized in iterative cycles of manual rebuilding using Coot[33] and refinement using Refmac5[34] and phenix.refine[35]. Data collection and refinement statistics are summarized in Supplementary Table S12, model quality was validated with MolProbity[36] as implemented in PHENIX.
Atomic coordinates and structure factors have been deposited in the Protein Data Bank under accession codes: 6ZVY (HaloTag7-Q156H-P174R-TMR).
Structural analysis was performed using PyMOL[37], phenix[35], and the APBS & PDB2PQR plug-in in Pymol using standard parameters (0.15 M ionic strength in monovalent salt, 310.0 K, protein dielectric of 2, and solvent dielectric of 78.0)[3].

**Polyacrylamide gel electrophoresis (PAGE):** HaloTag7 proteins (2 µM, 15 µL) were labeled using SiR-CA (10 µM) in activity buffer for 1 h at room temperature. After labeling, the proteins were separated by PAGE (4–20% 10 well Mini-Protean TGX, BioRad) as recommended by the manufacturer and revealed by in gel fluorescence using a ChemiDoc MD Imaging System (BioRad). SiR-CA labeled proteins were imaged using red epi illumination (695/55 nm), the proteins were revealed by coommassie staining (BioRad) and colorimetric imaging.

**Library generation:** The plasmid libraries consisting of site saturation mutagenesis performed on specific sites were prepared using degenerated primers according to Kille et al. (2014)[38]. The degenerated primers (Eurofins) were mixed in a ratio of NDT:VHG:TGG = 12:9:1 (N = any base, D = A, G or T , V = A, C or G, and H = A, C or T) and used for PCR amplification of two DNA fragments of pET51b(+)-His-tev-HaloTag7-EGFP. The saturation site belonged to an overlapping sequence between two DNA fragments. Plasmid libraries were prepared via Gibson assembly[29]. After electroporation in the *E. coli* strain *E. cloni* 10G (Lucigen), the library diversity was evaluated by serial dilution, plating on selective LB-agar plates (100 µg $mL^{-1}$ ampicillin) at 37 °C overnight and verification that more than 1,000 transformants were obtained by colony counting. Concomitantly, the library was isolated by plasmid extraction from a selective liquid LB culture (100 µg $mL^{-1}$ ampicillin) performed at 37 °C overnight (Qiagen kit). The plasmid libraries were sequenced by the Sanger method (Eurofins) to verify the proper incorporation of degenerate codons. Libraries were employed to transform *E. coli* strain BL21(DE3)-pLysS that were plated on selective LB agar plates (100 µg $mL^{-1}$ ampicillin) at 37 °C overnight. Single colonies were used to inoculate 400 µL selective LB medium (100 µg $mL^{-1}$ ampicillin) in a 96-deep well plate. Five wells were reserved for parental HaloTag7, five wells for CLIP-tag as a negative control and eight wells for sterility controls. The bacterial cultures were incubated at 37 °C overnight and 500 rpm. Then, 50 µL of the stationary phase cultures were employed to inoculate 950 µL selective LB medium (50 µg mL$^{-1}$ ampicillin) in a 96-deep well plate and incubated at 37 °C for 4 h at 500 rpm. The remaining culture was spun down (5,000 g, 15 min, 4 °C) and stored at 4 °C. Protein expression was induced by addition of 0.5 mM IPTG and grown at 17 °C overnight at 500 rpm. The cells were harvested by centrifugation (5,000 g, 15 min, 4 °C). The bacterial pellets were submitted to two cycles of freeze/thawing prior to resuspension in 300 µL lysis buffer (50 mM $K_2HPO_4$ pH = 8, 1 mg mL$^{-1}$ lysozyme, 2 mM $MgCl_2$, and 2.5 units mL$^{-1}$ benzonase (Turbo Nuclease, Jena Bioscience)) at 37 °C for 1 h. The cell lysate was cleared by centrifugation (5,000 g, 20 min, 4 °C). The cleared supernatant was transferred into non-binding black bottom 96-well plates for the screening assays.

**Screening assay:** Cell lysates (20 µL) were diluted in a non-binding black bottom 96-well plate into activity buffer (100 µL final, 0.5 mg mL$^{-1}$ BSA (Sigma)). Background SiR (620/20 ex, 680/30 em) and GFP (485/20 ex, 535/25 em) fluorescence intensity were measured prior to spiking SiR-CA (5 µL) in each well (5 nM, final SiR-CA concentration). After incubation at room temperature for 1 h, fluorescence intensities were measured again. An additional second labeling step was performed as previously described reaching 10 nM final SiR-CA concentration and intensities were again measured. The five control wells allowed to access the mean and s.d. GFP and SiR fluorescence intensities of the parental protein. Wells with GFP intensities lower than 10% of the control were discarded for the screening (too low expression). Wells with SiR fluorescence intensities 3- or 2-times s.d. larger than the control (first round: $mean_{par} \pm 3 \cdot s.d._{par}$, second and third round: $mean_{par} \pm 2 \cdot s.d._{par}$) were selected for further characterization. Plasmids of selected wells were obtained from stored bacterial cultures and sequenced. Selected variants for characterization were produced and purified from 50 mL selective LB cultures (as described above).
Each protein (1 µM) was labeled with SiR-CA in 100 µL activity buffer (containing 0.5 mg mL$^{-1}$ BSA (Sigma)) in a non-binding black bottom 96-well plate and incubated for 2 h at room temperature. The SiR and GFP fluorescence intensities were measured as previously described. Measurements were performed in triplicates, performing the independent labeling reactions in three separate wells. Mean and 90% confidence intervals were calculated for every variant and compared to the parental protein. Variants with significant changes in SiR fluorescence intensity compared to the parental protein were picked for further characterization (one-sided t test, $\alpha$ = 5%, DF = 4).

**Fluorescence increase characterization:** The most promising variants were subcloned into a pET51b(+) vector without the C-terminal EGFP fusion. Variant proteins were produced and purified from selective LB cultures (500 mL) as described above.
Fluorophores (Supplementary Table S3) were distributed into a non-binding black bottom 96-well plate (100 µL, 100 nM) and incubated at room temperature overnight. The next day 100 µL protein (2 µM, activity buffer containing 0.5 mg mL$^{-1}$ BSA (Sigma)) was added to the fluorophore and incubated for 4 h at room temperature. The respective fluorescence intensities were measured with a plate reader (TECAN Spark® 20M). The measurements were performed on four-time individually labeled protein (quadruplicates). Mean and 95% confidence intervals were calculated for every variant and compared to the parental protein (one sided t-test, $\alpha$ = 5%, DF = 6). Fluorescence excitation and emission spectra were measured using a plate reader (TECAN Spark® 20M).

**$D_{50}$ measurements:** The free acid of the fluorophore (5 µM) was diluted in 200 µL water-dioxane mixtures of 0%, 10%, 20%, 30%, 40%, 50%, 60%, 70%, 80%, 90%, and 100% in clear-bottom polypropylene 96-well plates (Greiner-bio one). Absorbance spectra were recorded on a plate reader (TECAN Spark® 20M) from 400 nm to 700 nm. Measurements were performed in triplicates. Data was baseline corrected and the absorbance values at $\lambda_{max}$ were plotted against the dielectric constants of water-dioxane mixtures[39]. The data was fitted with a sigmoidal curve (1) and the $D_{50}$ was determined as the point of inflection ($x_c$).

$$y(x) = \frac{a}{1+ e^{-k(x-x_c)}} \tag{1}$$

**Kinetics by plate reader:** Labeling kinetics of HaloTag7 variants (80 nM) were measured by time course fluorescence anisotropy measurements using TMR-CA (20 nM) in 200 µL activity buffer supplemented with 0.5 mg $mL^{-1}$ BSA at room temperature and on a plate reader (TECAN Spark® 20M) using the above stated filters for TMR and an injector system. G-factor and gain were calculated from three control measurements (buffer only, fluorophore in buffer, fully labeled protein in buffer). The data was fitted with a mono-exponential function (2) were $y_0$ corresponds to the y offset, $x_0$ to the x offset, $A$ to the amplitude and $\tau$ to the time constant. Using equation (3) the apparent second order rate constant ($k_{app}$) was calculated using the initial protein concentration $[P]_0$. Fitted parameters are reported as means from at least two measurements.

$$y(x) = y_0 + A \cdot e^{\frac{-(x-x_0)}{\tau}} \tag{2}$$

$$k_{app} = \frac{1}{\tau}[P]_0 \tag{3}$$

**Kinetics by Stopped-Flow:** Labeling kinetics of HaloTag7 and HaloTag9 with TMR-CA were measured by recording fluorescence anisotropy changes over time using a BioLogic SFM-400 stopped-flow instrument (BioLogic Science Instruments, Claix, France) in single mixing configuration at 37 °C. Monochromator wavelengths for excitation was set to 555 nm and a 570 nm long pass filter was used for detection. Protein and substrates were mixed in a 1:1 stoichiometry in activity buffer supplemented with 0.5 mg $mL^{-1}$ BSA. Concentrations were varied from 0.125 µM to 0.5 µM. The anisotropy of the free substrate was measured to obtain a baseline. The dead time of the instrument was measured according to the manufacturer's protocol (BioLogic Technical note #53) by recording the fluorescence decay during the pseudo first order reaction of *N*-acetyl-L-tryptophanamide with a large excess of *N*-bromosuccinimide and fitting the data to the first order reaction rate law. Recorded data was processed removing pre-trigger time points and averaging replicates. The data was fit to a two stage kinetic model (4, 5) using the DynaFit software[40]. Baseline anisotropy of the free fluorophore, substrate concentrations and dead time of the instrument were taken into account. Standard deviations (normal distribution verified) and confidence intervals of fitted parameters were estimated with the monte carlo method[41] with standard settings (N = 1000, 5% worst fits discarded). The derived parameters $K_D$ (dissociation constant) and $k_{app}$ (apparent second order rate constant) were calculated according to equation (6 and 7).

$$P + S \underset{k_{-1}}{\overset{k_1}{\leftrightarrow}} PS^* \tag{4}$$

$$PS^* \overset{k_2}{\rightarrow} PS \tag{5}$$

$$K_D = \frac{k_{-1}}{k_1} \tag{6}$$

$$k_{app} = k_1 \frac{k_2}{k_2 + k_{-1}} \tag{7}$$

**Thermostability:** Thermostability of His-tev-HaloTag7 or His-tev-HaloTag9 was measured at 0.5 mg $mL^{-1}$ in activity buffer on a nanoscale differential scanning fluorimeter Prometheus NT 48 (NanoTemper) over a temperature range from 20–95 °C with a heating rate of 1 °C $min^{-1}$ by monitoring changes in the ratio of the fluorescence intensities at 350 nm and 330 nm. The indicated melting temperature (mean±s.d., $N$ = 2 samples) corresponds to the point of inflection (maximum of the first derivative).

**Quantum yield:** Quantum yields were determined using a Hamamatsu Quantaurus QY. Fluorophores (0.5 µM) were directly added to the target protein (2.5 µM) in activity buffer. After incubation for 4 h at room temperature quantum yields were measured. Except for Cyanine3, where proteins (5 µM) were labeled with fluorophores (1 µM) in activity buffer for 12 h at room temperature and an Illustra MicroSPin G-50 desalting column (GE-Healthcare) was employed to remove excess of unreacted fluorophore.

**Extinction coefficient measurements:** Proteins (12 µM) were labeled with fluorophores (6 µM) in activity buffer for 3 h at room temperature. The absorbance spectra of labeled proteins (0.5 µM, 1.0 µM, 1.5 µM, 2.0 µM, and 3.0 µM) were recorded in clear bottom non-binding 96-well plates (200 µL) on a plate reader (TECAN Spark® 20M) from 400 nm to 700 nm. Data was baseline corrected and the maximum absorbance values were plotted against the concentration. The data was fitted to a linear function (8) and the extinction coefficients were calculated from the slope $b$.

$$y(x) = a + bx \tag{8}$$

**Computational chemistry:** Optimization of the quinoid and the spirolactone from of TMR were performed at the B3LYP/6-31G(d,p) level of theory using the software package Gaussian 16[42]. Solvent effects were modeled using the polarizable continuum model SMD. Molecules were visualized using the Avogadro software[43].

Molecular modeling of TMR on the surface of HaloTag7 or HaloTag9 was performed using MacroModel[44] and the Protein Preparation Wizard[45] both part of Maestro[46] (Schrödinger Software). Relative energies were calculated by molecular modeling using the force field OPLS3e[47] in water constraining the protein as well as the remainder of the ligand apart from $NMe_2$.

**Cell culture and transfection:** U-2 OS (ATCC) and U-2 OS Flp-In TREx Cep41-HaloTag7[23] cells were cultured in high-glucose phenol red free DMEM (Life Technologies) medium supplemented with GlutaMAX (Life Technologies), sodium pyruvate (Life Technologies) and 10% FBS (Life Technologies) in a humidified 5% $CO_2$ incubator at 37 °C. Cells were split every 3–4 days or at confluency. Cell lines were regularly tested for mycoplasma contamination. Cells were seeded on 8 well glass bottom dishes (Ibidi) three to one day before imaging. Transient transfections were performed using Lipofectamine™ 2000 reagent (Life Technologies) according to the manufacturer's recommendations: the DNA (0.3 µg) was mixed with OptiMEM I (10 µL, Life Technologies) and Lipofectamine™ 2000 (0.75 µL) was mixed with OptiMEM I (10 µL). The solutions were incubated for 5 min at room temperature, then mixed and incubated for an additional 20 min at room temperature. The prepared DNA-Lipofectamine complex was added to one of the wells in an 8 well glass bottom dish with cells at 50–70% confluency. After 12 h incubation in a humidified 5% $CO_2$ incubator at 37 °C the medium was changed to fresh medium. The cells were incubated under the same conditions for 24–48 h before imaging.

**Stable cell line establishment:** The Flp-In™ T-REx™ System (ThermoFisher Scientific) was used to generate stable cell lines exhibiting tetracycline-inducible expression of the gene of interest (GOI). Briefly, pcDNA5/FRT/TO-GOI or pcDNA5/FRT-GOI and pOG44 were co-transfected into the host cell line U-2 OS FlpIn TREx[48]. Homologous recombination between the FRT sites in pcDNA5/FRT/TO-GOI and the host cell chromosome, catalyzed by the Flp recombinase expressed from pOG44, produced the U-2 OS FlpIn TREx cells expressing stable and inducible the GOI. Selection was performed using 100 µg $mL^{-1}$ hygromycin B (ThermoFisher Scientific) and 15 µg $mL^{-1}$ blasticidine (ThermoFisher Scientific). Stable cell lines were seeded on glass bottom dishes as described in the previous section, and induced if necessary using 100 µg $mL^{-1}$ doxycycline (Sigma Aldrich) for 24–48 h prior to imaging. The following cell lines were established and used for further experiments: HaloTag7-T2A-EGFP, HaloTag9-T2A-EGFP (cytosolic expression); H2B-HaloTag7, Tomm20-HaloTag7, Cep41-HaloTag9, H2B-HaloTag9, Tomm20-HaloTag9, Fucci(SA) (non-inducible): HaloTag7-Geminin(1-110)-P2A-HaloTag9-Cdt(30-120), Fucci(SCA) (non-inducible): HaloTag7-Geminin(1-110)-P2A-HaloTag9-Cdt(1-100)Cy+, and Fucci(CA) (non-inducible): HaloTag7-Geminin(1-110)-P2A-HaloTag9-Cdt(1-100)Cy-. For more information see Supplementary Table S14.

**Fixation:** U-2 OS cells stably expressing Tomm20-HaloTag9 were seeded three days prior to fixation. They were transfected with H2B-HaloTag7 as described above two days prior fixation. Fixation was performed as follows: cells were prefixed in 2.4% [w/v] formaldehyde (PFA) in PBS for 45 s, permeabilized in 0.4% [v/v] Triton X-100 in PBS for 3 min and fixed in 2.4% [w/v] PFA in PBS for 30 min. PFA was quenched by 100 mM $NH_4Cl$ in PBS for 5 min. After washing three times for 5 min in PBS, the cells were labeled as described below.

**Labeling and sample preparation:** Cells were labeled with the respective fluorophores (Fluorophore-CA 1–2 µM, 1–3 h, 37 °C; or MaP618-Actin (Spirochrome), MaP555-Actin (Spirochorme) 2 µM, 3 h, 37 °C; or MaP555-DNA (Spirochrome), MaP555-Tubulin (Spirochorme) 1 µM, 3 h, 37 °C) in phenol-red free DMEM medium supplemented with GlutaMAX, sodium pyruvate, and 10% FBS (all Life Technologies), washed with the same medium (twice for 1 min, 37 °C) except for Actin, Tubulin, DNA, and Fucci for which labeling and imaging were performed in the same medium.

**Confocal microscopy:** Confocal fluorescence microscopy was performed on a Leica SP8 FALCON microscope (Leica Microsystems) equipped with a Leica TCS SP8 X scanhead; a SuperK white light laser, Leica HyD SMD detectors, a HC PL APO CS2 20x0.75 dry objective, a HC PL APO CS2 40x1.10 water objective

and a water immersion micro dispenser. Emission was collected as indicated in Supplementary Table S15. The microscope was equipped with a $CO_2$ and temperature controllable incubator (Life Imaging Services, 37 °C).
For brightness comparison, stable cell lines expressing HaloTag variants in the cytosol were seeded and labeled as described above. Cells were focused in the GFP channel and z-stacks were recorded with 1 µm step size over 22 µm. The summed stacks were analyzed as follows: the mean intensity of a rectangular ROI within the cell was normalized by the GFP intensity using a custom written Fiji macro[49,50].
Images of Cep41 were acquired transiently transfecting U-2 OS cells with Cep41-HaloTag7-T2A-EGFP or Cep41-HaloTag9-T2A-EGFP and labeling them as described above. Cells were focused in the GFP channel and z-stacks were recorded. Cep41-Halo images were background corrected, rescaled to the expression levels using the EGFP intensity values from within a ROI over the entire cell area and depicted using the same brightness and contrast settings.
Photostability measurements were performed using a PMT detector to collect emission. Stable cell lines expressing HaloTag7 and HaloTag9 as H2B fusions were seeded and labeled as described above. Cells were focused and a z-stack was recorded with 2 µm step size over 22 µm, using a pinhole of 5 Airy Unit (AU) and 2% (630 nm, SiR), 1.5% or 5% ( 615 nm, CPY, MaP618 or $JF_{614}$) and 2% (555 nm, TMR, MaP555) laser intensity. This was followed by the acquisition of 8 consecutive photobleaching frames in the focal plane at 100% laser intensity. Z-stack and photobleaching was repeated 60-times. The summed stacks were analyzed as follows: the mean intensity of ROIs around the nuclei were normalized to the mean intensity found at $t_0$.
Fluorescence excitation and emission spectra of MaP555-CA, MaP555-Actin, MaP555-Tubulin, MaP555-DNA, MaP618-CA, and MaP618-Actin were measured in live U-2 OS cells expressing either HaloTag7 or HaloTag9 in the cytosol or blank U-2 OS cells. MaP555 excitation: exciting at 475–575 nm in 2 nm steps collecting at 595–700 nm. MaP555 emission: exciting at 520 nm and collecting at 530–627 nm in 3 nm steps with a bandwidth of 10 nm. MaP618 excitation: exciting at 550–650 nm in 2 nm steps collecting at 670–780 nm. MaP618 emission: exciting at 600 nm and collecting at 610–707 nm in 3 nm steps with a bandwidth of 10 nm.

**FCS measurements:** FCS was performed on a Leica SP8 FALCON microscope (as described above) using a HC PL APO CS2 40x1.10 water objective with a motorized correction collar. FCS traces (30 s) were measured in U-2 OS cells expressing either HaloTag7 or HaloTag9 in the cytosol (no induction). The cells were labeled with MaP618-CA (150 nM, 2 h) and washed twice for 1 min. Excitation and emission collection was performed as indicated in Supplementary Table S15. Five traces per cell and a total of 36 cells from three biological replicates were measured. Data analysis (correlation and fitting) was performed using the LAS-X software (Leica Microsystems) fitting a free 3D diffusion model including a triplet component[51]. The diffusion amplitude ($G(0)$) as well as the mean photon counts (PC) over the 30 s trace were used to calculate the molecular brightness (*mB*, Equation 9).

$$mB = G(0) * \mathrm{PC} \quad (9)$$

**STED microscopy:** Imaging was performed on an Abberior easy3D STED/RESOLFT QUAD scanning microscope (Abberior Instruments GmbH, Göttingen, Germany) built on a motorized inverted microscope

IX83 (Olympus, Tokyo, Japan). The microscope was equipped with a pulsed STED lasers at 775 nm, and with a 640 nm excitation laser. Spectral detection was performed with avalanche photodiodes (APD) in the following spectral window: 650-725 nm. Images were acquired with a 100x/1.40 UPlanSApo Oil immersion objective lens (Olympus). Pixel size was 25 nm for all images. Laser powers and dwell times were kept constant so HaloTag9 and HaloTag7 images were comparable. For photobleaching measurements 31 consecutive STED frames were acquired and the intensity within a rectangular ROI was compared over time. Movement of mitochondria out or into the ROI was neglected.

**Fluorescence lifetime imaging microscopy:** FLIM was performed on a Leica SP8 FALCON microscope (as described above) at a pulse frequency of 80 MHz unless otherwise stated. Emission was collected as indicated in Supplementary Table S15.

For determination of fluorescence lifetimes cells stably expressing HaloTag7 or HaloTag9 in the cytosol were imaged, collecting 1,000 photons per pixel. The acquired images of cells were thresholded to remove background signal from empty coverslip space. Mean fluorescence lifetimes were calculated in the LAS X software (Leica Microsystems) by fitting a mono-exponential decay model (n-exponential reconvolution) to the decay ($\chi^2 < 1.2$).

Structural images (species separation) were acquired as indicated in Supplementary Table S15 and species separation was performed via phasor analysis (Leica Microsystems)[52–54]. Images in Supplementary Fig. S16 were analyzed using Pattern Matching in SymPhoTime64 (PicoQuant).

Long-term cell cycle measurements were performed using the Navigator function of the SP8. Cells were labeled with MaP618-CA (1 µM, no wash) and imaged after 1 h. FLIM images were acquired every 12 min and an autofocus z-stack measurement was performed before every image. Water immersion was controlled using the water immersion micro dispenser. Biosensor multiplexing was performed on the same set-up, acquiring FLIM images in both channels every 5 min for 2-4 h. Images were analyzed in LAS X (Leica Microsystems) applying pixel binning (2). FastFLIM images (average photon arrival times per pixel) are shown. Additionally, phasor analysis was used to evaluate the different cell populations.

**TauContrast microscopy:** TauContrast microscopy was performed on a STELLARIS 8 FALCON microscope (Leica Microsystems, FALCON system only used for comparison). The microscope was equipped with a White Light Laser with tunable excitation wavelengths 440-790 nm operating at 80 MHz. Spectral detection was performed with Power HyD X photon-counting detectors in the following spectral window: 630-700 nm. Images were acquired with a 86x/1.20 STED WHITE water immersion objective lens (Leica Microsystems). Pixel size was 176 nm for all images. TauContrast images were analyzed using the LAS X software. For comparison FastFLIM images were simultaneously acquired using the FALCON system.

**STED-FLIM microscopy:** Line sequential, confocal/STED-FLIM imaging was performed on a STELLARIS 8 STED FALCON microscope (Leica Microsystems). The microscope was equipped with a White Light Laser with tunable excitation wavelengths 440-790 nm operating at 80 MHz, and a pulsed STED laser at 775 nm operating at 80 MHz. Spectral detection was performed with Power HyD X photon-counting detectors in the following spectral window: 630-760 nm. Images were acquired with a 86x/1.20 STED WHITE water immersion objective lens (Leica Microsystems). Pixel size was 20 nm for all images. Images were analyzed using species separation via phasor analysis available in FALCON through the LAS X software.

**Software and image processing:** Statistical analysis as well as curve fitting was performed using OriginLab[55] or R[56] including packages readxl,[57] fBasics,[58] tidyverse,[59] gghighlight,[60] ggplot2,[61] ggrepel,[62] ggpubr,[63] broom.[64] All images were processed with ImageJ/Fiji[49,50] and macros written therein unless otherwise stated.

### References

1. Deprey, K. & Kritzer, J. A. HaloTag Forms an Intramolecular Disulfide. *Bioconjug. Chem.* (2021). doi:10.1021/acs.bioconjchem.1c00113
2. Berro, A. J. & Schreiter, E. R. Crystal structure of HaloTag bound to tetramethylrhodamine-HaloTag ligand. (2019). doi:10.2210/pdb6u32/pdb
3. Baker, N. A., Sept, D., Joseph, S., Holst, M. J. & McCammon, J. A. Electrostatics of nanosystems: Application to microtubules and the ribosome. *Proc. Natl. Acad. Sci. U. S. A.* **98**, 10037–10041 (2001).
4. Roberti, M. J. *et al.* TauSense: a fluorescence lifetime-based tool set for everyday imaging. *Nat. Methods* (2020).
5. Lambert, T. J. FPbase: a community-editable fluorescent protein database. *Nat. Methods* **16**, 277–278 (2019).
6. Shirmanova, M. V *et al.* FUCCI - Red : a single - color cell cycle indicator for fluorescence lifetime imaging. *Cell. Mol. Life Sci.* (2021). doi:10.1007/s00018-020-03712-7
7. Yoshizaki, H. *et al.* Activity of Rho-family GTPases during cell division as visualized with FRET-based probes. *J. Cell Biol.* **162**, 223–232 (2003).
8. Los, G. V *et al.* HaloTag: A Novel Protein Labeling Technology for Cell Imaging and Protein Analysis. *ACS Chem. Biol.* **3**, 373–382 (2008).
9. Grimm, J. B. *et al.* A general method to fine-tune fluorophores for live-cell and in vivo imaging. *Nat. Methods* **14**, 987–994 (2017).
10. Grimm, J. B. *et al.* A general method to improve fluorophores for live-cell and single-molecule microscopy. *Nat. Methods* **12**, 244–250 (2015).
11. Grimm, J. B. *et al.* Deuteration improves small-molecule fluorophores. *bioRxiv* (2020). doi:10.1101/2020.08.17.250027
12. Hoffman, D. P. *et al.* Correlative three-dimensional super-resolution and block-face electron microscopy of whole vitreously frozen cells. *Science* **367**, eaaz5357 (2020).
13. Grimm, J. B., Brown, T. A., Tkachuk, A. N. & Lavis, L. D. General Synthetic Method for Si-Fluoresceins and Si-Rhodamines. *ACS Cent. Sci.* **3**, 975–985 (2017).
14. Grimm, J. B. *et al.* A general method to optimize and functionalize red-shifted rhodamine dyes. *Nat. Methods* **17**, 815–821 (2020).
15. Wang, L. *et al.* A general strategy to develop cell permeable and fluorogenic probes for multicolour nanoscopy. *Nat. Chem.* **12**, 165–172 (2020).
16. Grimm, J. B., Gruber, T. D., Ortiz, G., Brown, T. A. & Lavis, L. D. Virginia Orange: A Versatile, Red-Shifted Fluorescein Scaffold for Single- And Dual-Input Fluorogenic Probes. *Bioconjug. Chem.* **27**, 474–480 (2016).
17. Butkevich, A. N. *et al.* Fluorescent Rhodamines and Fluorogenic Carbopyronines for Super-Resolution STED Microscopy in Living Cells. *Angew. Chem. Int. Ed.* **55**, 3290–3294 (2016).

18. Deo, C. *et al.* The HaloTag as a general scaffold for far-red tunable chemigenetic indicators. *Nat. Chem. Biol.* (2021). doi:10.1038/s41589-021-00775-w
19. Jonker, C. T. H. *et al.* Accurate measurement of fast endocytic recycling kinetics in real time. *J. Cell Sci.* **133**, jcs231225 (2020).
20. Lukinavičius, G. *et al.* A near-infrared fluorophore for live-cell super-resolution microscopy of cellular proteins. *Nat. Chem.* **5**, 132–139 (2013).
21. Zheng, Q. *et al.* Rational Design of Fluorogenic and Spontaneously Blinking Labels for Super-Resolution Imaging. *ACS Cent. Sci.* **5**, 1602–1613 (2019).
22. Liu, Z. *et al.* Systematic comparison of 2A peptides for cloning multi-genes in a polycistronic vector. *Sci. Rep.* **7**, 2193 (2017).
23. Frei, M. S. *et al.* Photoactivation of silicon rhodamines via a light-induced protonation. *Nat. Commun.* **10**, 4580 (2019).
24. Bajar, B. T. *et al.* Fluorescent indicators for simultaneous reporting of all four cell cycle phases. *Nat. Methods* **13**, 993–996 (2016).
25. Shcherbakova, D. M. *et al.* Bright monomeric near-infrared fluorescent proteins as tags and biosensors for multiscale imaging. *Nat. Commun.* **7**, 12405 (2016).
26. Lam, A. J. *et al.* Improving FRET dynamic range with bright green and red fluorescent proteins. *Nat. Methods* **9**, 1005–1012 (2012).
27. Ueno, Y. *et al.* Encapsulated energy-transfer cassettes with extremely well resolved fluorescent outputs. *J. Am. Chem. Soc.* **133**, 51–55 (2011).
28. Sakaue-Sawano, A. *et al.* Genetically Encoded Tools for Optical Dissection of the Mammalian Cell Cycle. *Mol. Cell* **68**, 626–640 (2017).
29. Gibson, D. G. *et al.* Enzymatic assembly of DNA molecules up to several hundred kilobases. *Nat. Methods* **6**, 343–345 (2009).
30. Kabsch, W. XDS. *Acta Crystallogr. Sect. D Biol. Crystallogr.* **66**, 125–132 (2010).
31. McCoy, A. J. *et al.* Phaser crystallographic software. *J. Appl. Crystallogr.* **40**, 658–674 (2007).
32. Smart, O. S. *et al.* Grade Web Server. (2011).
33. Emsley, P., Lohkamp, B., Scott, W. G. & Cowtan, K. Features and development of Coot. *Acta Crystallogr. Sect. D Biol. Crystallogr.* **66**, 486–501 (2010).
34. Murshudov, G. N. *et al.* REFMAC5 for the refinement of macromolecular crystal structures. *Acta Crystallogr. Sect. D Biol. Crystallogr.* **67**, 355–367 (2011).
35. Adams, P. D. *et al.* PHENIX: A comprehensive Python-based system for macromolecular structure solution. *Acta Crystallogr. Sect. D Biol. Crystallogr.* **66**, 213–221 (2010).
36. Chen, V. B. *et al.* MolProbity: All-atom structure validation for macromolecular crystallography. *Acta Crystallogr. Sect. D Biol. Crystallogr.* **66**, 12–21 (2010).
37. Schrödinger, L. The PyMOL Molecular Graphics System, Version 2.1.1. (2015).
38. Kille, S. *et al.* Reducing Codon Redundancy and Screening Effort of Combinatorial Protein Libraries Created by Saturation Mutagenesis. *ACS Synth. Biol.* **2**, 83–92 (2013).
39. Åkerlöf, G. & Short, O. A. The Dielectric Constant of Dioxane-Water Mixtures between 0 and 80°. *J. Am. Chem. Soc.* **58**, 1241–1243 (1936).
40. Kuzmic, P. Program DYNAFIT for the Analysis of Enzyme Kinetic Data : Application to HIV Proteinase. *Anal. Biochem.* **273**, 260–273 (1996).
41. Straume, M. & Johnson, M. L. B. T.-M. in E. Monte Carlo Method for determining

complete confidence probability distributions of estimated model parameters. in *Numerical Computer Methods* **210**, 117–129 (Academic Press, 1992).
42. Frisch, M. J. *et al.* Gaussian 16 Rev. B.01. (2016).
43. Hanwell, M. D. *et al.* Avogadro: an advanced semantic chemical editor, visualization, and analysis platform. *J. Cheminform.* **4**, 17 (2012).
44. Schrödinger Release 2020-3: MacroModel, Schrödinger, LLC, New York, NY. (2020).
45. Madhavi Sastry, G., Adzhigirey, M., Day, T., Annabhimoju, R. & Sherman, W. Protein and ligand preparation: parameters, protocols, and influence on virtual screening enrichments. *J. Comput. Aided. Mol. Des.* **27**, 221–234 (2013).
46. Schrödinger Release 2020-3: Maestro, Schrödinger, LLC, New York, NY. (2020).
47. Harder, E. *et al.* OPLS3: A Force Field Providing Broad Coverage of Drug-like Small Molecules and Proteins. *J. Chem. Theory Comput.* **12**, 281–296 (2016).
48. Malecki, M. J. *et al.* Leukemia-Associated Mutations within the NOTCH1 Heterodimerization Domain Fall into at Least Two Distinct Mechanistic Classes. *Mol. Cell. Biol.* **26**, 4642–4651 (2006).
49. Rueden, C. T. *et al.* ImageJ2: ImageJ for the next generation of scientific image data. *BMC Bioinformatics* **18**, 529 (2017).
50. Schindelin, J. *et al.* Fiji: An open-source platform for biological-image analysis. *Nat. Methods* **9**, 676–682 (2012).
51. Widengren, J., Mets, Ü. & Rigler, R. Fluorescence correlation spectroscopy of triplet states in solution: A theoretical and experimental study. *J. Phys. Chem.* **99**, 13368–13379 (1995).
52. Digman, M. A., Caiolfa, V. R., Zamai, M. & Gratton, E. The phasor approach to fluorescence lifetime imaging analysis. *Biophys. J.* **94**, L14–L16 (2008).
53. Digman, M. A. & Gratton, E. The phasor approach to fluorescence lifetime imaging: Exploiting phasor linear properties. in *Fluorescence Lifetime Spectroscopy and Imaging* (eds. Marcu, L., French, P. M. W. & Elson, D. S.) 235–248 (CRC Press, 2014).
54. Wang, P. *et al.* Complex Wavelet Filter Improves FLIM Phasors for Photon Starved Imaging Experiments. *Biomed. Opt. Express* in press (2021).
55. Origin(Pro), Version 2018b, OriginLab Corporation, Northampton, MA, USA.
56. R Core Team. R: A Language and Environment for Statistical Computing. (2019).
57. Wickham, H. & Bryan, J. readxl: Read Excel Files. (2019).
58. Wuertz, D., Setz, T. & Chalabi, Y. fBasics: Rmetrics - Markets and Basic Statistics. (2017).
59. Wickham, H. *et al.* Welcome to the tidyverse. *J. Open Source Softw.* **4**, 1686 (2019).
60. Yutani, H. gghighlight: Highlight Lines and Points in ‘ggplot2’. (2018).
61. Wickham, H. ggplot2: Elegant Graphics for Data Analysis. (2016).
62. Slowikowski, K. ggrepel: Automatically Position Non-Overlapping Text Labels with ‘ggplot2’. (2019).
63. Kassambara, A. ggpubr: ‘ggplot2’ Based Publication Ready Plots. (2019).
64. Robinson, D. & Hayes, A. broom: Convert Statistical Analysis Objects into Tidy Tibbles. (2019).
